# AdipoAtlas: A Reference Lipidome for Human White Adipose Tissue

**DOI:** 10.1101/2021.01.20.427444

**Authors:** Mike Lange, Georgia Angelidou, Zhixu Ni, Angela Criscuolo, Jürgen Schiller, Matthias Blüher, Maria Fedorova

**Affiliations:** Institute of Bioanalytical Chemistry, Faculty of Chemistry and Mineralogy, University of Leipzig, Germany; Center for Biotechnology and Biomedicine, University of Leipzig; Thermo Fisher Scientific, Dreieich, Germany; Institute of Medical Physics and Biophysics, Medical Faculty, University of Leipzig; Medical Department III (Endocrinology, Nephrology and Rheumatology), University of Leipzig, Leipzig, Germany; Helmholtz Institute for Metabolic, Obesity and Vascular Research (HI-MAG) of the Helmholtz Zentrum München at the University of Leipzig and University Hospital Leipzig, Leipzig, Germany

**Keywords:** human white adipose tissue, subcutaneous WAT, visceral WAT, obesity, lipidomics, LC-MS/MS, lipid identification, semi-absolute lipid quantification, ceramides, plasmalogens, triacylglycerols

## Abstract

Obesity, characterized by expansion and metabolic dysregulation of white adipose tissue (WAT), has reached pandemic proportions and acts as a primer for a wide range of metabolic disorders. Remodelling of WAT lipidome in obesity and associated comorbidities can explain disease etiology and provide valuable diagnostic and prognostic markers. To support understanding of WAT lipidome remodelling at the molecular level, we performed in-depth lipidomics profiling of human subcutaneous and visceral WAT of lean and obese individuals. Tissue-tailored preanalytical and analytical workflows allowed accurate identification and semi-absolute quantification of 1636 and 737 lipid molecular species, respectively, and summarized here in a form of human WAT reference lipidome. Deep lipidomic profiling allowed to identify main lipid (sub)classes undergoing depot/phenotype specific remodelling. Furthermore, previously unanticipated diversity of WAT ceramides was uncovered. AdipoAtlas reference lipidome will serve as a data-rich resource for the development of WAT-specific high-throughput methods and as a scaffold for systems medicine data integration.

## Introduction

The “industrial revolution” in modern omics technologies significantly enriched our understanding of human biology. Application of high-throughput transcriptomics and proteomics allowed to compile the Tissue Atlas within the Human Proteome Atlas project with expression levels of mRNA and proteins reported for 44 healthy human tissues, serving as a powerful resource for exploration of functional tissue specificities, future drug targets and potential biomarkers (Uhlén et al., 2015). Lipidomics, an omics branch aiming to identify and quantify individual lipid species, is not yet as advanced in the characterization of cell-, tissue-, and organ-specific lipid compositions. The majority of lipidomics studies aim for high-throughput screening of large sample cohorts and clinical translation (Huynh et al., 2019; Seah et al., 2020). Such analytical workflows, targeting robust applications, are optimized for bulk lipid extraction followed by a single analysis method and relative (disease vs control) quantification.

Considering the cooperative action of lipids in biological membranes and tight coregulation of anabolic and catabolic pathways of lipid metabolism, identification of tissue and cell type specific lipid signatures (reference lipidomes) is urgently required to facilitate deeper understanding of lipid biology in health and disease. Lipid cooperative actions are highly tissue/cell type specific at all levels of their functional activities including plasticity of cellular membranes, energy storage, redistribution, and coordinated signalling (Frayn et al., 2006; Furse et al., 2020). Furthermore, capturing alterations in lipid metabolism might be as important as identifying static lipid signatures resistant to certain (patho)physiological stimuli. Current advances in systems biology and medicine allow holistic integration of single and multiple omics levels (Alves et al., 2021). Several genome scale metabolic networks have been reconstructed, demonstrating high power in explaining human biology via integration of big omics datasets (Brunk et al., 2018; Noronha et al., 2019). Application of systems biology tools to lipid metabolism requires a detailed characterization of lipid molecular species both in a qualitative and quantitative manner. Availability of tissue-specific reference lipidomes would enable monitoring of the specificity of lipid metabolism and aid in understanding the cross-talk within and across different tissues.

Deep lipidome profiling cannot be performed in a high-throughput manner as it requires tissue specific optimization and application of several orthogonal analytical methods to ensure simultaneous coverage of lipid classes with different polarities, ionization properties and range of endogenous concentrations. By now, the best characterized composition is available for the blood plasma lipidome with around 600 lipid species described at lipid class and lipid molecular species levels (Bowden et al., 2017; Burla et al., 2018; Criscuolo et al., 2019). However, detailed quantitative inventory of peripheral tissue lipidomes are scarce. Currently a lot of scientific attention was attracted towards adipose tissue metabolism. Obesity, characterized by white adipose tissue (WAT) expansion and metabolic dysregulation, has reached pandemic proportions in modern societies with a prevalence of more than 20% of the population (Blüher, 2019). Obesity is associated with an increased threat of premature death due to the significantly higher risk of developing type 2 diabetes mellitus (T2DM), hypertension, coronary heart disease, stroke, and several types of cancer. Remodelling of WAT metabolism in obesity and, importantly, in development of metabolic complications is a cornerstone in understanding disease etiology. Adipose tissue is the main lipid storage organ characterized by its extraordinary capacity to store excess of nutrients in the form of triglycerides (TG), buffering this way the excess of free fatty acids (FFA) and preventing ectopic lipid accumulation in peripheral tissues. At conditions of chronic energy surplus WAT lipid metabolism undergoes significant remodelling to support oversupply of diet-derived fatty acids and carbohydrates, manifested in accumulation of TG in adipocyte lipid droplets, cellular hypertrophy and subsequent increase of WAT mass. So far WAT metabolism was studied from many different angels including genetic predisposition to obesity via genome wide association studies (Raulerson et al., 2019), changes in transcriptomics (Haffa et al., 2019), epigenetic (Martínez et al., 2014), and proteomics (Gómez-Serrano et al., 2016) patterns of WAT upon obesity development. However, studies reporting detailed quantitative description of depot specific (subcutaneous vs visceral) WAT lipidomes in lean and obese human individuals are limited.

Here, we present AdipoAtlas – a mass-spectrometry based reference lipidome of human WAT reporting over 1600 and 700 lipid species on qualitative and quantitative levels, respectively. To ensure optimal WAT lipidome coverage and recovery we carefully optimized each step of the preanalytical workflow from sample preparation over lipid extraction to fractionation. Subsequently, a combination of various LC-MS platforms coupled to curated software assisted and manual identification, and quantification workflows was applied to provide an inventory describing most biologically important lipid classes within adipose tissue. In order to display its utility, AdipoAtlas was used as a reference to illustrate the remodeling of WAT lipidome upon development of obesity in two different adipose tissue depots, subcutaneous and visceral. We show that ceramides containing the unusual sphingadienine base and TG containing polyunsaturated fatty acid (PUFA) residues are enriched in obese WAT. Moreover, we identified distinct responses of adipose tissue depots to increased metabolic demand by upregulation of depot-specific plasmalogen synthesis. *AdipoAtlas* presents tissue-specific quantitative lipidome, a data-rich resource freely available to all lipid researchers. AdipoAtlas will support further understanding of lipidomic alterations within human adipose tissue and can act as a guideline to generate other tissue specific lipidome maps.

## Results

### WAT-tailored lipid extraction and fractionation

WAT acts as the main lipid storage organ with TG present at exceedingly high concentrations masking other less abundant lipid classes (Figure 1A). For accurate molecular mapping of the WAT lipidome, both extraction and fractionation were optimized to ensure coverage of both highly abundant storage (TG) and low abundant membrane and signaling (phospholipids and sphingolipids, PL and SP) lipids. To this end, we created tissue pools of WAT from obese and lean individuals representing subcutaneous (SAT) and visceral (VAT) depots. Pooled samples were used to test three common extraction protocols (Folch, MTBE, and Hex/IPA) (Iverson et al., 2001; Matyash et al., 2008; Slatter et al., 2016). The most efficient extraction method was chosen based on the recovery of unpolar and polar lipid classes assessed by quantitative high-performance thin-layer-chromatography (qHPTLC) and ^31^P-NMR (Figure 1B and C; Supplementary Figure S1). Overall, the Folch two-phase extraction protocol was shown to be the most efficient in recovering both unpolar and polar lipids in human WAT.

**Figure 1:**
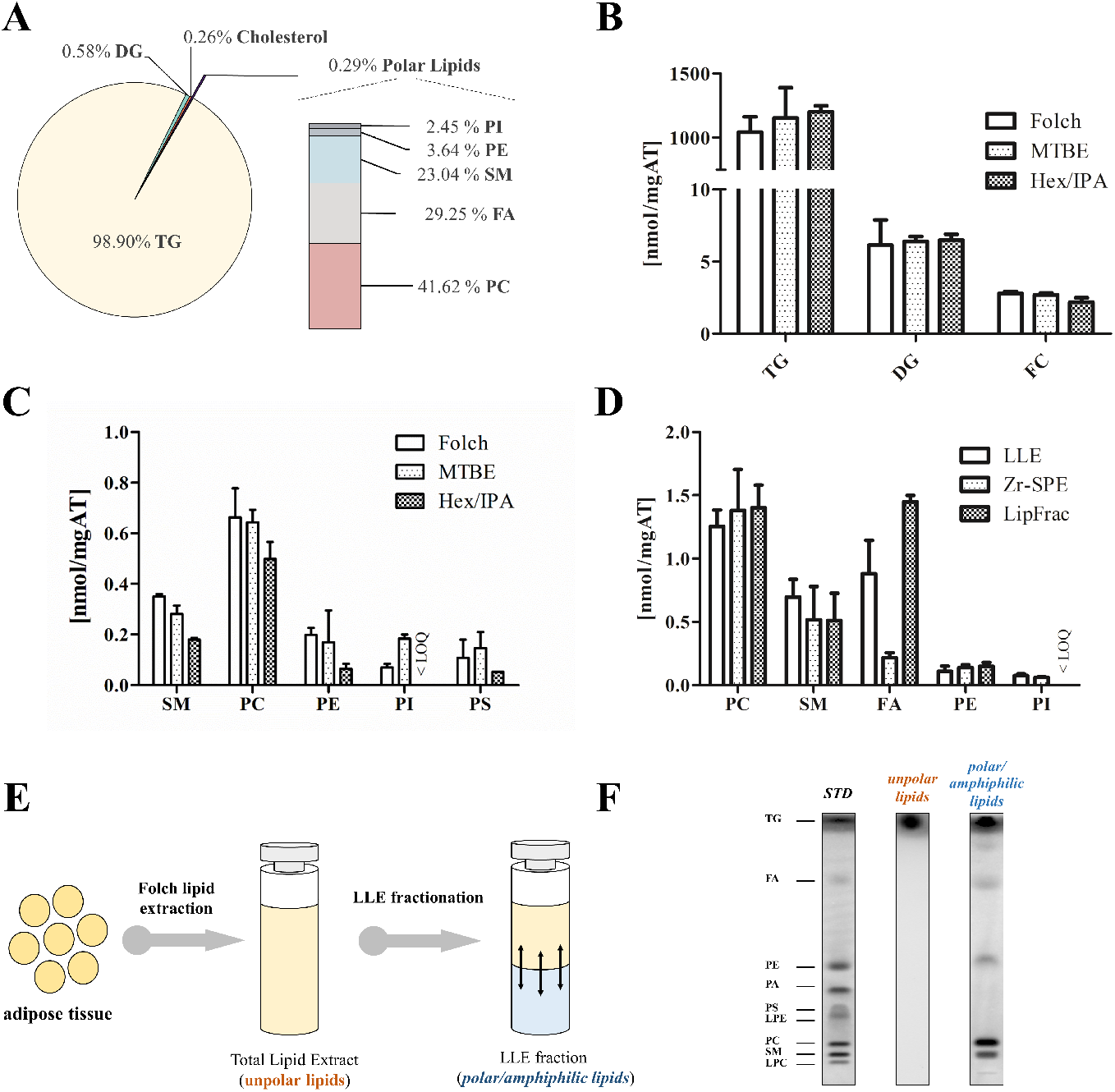
Optimization of sample preparation protocols for global lipidome profiling of human white adipose tissue (WAT). Pooled tissue samples from obese visceral and subcutaneous WAT were used for optimization purposes to reflect general abundance of lipid classes within human WAT. **A:** Lipid class specific WAT lipidome composition as determined by quantitative high-performance thin layer chromatography (qHPTLC) of Folch lipid extracts in combination with liquid/liquid extraction (LLE) for polar lipid enrichment. **B:** Extraction efficiency of unpolar lipids as determined by qHPTLC. WAT lipids were extracted by either the Folch, the methyl-*tert*-butyl-ether (MTBE) or the hexane/*i*-PrOH/HOAc (Hex/IPA) method. **C:** Extraction efficiency of phosphate containing polar lipids by the different lipid extraction protocols was determined with ^31^P-NMR. **D:** Efficiency to separate polar and unpolar lipids from WAT lipid extracts was compared using polarity separation by LLE, separation based on the presence of phosphate groups in lipids by zirconia-oxide based solid phase extraction (Zr-SPE) or aminopropyl SPE based lipid class fractionation (LipFrac). **E:** Schematic depiction of the optimized lipid extraction and fractionation protocol. **F:** qHPTLC analysis of WAT lipidome before (total lipid extract dominated by unpolar lipids) and after (enriched polar and amphiphilic lipids; ethanol fraction) LLE fractionation.

Even with the optimal extraction, polar lipids represented a minor fraction of WAT lipidome (Figure 1A and C). To facilitate deep lipidomic profiling, we performed fractionation of WAT lipid extracts. Lipid fractionation methods utilize differences in molecular motifs or polarity of lipid classes, and here we compared three orthogonal protocols based on (*ii*) hydrophobicity and ionization state (amino propyl solid phase extraction, SPE; hereafter LipFrac) (*ii*) presence of phosphodiester groups (Zr-SPE), and (*iii*) polarity (using liquid-liquid extraction, LLE). All three protocols showed similar recovery of PC, PE, and SM lipids (Figure 1D). LipFrac resulted in the highest recovery of free fatty acids but discriminated acidic PL (e.g., PI) due to their strong binding to the stationary phase. Zr-SPE effectively enriched phosphate group containing lipid classes but increased content of lysoPL and FFA due to PL alkaline hydrolysis during the elution step. LLE delivered sufficient enrichment efficiency for all polar lipid classes and was the most time efficient protocol (Figure 1D).

Here, we performed WAT specific optimization of lipid extraction and fractionation protocols, and identified the most efficient sample preparation strategy based on Folch lipid extraction followed by polarity based LLE allowing non-discriminative recovery and enrichment of lipids of different classes (Figure 1E and F). The workflow presented here utilized qHPTLC for bulk lipid quantification and can be easily adapted for optimization of extraction and enrichment protocols to any other tissue samples.

### WAT lipidome profiling

To increase biological meaningfulness of lipidomics data, lipid annotations should provide information on the lipid class as well as on the discrete fatty acyl chain composition rather than total number of carbon atoms and double bond equivalents. LC-MS/MS based lipidomics allows accurate identification of lipids at the molecular species level (e.g., PC 16:0_18:2), but in complex lipid mixtures with high dynamic range of lipid concentrations, the coverage of identified lipidome will depend on the total resolution of the analytical platform. For deep WAT lipidome profiling, we utilized three LC systems each coupled on-line to high resolution accurate mass (HRAM) tandem mass spectrometry (MS/MS) (Figure 2A). Highly hydrophobic TG lipids represented by a large diversity of molecular species and dynamic range of concentrations were separated using C30 reversed phase liquid chromatography (RPC), whereas the more polar LLE fraction (PLs, SLs, DGs) was resolved using C18 RPC (Figure 2B). Highly polar acyl carnitines (CAR) were separated by hydrophilic interaction liquid chromatography (HILIC) (Figure 2B), as they are not sufficiently retained on RP columns. Moreover, MS analysis using data-dependent acquisition (DDA) relied on different HRAM platforms and was specifically tailored to enhance identification depth for each targeted lipid class. To this end, C30 RPC separated TG were analyzed in positive polarity on a Q Exactive Plus mass spectrometer with traditional DDA, and on a Orbitrap Fusion Lumos Tribrid mass spectrometer using AcquireX deep scan acquisition workflow for in-depth identification of TG molecular species. Less abundant amphiphilic lipids on the other hand were detected by DDA on a Q Exactive platform and a Orbitrap Fusion Lumos instrument both in the positive and negative modes. As we originally failed to detect any cholesteryl esters (CE), retinol esters, desmosterol esters or cardiolipins in WAT total extracts we set up targeted parallel reaction monitoring (PRM) for the detection of most prominent species. That allowed us to detect 7 CE otherwise masked by highly abundant TGs. However, both retinol and desmosterol esters as well as cardiolipins remained undetected in human WAT. Overall, 111 LC-MS/MS analyses were performed to support lipid identification (Figure 2A and B, Material and Methods Figure 1).

**Figure 2:**
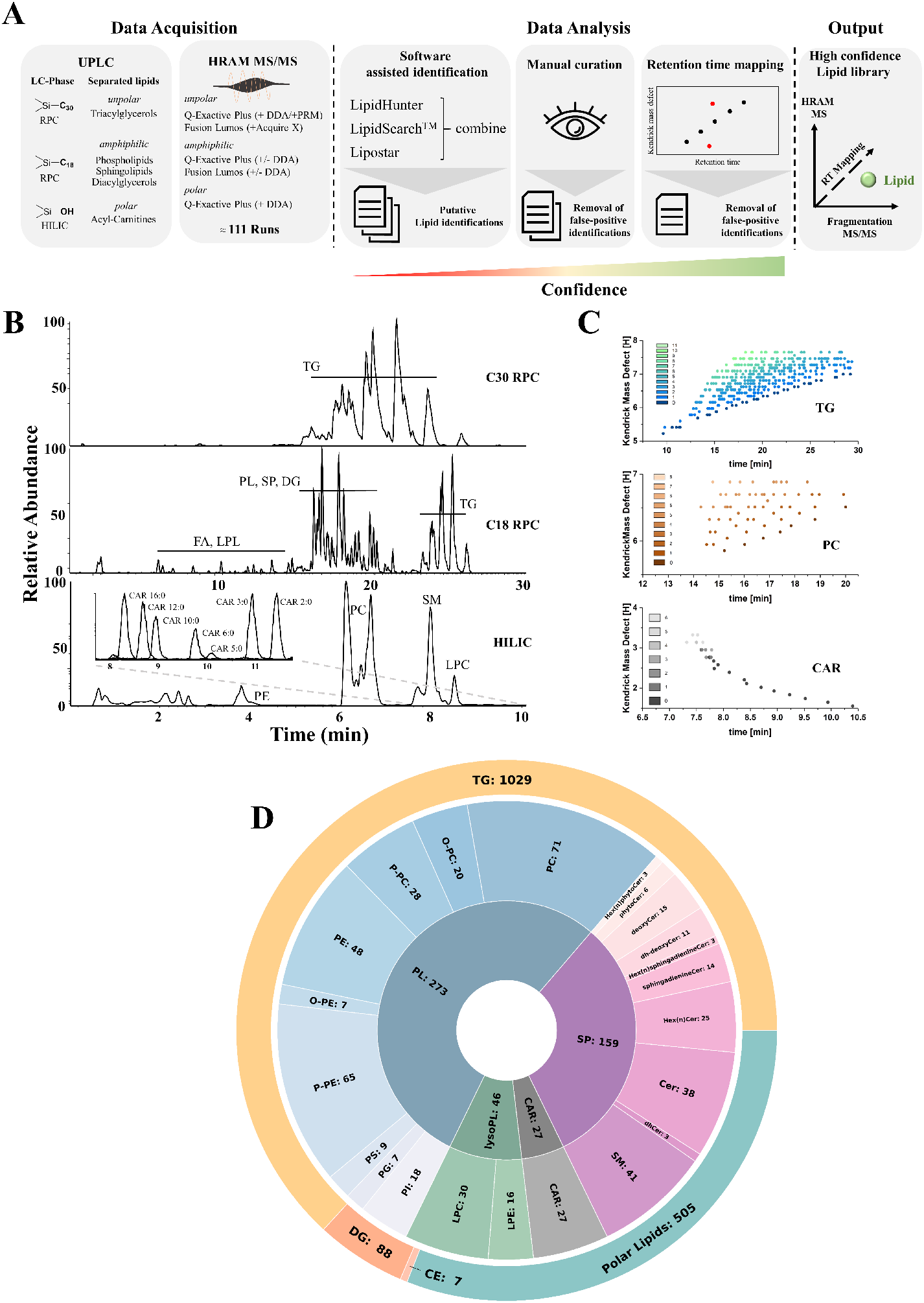
Workflow adapted for high confidence lipid identification from human white adipose tissue (WAT). **A:** Schematic depiction of the identification strategy for three dimensionally curated, high confidence lipid library of human WAT. Liquid chromatography (LC) and mass spectrometry (MS) platforms were tailored to allow for optimal separation and coverage of unpolar, amphiphilic and polar lipid classes. Data analysis featured three independent software tools followed by manual curation of lipid annotations. Putative lipid identifications were furthermore subjected to retention time mapping in order to increase identification confidence. **B:** Lipid class specific LC separation was applied to allow for the highest possible chromatographic resolution to achieve optimal lipidome coverage. LC-MS chromatograms of unpolar lipids separated by C30 reversed phase chromatography (RPC), amphiphilic lipids by C18 RPC and polar acylcarnitines (CAR) by hydrophilic interaction chromatography (HILIC). **C:** Exemplary depiction of retention time mapping for TG, PC and CAR lipid classes. Kendrick mass defect to the hydrogen base (KMD[H]) was plotted against lipid retention time to increase confidence of lipid identification. **D:** Graphical representation of human WAT lipid molecular species grouped by the corresponding lipid class obtained by high confidence identification strategy.

To ensure high-confidence accurate lipid identification, we used three software tools – LipidHunter (Ni et al., 2017), LipidSearch (Thermo Scientific, San Jose, CA), and Lipostar (Goracci et al., 2017). Obtained results were cross-matched, and the list of putative lipid identifications was manually curated to exclude false-positive identifications. We further validated manually curated lipid annotations by plotting the retention time of a given lipid species against its Kendrick mass defect to the hydrogen base (KMD) (Figure 2C). Detailed description of manual MS/MS curation and retention time mapping for lipids of different (sub)classes is provided in Supplementary File 1. In some cases (MG, short acyl chain TG, CAR) coelution with lipid standards was used to validate their identity (Supplementary Figure 2). Such rigorous curation of lipid annotations allowed us to resolve lipid classes, which are often not discriminated. For instance, using defined set of specific fragment ions as well as retention time mapping, unambiguous identification of acyl-, alkyl-, and alkenyl-PL as well as lysoPL became possible (Supplementary File 1).

Overall, we obtained a list of 1636 lipids representing 23 lipid subclasses (Figure 2D; Table S1). TG display the highest lipidome complexity by making up 63.2% of all identified lipids followed by PL (16.8%), SP (9.8%), DG (5.4%), lysoPL (2.8%), acylcarnitines (1.7%) and CEs (0.4%). Eventually we achieved a three dimensionally curated (RT-MS-MS/MS), high confidence lipid inventory of human WAT covering all major lipid classes including glycerolipids, phospholipids, sphingolipids, cholesteryl esters, and acylcarnitines, representing the most detailed description of human WAT lipidome to date.

### Quantitative analysis of human WAT lipidome

Significance of accurate quantification for harmonization of lipidomics data was recently underlined by the lipidomics community (Liebisch et al., 2019). However, several analytical challenges need to be accounted for when aiming for accurate lipid quantification. MS signal intensity is molecular structure dependent and as such requires application of internal standards (ISTDs) to support accurate quantification. Highest possible quantitative accuracy can be obtained by using isotopically labelled ISTDs for each molecular species in the sample, which is unfortunately still not feasible at the whole lipidome level. Here we performed semi-absolute quantification of identified WAT lipids using lipid subclass specific ISTDs at concentrations close to the endogenous analytes (Figure 3A). WAT specific ISTD mixture was designed to cover the whole range of identified lipid classes by using different classes of PLs (deuterated PC, PE, PS, PG, PI, LPC, LPE), GL (^13^C-labeled DG, TG), CE (deuterated CE), SP (natural Cer, dihydroceramides (DhCer), deoxyceramides (DeoxyCer), phytoceramides (PhytoCer), hexosylated ceramides (GlcCer, LacCer), SM), CAR (deuterated) and free fatty acids (^13^C-labeled). Isotopically labeled or naturally occurring lipids absent in WAT were used (Table S2). Next, 6-point calibration curves were generated for each ISTD spiked in the adipose tissue matrix to determine the linear response range (Supplementary Figures S3 and S4). Final ISTD amounts in the mixture were chosen to represent intensity close to the native lipids of the corresponding lipid class while still displaying a linear behavior of ISTD in the concentration-response relationship. Pooled WAT samples were spiked with the designed ISTD mixture prior to lipid extraction and fractionation, and analyzed using full scan LC-MS or PRM platforms utilizing three type of LC separations as described above (C30 RPC, C18 RPC, and HILIC; 90 LC-MS analysis).

**Figure 3:**
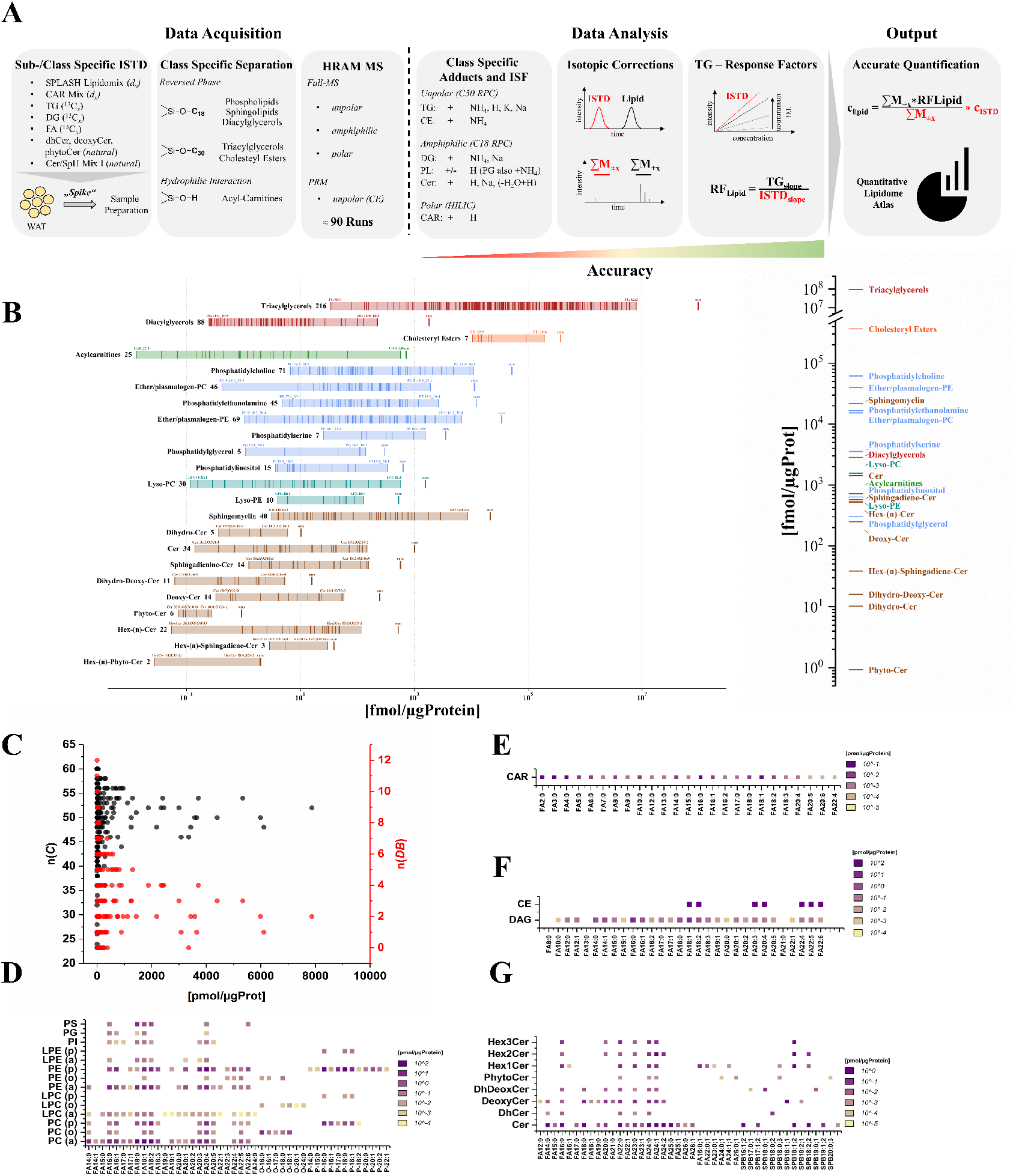
Quantitative representation of human WAT lipidome and description of used analytical strategy. **A:** Schematic depiction of the quantitative lipidomics workflow. Subclass specific lipid internal standards (ISTD) were spiked into WAT samples prior to lipid extraction. Lipid subclass specific liquid chromatography (LC) and high resolution, accurate mass (HRAM) MS enabled quantification of the highest possible number of lipid species. Lipid class specific differential adduction and MS stability was assessed by quantification of all adducts and in-source fragments (ISF) of the corresponding lipid molecular species. Type I isotopic correction as well as correction for incomplete isotopic labeling of the ISTD were additionally applied to increase quantitative accuracy. Quantification of triacylglycerol (TG) was further refined by the generation of molecular structure dependent response factors (RF). The use of lipid class-/subclass specific ISTDs allowed for confident, accurate quantification. **B:** Quantitative distribution of lipid class and corresponding lipid molecular species within subclasses of human WAT. Total lipid class concentration is represented by bold lines (*SUM*) and each single lipid molecular species is represented by thin lines. **C:** Distribution of single TG based on bulk fatty acid chain length (expressed as total carbon number, n(C); black dots) and unsaturation (expressed as double bond number, n(DB); red dots). Quantitative distribution of fatty acids, fatty alcohols, fatty vinyl alcohols and sphingoid bases across **D:** phospholipids, **E:** acylcarnitines, **F:** cholesteryl esters and diacylglycerols and **G:** sphingolipids.

All obtained data were corrected for lipid isotopic patterns (type I) as well as for incomplete isotopic enrichment of deuterated ISTDs (Figure 3A). Since not only the lipid class but also the fatty acyl chain composition in a given lipid will determine its MS response, we defined acyl chain specific response factors for the most abundant and diverse WAT lipid class, TG, which were used to increase quantitative accuracy of TG (Supplementary Figure S5). Thus, using comprehensive separation strategy coupled to HRAM MS detection and in-house designed ISTD mixture customized to WAT, we performed semi-absolute quantification of human WAT lipidome covering 522 lipid molecular species as well as 215 TG quantified at lipid class level, providing the most detailed semi-quantitative mapping of human adipose tissue lipidome to date (Figure 3B, Table S3).

### Adipose tissue lipids span over a wide concentration range and display lipid class specific fatty acyl signatures

Quantified WAT lipids displayed a huge dynamic range of concentrations from 12 amol/µg protein (CAR 20:5) up to 8 nmol/µg protein (TG 52:2), spanning over eight orders of magnitude (Table S3). Total TG concentration (96.2 nmol/µg protein) within WAT is overarching other lipid classes by two orders of magnitude, resembling their role as energy storage lipids. Importantly, concentrations of individual TG species ranged over five orders of magnitude showing the molecular species dependent abundance, with just 20 most abundant TG resembling over 71% of the total TG amount. Those top 20 TG contained primarily saturated and monounsaturated fatty acyl chains with an average of two double bonds per three acyl chains (Figure 3C). CE were the second most abundant lipid class of which CE 20:4 was the most concentrated (Figure 3F).

Unpolar lipids were followed by PC, PE, and SM. Interestingly, PC and PE lipids showed inverted distribution of the corresponding subclasses (Figure 3D). Thus, diacyl-PC were ~ 4.5 times more abundant than ether-PCs, whereas ether-PEs were ~ 3 times more abundant than diacyl-PE. Specifically, plasmalogen PE were the most abundant lipid subclass within PE lipids. Closer inspection of the fatty acyl chain distribution revealed class/subclass specific differences. For PC lipids, the fatty acyl chain abundance was largely similar between acyl-alkyl-, and alkenyl-species, whereas for PE a higher concentration of PUFA-containing alkenyl-PE (plasmalogens) was observed over diacyl-PE. Interestingly, plasmalogen lipids were previously reported to be enriched in brain and heart tissues. Here, we demonstrate that in human WAT plasmalogen PE represent the 4th most abundant lipid class with a total concentration of 11.3 pmol/µg protein and the most abundant molecular species of plasmalogen PE are rich in PUFA. LysoPC and lysoPE lipids were one order of magnitude less abundant than PCs and PE, with acyl chain composition similar to the corresponding diacyl-species, indicating active lipid remodeling within these PL classes via the Lands cycle (Figure 3D). Other PL were detected only as diacyl species with PS being the next abundant class followed by PI and PG. All minor PL lipids showed fatty acyl chain distribution characteristic for those classes (Paul et al., 2019; Skotland and Sandvig, 2019). Thus, most abundant PS molecular species were PS 18:0_18:1 and PS 18:0_18:2, whereas PIs were rich in FA 20:4. The most abundant PG was PG 18:1_18:1 (Figure 3D).

DG and acylcarnitines (CAR) displayed quantities in the medium abundance range reasoning for their role as intermediates within lipid metabolism. Indeed, DG acyl chain distribution was quite similar to PL indicating the role of DG as structural precursors of membrane lipids (Figure 3E). The most abundant acylcarnitine was CAR 2:0 in line with its proposed function as a sink for acetyl equivalents, accumulating due to the constant energy surplus, to prevent complete CoA consumption, especially in highly metabolically active tissues as WAT (Figure 3E) (Lopaschuk et al., 1994; Ramsay et al., 2001; Schooneman et al., 2013). Other abundant species CAR 3:0 and CAR 5:0 possibly originate from branched chain amino acid oxidation (Newgard, 2012; Newgard et al., 2009). Additionally relatively high concentrations of CAR 16:0, 16:1, 18:0, 18:1, and 18:2 were observed indicating acylcarnitines shuttling of medium chain fatty acids to mitochondrial β-oxidation (Schooneman et al., 2013). This is, to the best of our knowledge, so far the first quantitative assessment of CAR species within human WAT further supporting its role as a highly metabolically active organ.

Another lipid class of high metabolic importance often implicated in development of obesity related pathologies are ceramides (Cer) (Figure 3G). We found preferential incorporation of saturated or monounsaturated long and very-long chain fatty acids in Cer lipids, typical for this lipid class. The most abundant (203 fmol/µg protein) Cer in human WAT was Cer 34:1;O2 represented by two isomeric species – Cer 18:1;O2/16:0 and Cer 16:1;O2/18:0. Cer 18:1;O2/16:0 and corresponding ceramide synthase (CerS6) were previously associated with weight gain and glucose intolerance (Turpin et al., 2014). On the other hand, Cer with sphingoid bases others than SPB 18:1;O2 are rarely monitored, thus usually not reported (Chaurasia et al., 2016; Turpin et al., 2014). Overall, we demonstrated previously unanticipated diversity of Cer subclasses in human WAT. Cer were represented by species with varying length of sphingoid bases of which SPB 18:1;O2 and SPB 16:1;O2 were the most abundant. Dihydroceramides (dhCer), precursors in *de novo* Cer biosynthesis, were one order of magnitude lower than ceramides themselves. The next most abundant Cer subclass were hexosylated Cer derivatives, closely followed by sphingadienine-Cer, a class of lipids only recently discovered and monitored in human blood plasma (Karsai et al., 2020). Finally, to our surprise, human WAT was enriched in deoxy-Cer lipids. This potentially cytotoxic Cer subclass was detected in human blood plasma where it represents a minor (0.1-0.3%) fraction of total sphingoid bases. Here, WAT contained significant amounts of deoxy-Cer lipids, corresponding to 12.6% of all Cer subclasses. Considering close interconnection of sphingolipid metabolic pathways, and already established functional differences of structurally diverse Cer lipids (Chaurasia et al., 2016; Sokolowska and Blachnio-Zabielska, 2019), AdipoAtlas significantly enriches the current knowledge on the human WAT lipid composition.

### PUFA containing TG are specifically upregulated in obese WAT

Having detailed semi-quantitative map of human WAT in hands, we compared global lipidome compositions of subcutaneous and visceral WAT from lean and obese individuals. As expected, we found a statistically significant upregulation of TG in obese adipose tissue (Figure 4A). Interestingly, all obesity upregulated TG contained at least one PUFA residue with FA 20:4, FA 20:5, FA 22:5 and FA 22:6 acyl chains being the most upregulated PUFAs in that respect (Figure 4B). Conversely, TG species containing mostly SFA and MUFA residues were markedly decreased in the adipose tissue of obese individuals (Figure 4A). Correlation analysis further confirmed strong co-regulation of PUFA-TG (Figure 4C). Previously, obesity driven global accumulation of long-chain PUFA containing TG was demonstrated in murine models and human biopsies (Cao et al., 2008; Liesenfeld et al., 2015; Pietiläinen et al., 2011). Increased activity of the fatty acid elongase Elovl6 was proposed to play a role as a common phenomenon in rodent and human adipose tissue in response to excessive nutrient consumption (Liesenfeld et al., 2015). This suggests that besides upregulation of total enzymatic fatty acid elongation/desaturation machinery, the generation of specific TG molecular species is regulated during obesity development, and opens up the question how and why specific lipogenic enzymes generate distinct TG species and what is their role in obesity development.

**Figure 4:**
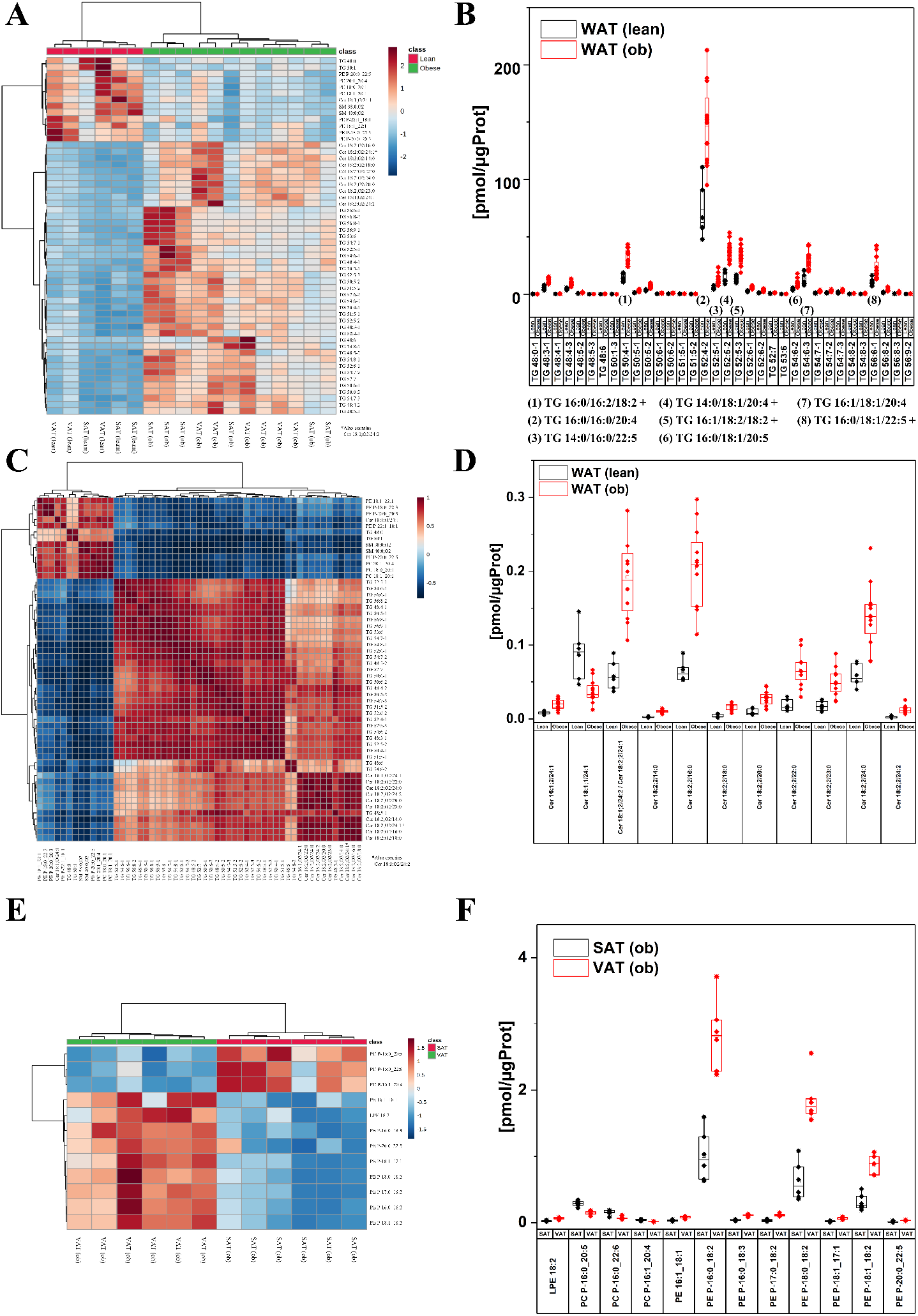
Human white adipose tissue (WAT) displays distinct lipidome profiles depending on WAT depot (visceral vs subcutaneous, VAT vs SAT) and phenotype (lean vs obese, lean vs ob). The global lipidome was quantified for representative pools of WAT from visceral and subcutaneous depots of lean (*n*=5) and obese (*n*=81) individuals. **A:** Heatmap displaying statistically significantly regulated lipid molecular species between the WAT of obese and lean patients. **B:** Concentration of differentially regulated triacylglycerols (TG) between obese and lean WAT. **C:** Pearson correlation of significantly regulated lipids between lean and obese WAT. **D:** Concentration of statistically significantly regulated ceramide (Cer) species between lean and obese WAT. **E:** Heatmap displaying statistically significantly regulated lipid molecular species between obese visceral (VAT(ob)) and obese subcutaneous (SAT(ob)) depots. **F:** Concentrations of phospholipid and plasmalogen phospholipid species that are statistically significantly regulated between VAT (ob) and SAT (ob). Statistical significance was determined by Student’s t-test (FDR adjusted) with a cutoff of p ≤ 0.05 and a minimum fold change ≥ 2.

### Increase in sphingadienine containing ceramides is a hallmark of obese adipose tissue

Obesity is one of the main risk factors for the development of type 2 diabetes mellitus, precluded by organ specific or systemic insulin resistance (Wondmkun, 2020). Ceramides are well known mediators of insulin resistance with Cer levels in different organs reflecting insulin sensitivity state (Sokolowska and Blachnio-Zabielska, 2019). Here, in the obese adipose tissue we found a marked upregulation of ceramides with FA 24:1 acyl chains, and ceramides with the unusual sphingoid base sphingadienine (SPB 18:2;O2) (Figure 4A and D). Sphingadienine (SPB 18:2;O2) ceramides are a so far functionally undescribed class, synthesis of which has been proposed in adipose tissue but not confirmed until now (Karsai et al., 2020). Upregulated sphingadienine Cer contained diverse range of esterified acyl chains (from FA 14:0 up to FA 24:0) (Figure 4D). Correlation analysis showed that there is a strong co-regulation between all upregulated SPB 18:2;O2 containing ceramides (Figure 4B) indicating an increased biosynthesis of this unusual sphingoid base or/and an increased acylation rate of the sphingadienine base.

### Plasmalogen phospholipids are a depot specific signature in acquired obesity

Encouraged by the fact that AdipoAtlas provided new insights in the lipidomics signature of obese vs lean adipose tissue, we next looked at the possible difference in lipid compositions of obese subcutaneous and visceral AT depots. Interestingly, a clear discrimination of SAT and VAT depots was possible based on their respective plasmalogen PL signatures (Figure 4E). Thus, higher amounts of plasmalogen PC with long chain PUFA (e.g. FA 20:4, FA 20:5, FA 22:6) were characteristic to subcutaneous obese WAT, whereas plasmalogen PE accumulated in visceral obese WAT. The majority of VAT upregulated plasmalogen PE carried 18 carbon long fatty acyl chains (Figure 5B). Differential regulation of plasmalogen PE was already indicated in previous studies (Barchuk et al., 2020; Liesenfeld et al., 2015; Pietiläinen et al., 2011). Here we further demonstrated involvement of plasmalogen PC, and, importantly, fatty acyl specificity within the regulated lipid species.

**Figure 5:**
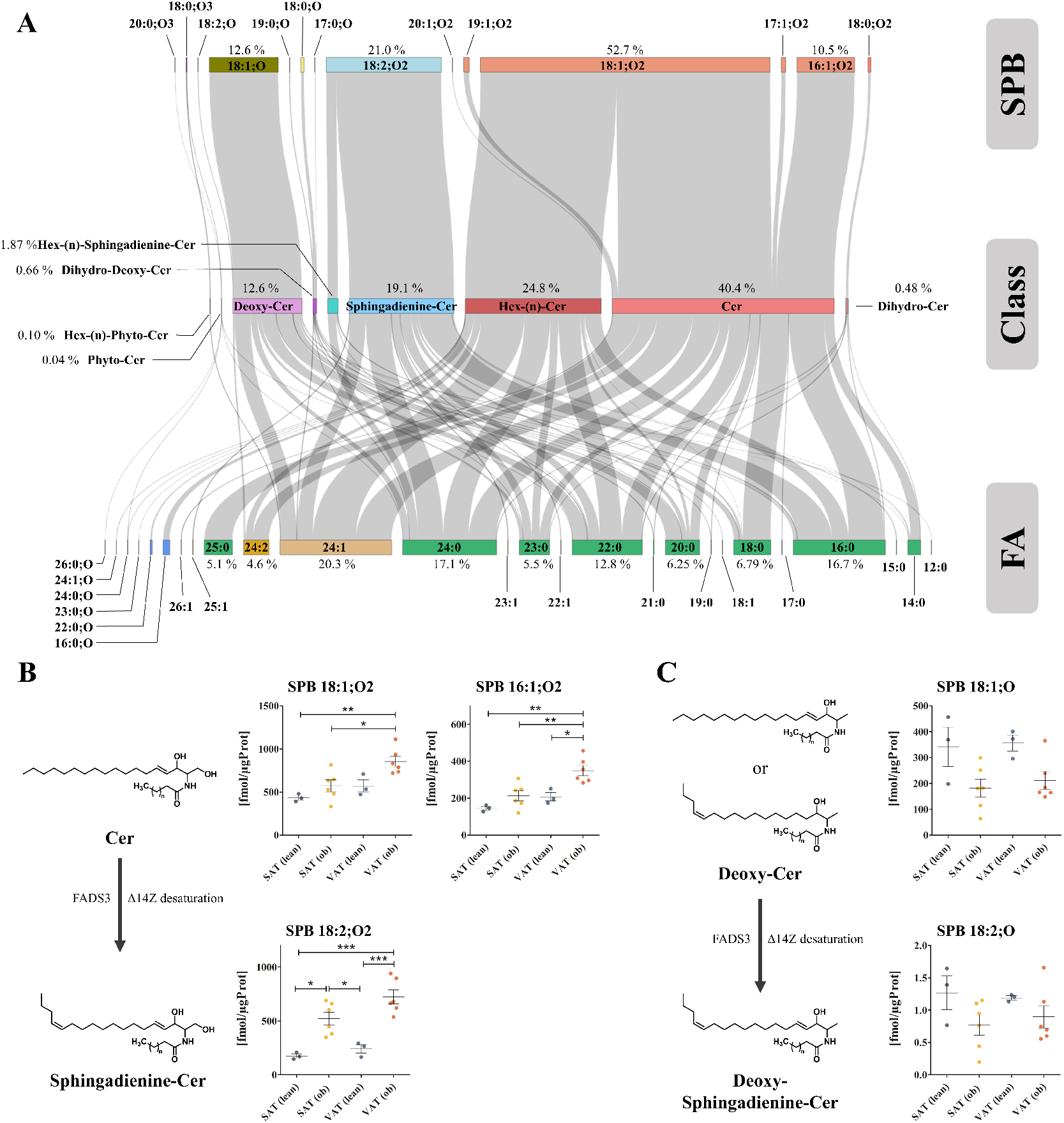
Complexity of the human white adipose tissue (WAT) sphingolipidome: **A:** Sankey plot displays the concentration of ceramide (Cer) subclasses, its corresponding esterified sphingoid bases (SPB) and fatty acids (FA). Depicted concentrations were calculated by averaging concentrations of WAT from subcutaneous and visceral depots of lean and obese patients in order to reflect the general WAT sphingolipidome. Length of boxes correspond to the determined concentrations. **B:** Differential regulation of sphingosine and sphingadienine SPBs over Cer subclasses in obese (ob) and lean tissues from visceral (VAT) and subcutaneous (SAT) depots. **C:** Differential regulation of deoxy-sphingosine and deoxy-sphingadienine SPBs over Cer subclasses in obese (ob) and lean tissues from visceral (VAT) and subcutaneous (SAT) depots. Statistical significance was calculated by ANOVA. * p ≤ 0.05; ** p ≤ 0.01; *** p ≤ 0.005.

## Discussion

Deep profiling of tissue-specific lipidomes is essential to support our understanding of human biology by elucidating not only tissue/organ specific lipid remodeling mechanisms but also the cross-talk between different tissues and its impact on systemic regulation of the lipid metabolism. In contrast to robust screening applications necessary for the analysis of large sample cohorts and potential clinical translation, deep lipidomics profiling of a particular tissue, organ or cell type cannot be performed in a high-throughput manner and requires rigorous tissue-tailored optimizations and application of multiple orthogonal analytical workflows. Here we provide the example of analytical strategy targeting deep lipidomic profiling of human WAT, which can be transferred and adapted for the generation of reference lipidomes from any other human tissues. The workflow included three main steps to ensure non-discriminative deep lipidome coverage providing qualitative and quantitative inventory of human WAT lipidome:

I. Tissue-tailored lipid extraction and fractionation to ensure non-discriminative coverage of all lipid (sub)classes was optimized by testing several protocols using quantitative high-performance thin layer chromatography (qHPTLC) as a robust readout method allowing fast quantitative assessment of extraction and fractionation efficiencies.
II. Rigorous identification of lipid molecular species was performed by employing multiple separation (C18 RPC, C30 RPC, HILIC) and MS analysis platforms (+/- DDA, AcquireX, PRM). LC-MS/MS analysis of four samples pools (SAT and VAT from lean and obese individuals) resulted in over 110 datasets used for lipid identification. Importantly, identification was performed using three independent software tools and was followed up by manual curation of MS/MS spectra and retention time mapping. Such a meticulous strategy was rewarded by accurate identification of over 1600 lipid molecular species, including rarely resolved ones. For instance, using specific sets of fragment ions and subclass specific elution order, unambiguous identification of diacyl-, alkyl-, and alkenyl-PC and PE lipids was achieved. Furthermore, three-dimensional (LC-MS-MS/MS) curation including control for in-source fragmentation artefacts allowed to uncover unexpected diversity of WAT sphingolipidome. To support accurate lipid identification at the molecular species level we provide here a summary of lipid fragmentation patterns and retention time maps (Supplementary File 1), which can be used by other researchers aiming for deep lipidome mapping.
III. Semi-absolute quantification of human WAT was based on lipid class specific ISTDs that were carefully selected to represent the diversity of endogenous lipids. To this end, a tissue-tailored ISTD mixture was designed and validated. Quantitative data were processed using several types of isotopic corrections, adducts and in-source fragments. Additionally, response factors accounting for the distribution of diverse acyl chains in TG lipids were calculated and applied to enhance the accuracy of the quantification results. That allowed us to provide semi-quantitative values for 23 lipid subclasses and for the first time to appreciate the extraordinary dynamic range of lipid concentrations within and between different lipid subclasses in human WAT.

Human WAT reference lipidome was reconstructed using pooled samples representative of SAT and VAT depots from lean and obese individuals. Although application of pooled samples limits a detailed assessment of diseases specific lipid alterations, several depot and phenotype specific lipid features became apparent. Accumulation of TG lipids in WAT is a known hallmark of obesity. AdipoAtlas demonstrated a large diversity of WAT TG represented by 1029 molecular species. Interestingly, TG also were characterized by the largest dynamic range of concentrations with only 20 TG covering over 70% of total TG concentration. Thus, the most abundant TG 52:2 corresponded to 6.8 µg/µg of AT proteins. Although top 20 most abundant WAT TG were mostly saturated (2 double bonds per 3 acyl chains on average), PUFA rich TG species were significantly upregulated in obese WAT. These results are in line with previous studies illustrating the enrichment of PUFA TG in obese adipose tissue (Pietiläinen et al., 2011; Liesenfeld et al., 2015). However mechanistic understanding of such specificity remains limited.

One of the main discoveries provided by AdipoAtlas is the previously uncovered diversity of WAT sphingolipidome. Sphingolipids, and especially ceramides, recently attracted a lot of scientific attention. Tissues and blood plasma levels of these lipids emerge as important predictors of metabolic malfunction in human pathologies associated with altered lipid metabolism. Ceramide accumulation was associated with deleterious metabolic outcomes including insulin resistance, ectopic lipid accumulation, apoptosis and fibrosis (Turpin-Nolan and Brüning, 2020). Ceramide biosynthesis and catabolic pathways were identified as favourable targets for pharmacological intervention in metabolic diseases. Thus, knockdown or small molecule-based inhibition of serine palmitoyltransferase (SPTLC), dihydroceramide desaturase 1 (DEGS1), selected isoforms of ceramide synthases (CerS), as well as overexpression of ceramidases (acid ceramidase and adiponectin receptor) all showed beneficial effects in metabolically challenged mice (Chaurasia et al., 2019; Correnti et al., 2014; Glaros et al., 2008; Holland et al., 2007).

Elevated Cer levels were found in WAT in obesity and obesity associated diseases including type 2 diabetes mellitus (Chaurasia et al., 2016), fatty liver diseases (Kolak et al., 2007), and metabolic syndrome (Choromańska et al., 2019). Details on Cer tissue depot specificity (SAT vs VAT) remain less obvious. Majority of the human studies used either SAT or VAT, and reports directly comparing SAT vs VAT Cer levels are rare. However, current data indicate the significance of Cer acylated with FA 16:0 as the molecular species associated with adverse metabolic outcomes (Turpin et al., 2014). This observation is further supported by transcriptomics analyses showing significant upregulation of CerS6, an isoform of ceramide synthase preferentially acylating FA 16:0 into Cer, in WAT of obese individuals (Turpin et al., 2014). It is important to note that most of the studies reporting Cer levels in human WAT utilize targeted detection methods that cover only “classical” species (Cer, dhCer, and sometimes their glycosylated derivatives) in which SPB 18:1;O2 (sphingosine) is acylated with different fatty acyl chains. Using our advanced analytical workflow, we demonstrated that WAT sphingolipidome is more complex, with SPB 18:1;O2 Cer representing only 69.2% of all classical Cer (Figure 5). AdipoAtlas facilitated identification of four additional bases including SPB 16:1;O2 (26.1 %), SPB 19:1;O2 (2.6%), SPB 17:1;O2 (2.1%) and SPB 20:1;O2 (0.7%). Both SPB 18:1;O2 and SPB 16:1;O2 were elevated in obese WAT and their levels were higher in VAT in comparison to SAT of lean and obese origin (Figure 5B). Overall, “classical” Cer represent only 40% of total Cer species quantified in WAT. The next most abundant subclass was glycosylated Cer (25%) containing up to three hexoses (Hex(n)Cer).

Interestingly, we furthermore identified two high abundant “atypical” Cer classes in human WAT, namely deoxyCer and sphigadienineCer. DeoxyCer are acylated derivatives of 1-deoxy-sphingosine (SPB 18:0;O), synthesised by SPT from palmitate and alanine instead of serine (Duan and Merrill, 2015). They are typically considered as toxic by-products in Cer metabolism, as they can neither be degraded via classical Cer catabolic pathways nor be converted to SM and glycoCer species. DeoxyCer were only recently identified in human adipose tissue with higher levels in VAT relative to serum, particularly in obese individuals with type 2 diabetes mellitus (Hannich et al., 2020). Plasma levels of deoxyCer were positively associated with age, BMI and waist-to-hip ratio (Beyene et al., 2020; Othman et al., 2012) proposing them as a hallmark of metabolic complications. Interestingly, in plasma SPB 18:1;O represents only a minor (0.1-0.3%) fraction (Othman et al., 2012), whereas AdipoAtlas revealed ten times higher values in human WAT (12.6%) (Figure 5A). This suggests a significant enrichment of these potentially toxic species in WAT although the exact role of deoxyCer remains to be uncovered. Previously believed to be mainly of hepatic origin, deoxyCer were shown to be directly synthesized by adipocytes during differentiation (Hannich et al., 2020). Thus, AdipoAtlas as well as recently reported data (Hannich et al., 2020), allows to propose human WAT as an important reservoir and a source of potentially toxic deoxyCer.

SphigadienineCer represented another abundant Cer subclass in human WAT (Figure 5). In comparison to “classical” Cer, sphigadienineCer contain one more double bound at the position Δ14Z (SPB 18:2;O2). Previously, SPB 18:2;O2 Cer were shown to reflect metabolic fitness due to inverse association with homeostatic model assessment for insulin resistance, BMI and incidence of cardiovascular events (Chew et al., 2019; Othman et al., 2015). Although the existence of SPB 18:2;O2 is long known, the enzyme (fatty acid desaturase 3; FADS3), responsible for the introduction of Δ14Z double bond, was discovered only recently (Karsai et al., 2020). According to gene expression data from GTEx portal (https://gtexportal.org), highest FADS3 expression levels were found in peripheral nerve, aorta, and WAT. Gender specific expression analysis further unravelled highest expression of FADS3 in female WAT. So far sphigadienineCer were not characterized in adipose tissue. Here we report that they represent 19% of all Cer subclasses within human WAT (Figure 5A). Moreover, 18:2;O2 Cer were elevated in both SAT and VAT obese depots with significant enrichment of this lipid class in VAT vs SAT. Importantly, elevated levels of 18:2;O2 Cer represented a specific signature of obese WAT showing statistical significance even for the pooled samples utilized in this study (Figure 4D). Although depot and phenotype specific increase in sphigadienineCer displayed the trend similar to Cer, their fatty acyl chain distribution was somewhat different, with a lower portion of FA 18:0 acylated into 18:2;O2 Cer in comparison to 18:1;O2 Cer. Our results strongly suggest that sphingadieneCer accumulation is a hallmark of obesity. Previously published data showed significant gender-specificity for deoxy- and sphigadienineCer subclasses with SPB 18:2;O being more abundant in men and SPB 18:2;O2 Cer in women. Interestingly, we observed inverse correlation between those lipid classes in lean and obese WAT (Figure 5B), which might be explained by 2/3 prevalence of female WAT donors in our sample pools. Further analysis of individual samples based on AdipoAtlas list will provide deeper insights in gender and disease specific signatures of ceramide lipids.

Another emerging class of lipids potentially involved in regulation of cellular and systemic lipid homeostasis are etherPL (ePL), including plasmalogens (pPL). Previously, brain and heart were identified to be rich in pPL. AdipoAtlas showed that ePC and ePE lipids composed 41% of total PC and PE. Specifically, we demonstrated that PUFA rich ePE (40 pmol/µg protein) represented the 4th most abundant lipid class in human WAT, closely following the most abundant PC phospholipids (62 pmol/µg protein). We identified depot specific signatures of ePL with higher levels of PUFA pPC in SAT, and enrichment of C18 fatty acyl chain containing pPE in VAT. Ether lipids, and plasmalogens especially, play a central role in lipid quality control, adaptive responses to the change in lipid saturation levels, and maintenance of membrane fluidity (Jiménez-Rojo and Riezman, 2019). pPE compose over 20% of inner leaflet of plasma membrane (Lorent et al., 2020), making them important players in membrane remodelling during adipocyte hypertrophic growth (Pietiläinen et al., 2011). Moreover, levels of circulating plasmalogens were inversely associated with hypertension, prediabetes, type 2 diabetes mellitus, cardiovascular diseases, and obesity (Paul et al., 2019). Interestingly, inverse co-regulation of ether lipids and sphingolipids was recently demonstrated, with depletion of ePL leading to the Cer accumulation and vice versa (Jiménez-Rojo et al., 2020). AdipoAtlas provides an inventory of ePL molecular species including resolved diacyl-, alkyl- and alkenyl-PL which can be used as a resource for close, targeted follow up of this inverse correlation in larger sample cohorts.

Overall, deep lipidomic profiling allowed to reconstruct human WAT reference lipidome. AdipoAtlas provides an inventory of over 1600 lipid molecular species from 23 lipid (sub)classes fortified by their semi-quantitative values in two WAT depots (subcutaneous and visceral) from lean and obese individuals. That allowed us to demonstrate an amazing diversity of adipose tissue lipids together with assessment of their quantities between and within lipid classes. Several important lipid signatures characteristic for obesity were discovered or reproduced pointing out qualitative and quantitative accuracy of the applied methodology. With AdipoAtlas freely available for all researchers interested in WAT biology, it can be further used to design human WAT-specific high-throughput experiments targeting quantification of any given lipid (sub)class in large sample cohorts. Moreover, AdipoAtlas will provide so far missing scaffold for systems biology integration of lipidomics data via reconstruction of lipid-centric genome scale metabolic models, linking big omics data with identification of disease characteristic metabolic and signaling pathways.

## Acknowledgments

Financial support from the German Federal Ministry of Education and Research (BMBF) within the framework of the e:Med research and funding concept for SysMedOS project (to MF) are gratefully acknowledged. We thank Prof. Ralf Hoffmann (Institute of Bioanalytical Chemistry, University of Leipzig) for providing access to his laboratory.

## Author Contributions

MF conceived the project, guided the research, assisted with the experiments and data interpretation, and wrote the manuscript. ML designed and performed most of the experiments, analyzed and interpreted data, wrote the manuscript. GA performed lipid identification including manual annotation and retention time mapping. ZN performed lipid identification, dataset merging, prepared the illustrations. AC performed parts of the LC-MS/MS experiments. JS assisted with ^31^P NMR and analysis of lipids from TLC plates. MB provided human WAT samples. All authors edited and approved the manuscript.

