## Supplemental File 1 for "AdipoAtlas: A Reference Lipidome for Human White Adipose Tissue"

### Supplementary File 1.
### Identification strategy used for lipid identification from LC-MS/MS datasets

**CAUTION:** Retention times, elution orders, adducts formation and their relative abundance, extent of in source fragmentation (ISF), and fragmentation patterns are all described for reversed phase chromatography [C18 or C30 (stationary phase) and water-acetonitrile-isopropanol supplemented with formic acid and ammonium formate (mobile phase)] coupled on-line to Q Exactive Plus MS or Fusion Lumos systems with beam type CID (here HCD). Note, that other types of chromatography, MS and MS/MS setups might result in different values and patterns.

#### Phospholipids

##### 1. Manual annotation of Lysophosphatidylcholines (LPC)

Manual confirmation to assure accurate identification of A-LPC, O-LPC and P-LPC (A- vs O- vs P-LPC) subclasses was based on specific fragment ions, their ratio and retention time mapping.

**1.1. Monitored adducts:** LPC were monitored as protonated or formate adducts in positive and negative ion modes, respectively. $[M-CH_3]^-$ ions, usually formed as the result of in-source fragmentation (ISF) by the collisional decomposition of anionic format adducts with the loss of neutral methyl format at the intermediate region of the instrument between the atmospheric pressure of the ESI source and the vacuum parts of the analyzer, were below 1% relative to the format adducts even for the most abundant LPC lipids and thus were not considered for the identification. However, $[M-CH_3]^-$ ions might be more prominent on the QTOF instruments operating with a higher pressure at the front end of the instrument or Orbitrap-based platforms with ion funnel configuration of the transmission devices[1–3].

**1.2. Fragmentation patterns:** General fragmentation pattern and representative MS/MS spectra for LPC ionized in positive and negative modes are illustrated below:

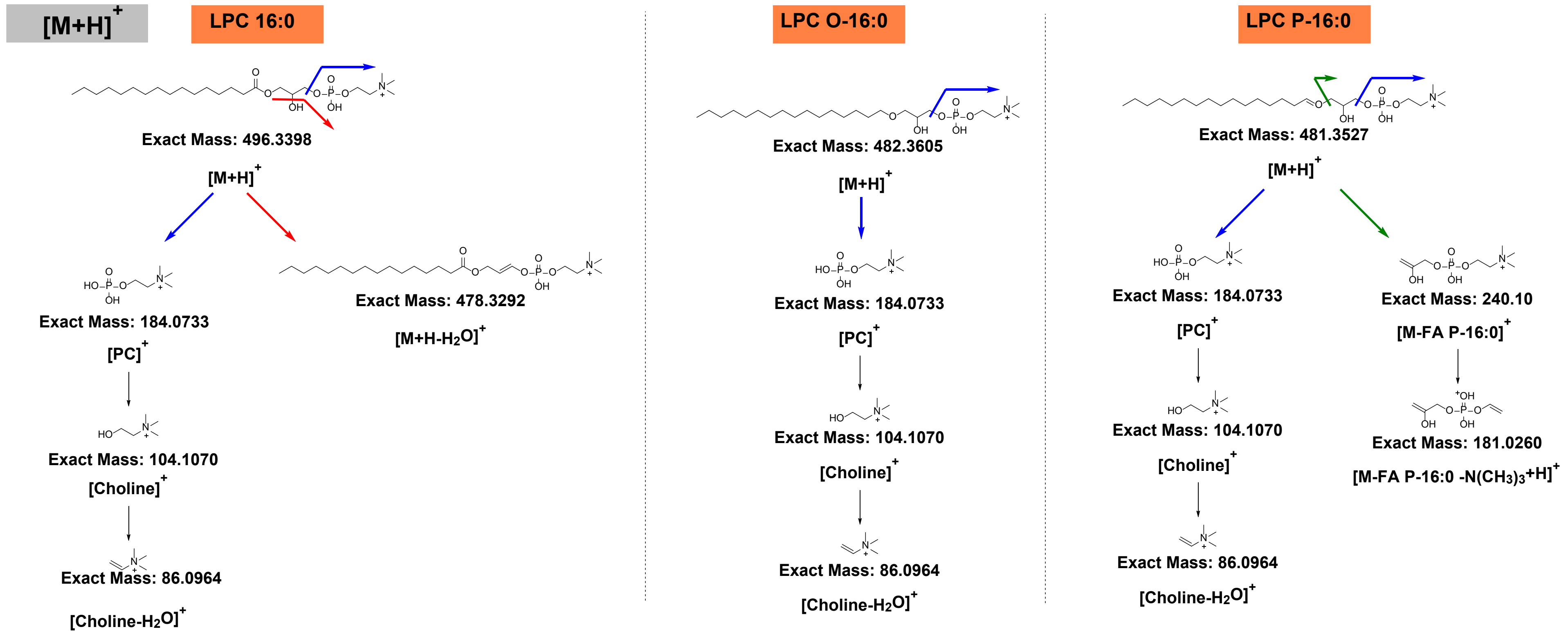


**Scheme 1.1.** General HCD fragmentation pattern of protonated acyl, alkyl and alkenyl linked LPC $[M+H]^+$.

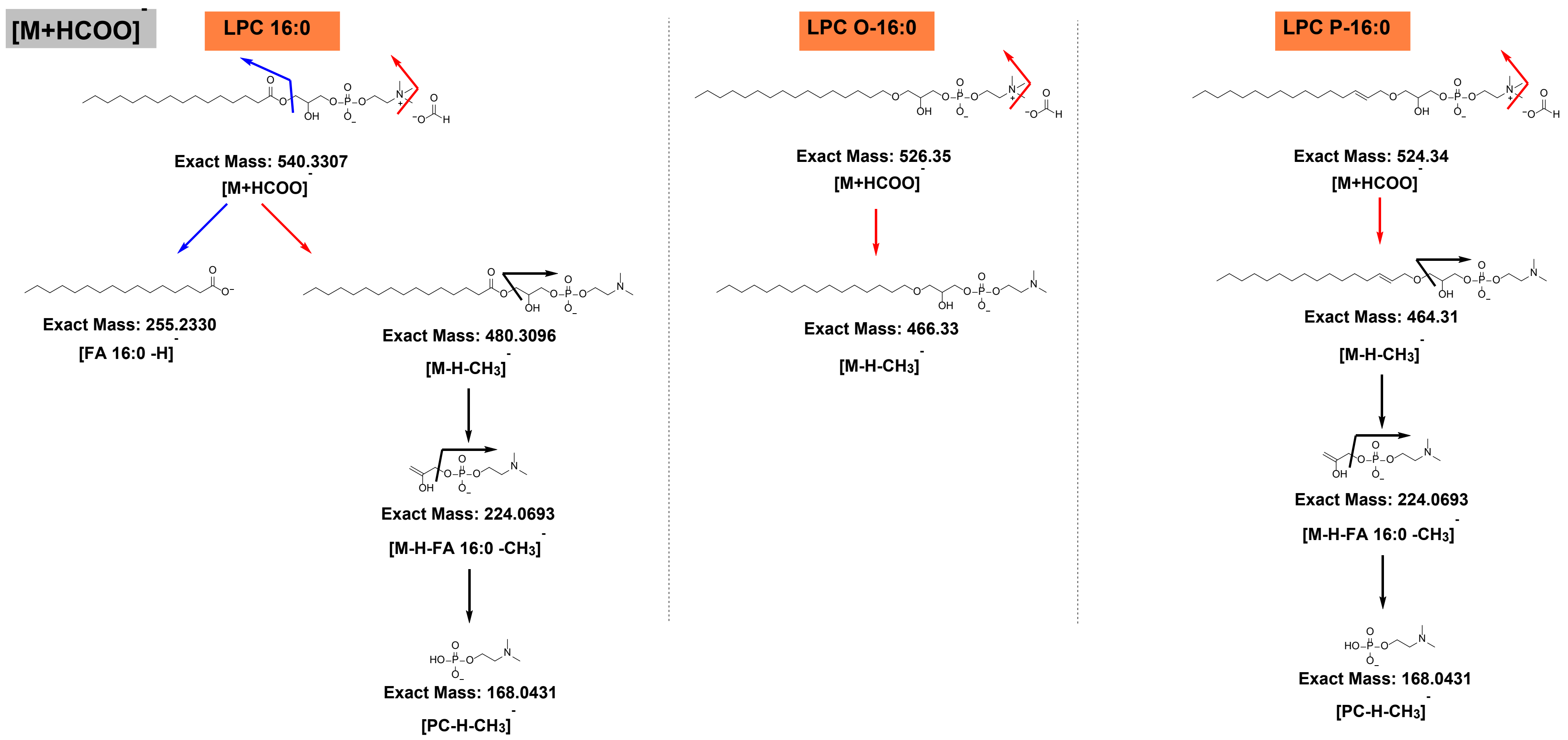


**Scheme 1.2.** General HCD fragmentation pattern of acyl, alkyl and alkenyl linked LPC format adduct $[M+HCOO]^-$.

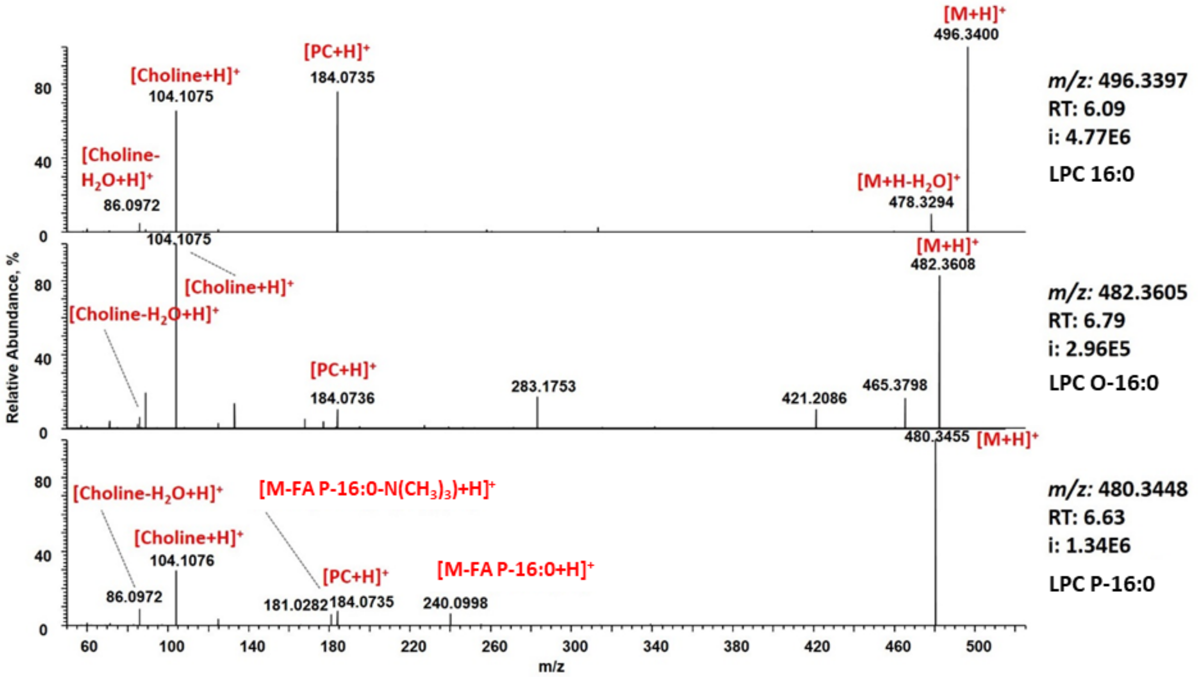


**Figure 1.1.** Representative HCD spectra of protonated LPC 16:0 (top), LPC O-16:0 (middle), and LPC P-16:0 (bottom).

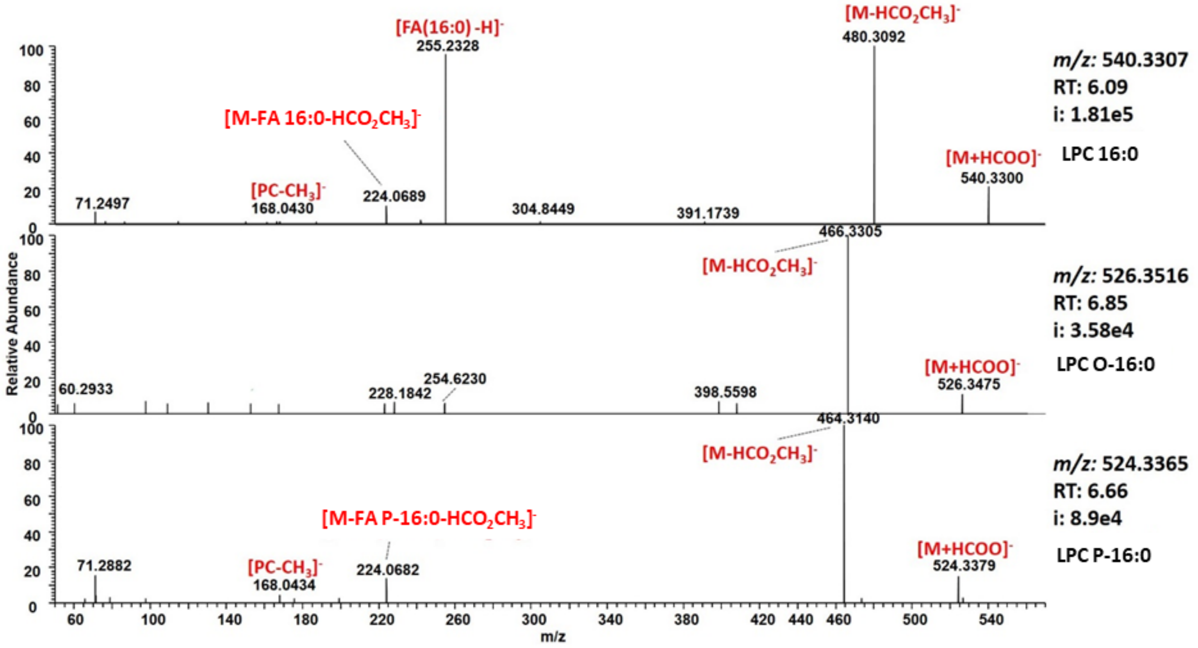


**Figure 1.2.** Representative HCD spectra of formate adduct of LPC 16:0 (top), LPC O-16:0 (middle), and LPC P-16:0 (bottom).

**Table 1.1.** Summary of LPC neutral loss (NL) and fragment ions (FI) obtained by positive and negative ion mode HCD.

| NL/FI | A-LPC | O-LPC | P-LPC |
|---|---|---|---|
| **Positive ion mode HCD** | | | |
| NL: -18 (-water) | + | | |
| FI: 240 (dehydroglycerophosphocholine) | | | + |
| FI: 181 (dehydroglycerophosphocholine - trimethylamine) | | | + |
| FI: 184 (phosphocholine) | + | + | + |
| FI: 104 (choline) | + | + | + |
| FI: 86 (dehydrocholine) | + | + | + |
| **Negative ion mode HCD** | | | |
| NL: - 60 (methyl format) | + | + | + |
| FI: fatty acyl anion | + | | |
| FI: 224 (demethylated dehydroglycerol phosphocholine) | + | | + |
| FI: 168 (demethylated phosphocholine) | + | | + |

**Positive ion mode HCD:**

- Water loss from the precursor is usually observed only for A-LPC.
- Fragment ion at *m/z* 240 formed by the NL of aldehyde (and low abundant fragment at *m/z* 181 formed by a consecutive loss if trimethylamine) are characteristic for P-LPC.
- O-LPC has the least informative fragmentation pattern, with no specific fragment ions other than one at *m/z* 184, 104, and 86.
- Ratio of HG-derived ions (*m/z* 240, 184, and 104) can be used to differentiate between A-, O- and P-LPCs (Figure 1.3).

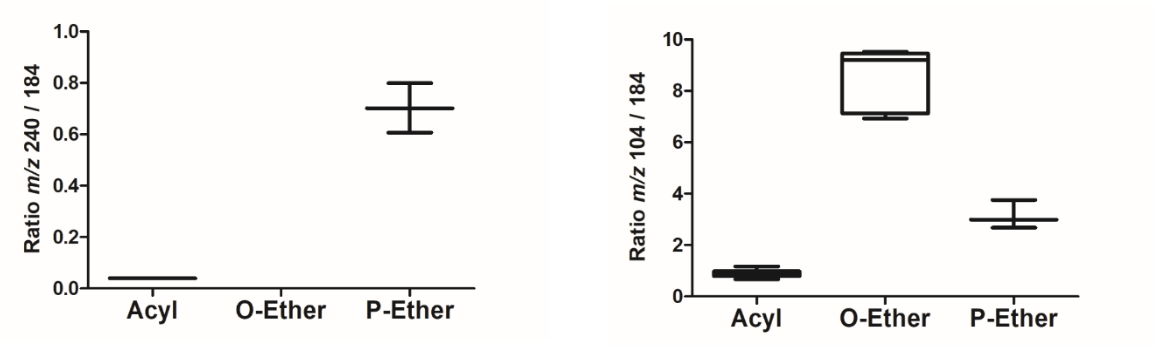


**Figure 1.3.** Fragment ions intensities ratio for 104/184 (left) and 240/184 (right) in the positive ion mode calculated using 16 LPC lipids identified in the study.

**Negative ion mode HCD:**

- Fatty acyl anion is characteristic only for A-LPC
- Fragment ions at *m/z* 224 (demethylated dehydroglycerol phosphocholine formed by the NL of carboxylic acid (A-LPC) or aldehyde (P-LPC)) and 168 (dimethylated ethylene phosphate) are characteristic for A- and P-LPC, but not O-LPC.

Observed fragmentation pattern and fragment ions intensity ratio are in agreement with previously published tandem MS studies of A-, O- and P-LPCs.[2,4]

##### 1.3. RT mapping

To establish retention time rules, molecular LPC species with single proposed structure (no possible isomers between A-LPC, O- and P-LPCs) were used. Here six lipid species representing A-LPC and O-LPC without double bounds in carbohydrate chains (marked in green in Table 1.2) were used as "RT anchors" which allowed to establish that Δ RT between A- and O-LPC pairs carrying the same carbohydrate chains correspond to 0.9-1.0 min.

For possible isomeric LPC with two proposed molecular species (e.g. LPC P-16:0 vs LPC O-16:1), Δ RT to corresponding A-LPC (e.g. LPC 16:0) was calculated, and if ΔRT was above 1.0 min or had a negative value, that ID was considered as a false positive (red values in the Table 1.2). Thus, ΔRT between A- and O-LPC and A- and P-LPC with the same carbon number were identified as 0.8-1.0 and 0.7-0.8 min, respectively.

**Table 1.2.** Calculated RT differences between A-, O- and P-LPC lipids used to identify true and false positive IDs.

| Bulk | [M+H]+ | RT_C18 | O-A | P-A | O-P |
|---|---|---|---|---|---|
| LPC 16:0 | 496.3397 | 5.97 | | | |
| LPC O-16:0 | 482.3605 | 6.88 | 0.91 | | 0.22 |
| LPC P-16:0 | 480.3448 | 6.66 | | 0.69 | |
| LPC O-16:1 | | | 2.35 | | |
| LPC 16:1 | 494.3241 | 4.31 | | | |
| LPC 18:0 | 524.371 | 8.16 | | | |
| LPC O-18:0 | 510.3918 | 9.17 | 1.01 | | 0.25 |
| LPC P-18:0 | 508.3761 | 8.92 | | 0.76 | |
| LPC O-18:1 | | | 2.64 | | |
| LPC 18:1 | 522.3554 | 6.28 | | | |
| LPC P-18:0 | 508.3761 | 7.1 | | -1.06 | |
| LPC O-18:1 | | | 0.82 | | 0.12 |
| LPC P-18:1 | 506.3605 | 6.98 | | 0.7 | |
| LPC O-18:2 | | | 2.09 | | |
| LPC 18:2 | 520.3397 | 4.89 | | | |
| LPC 20:0 | 552.4023 | 10.35 | | | |
| LPC 20:1 | 550.3867 | 8.29 | | | |
| LPC P-20:0 | 536.4074 | 9.13 | | -1.22 | |
| LPC O-20:1 | | | 0.84 | | |
| LPC 24:0 | 608.4649 | 14.18 | | | |
| LPC O-24:0 | 594.4857 | 15.08 | 0.9 | | |

Finally, structure – RT relationships for all identified LPC species were visualized by plotting Kendrick mass defect by hydrogen (KMD(H)) vs RT plot to control identification accuracy (Figure 1.4). All species falling out of the diagonal (different carbon number but the same DBE) and horizonal (same carbon number but different DBE) trendlines were excluded.

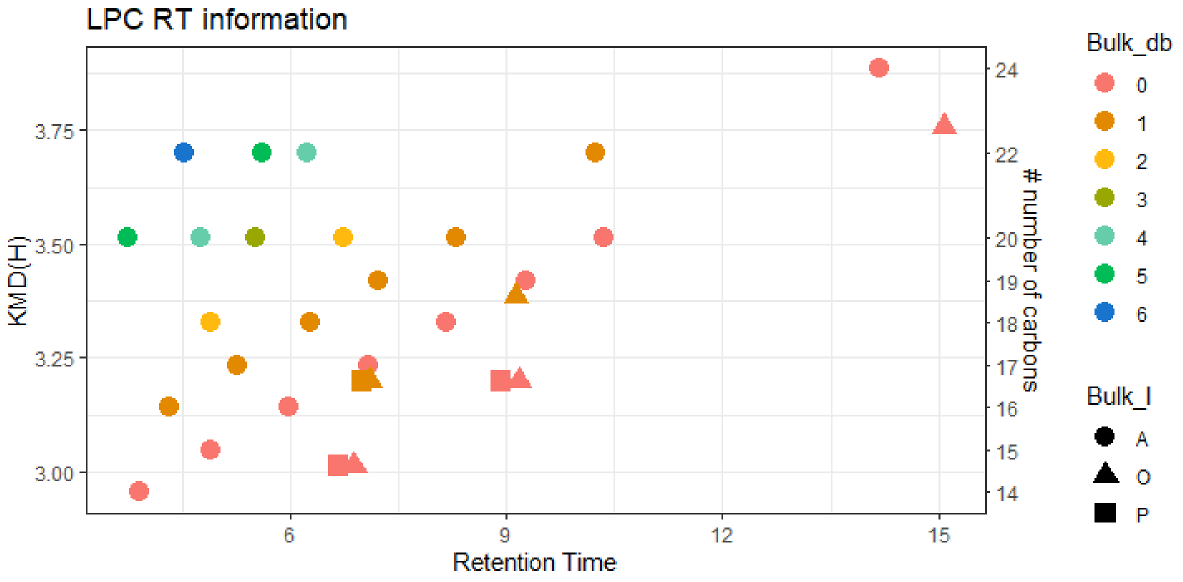


**Figure 1.4**. KMD(H), LPC carbohydrate chain carbon number vs RT plot for all identified LPC species. Symbols color represents the number of double bounds in carbohydrate chains. Circles – acyl LPC (A-LPC), triangles – alkyl LPC (O-LPC), and square – alkenyl ether LPC (P-LPC).

#### 2. Lysophosphatidylethanolamines (LPE)

Manual confirmation of identities for A-LPE, O-LPE and P-LPE (A- vs O- vs P-LPE) lipid molecular species was based on specific fragment ions, their ratio and retention time mapping.

##### 2.1. Monitored adducts

LPE were monitored as protonated or deprotonated ions in positive and negative ion modes, respectively.

##### 2.2. Fragmentation patterns

General fragmentation pattern and representative MS/MS spectra for LPE ionized in positive and negative modes are illustrated below:

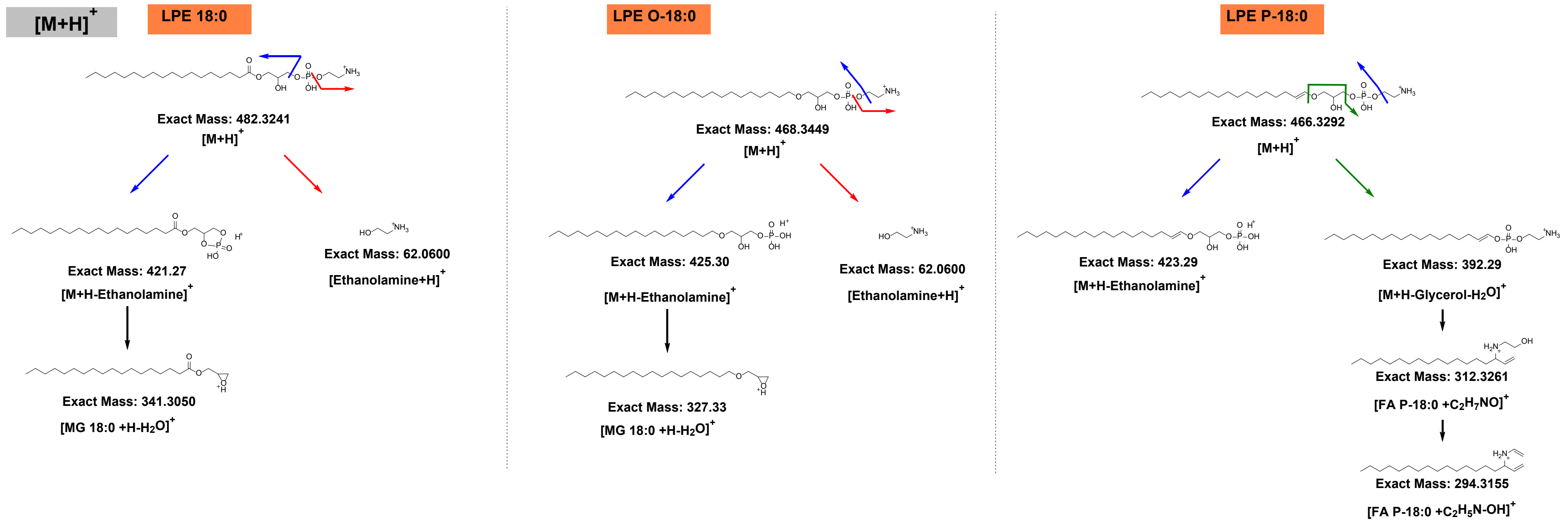


**Scheme 2.1**. General HCD fragmentation pattern of protonated acyl, alkyl and alkenyl linked LPE $[M+H]^+$ precursor ion.

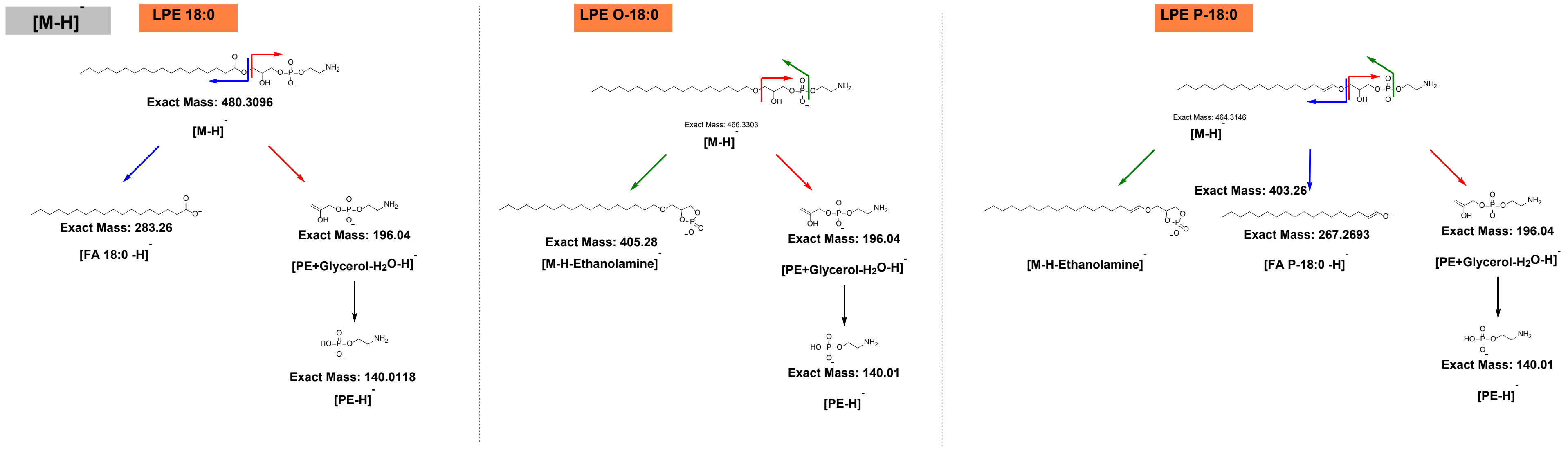


**Scheme 2.2**. General HCD fragmentation pattern of deprotonated acyl, alkyl and alkenyl linked LPE [M-H]$^-$ precursor ion.

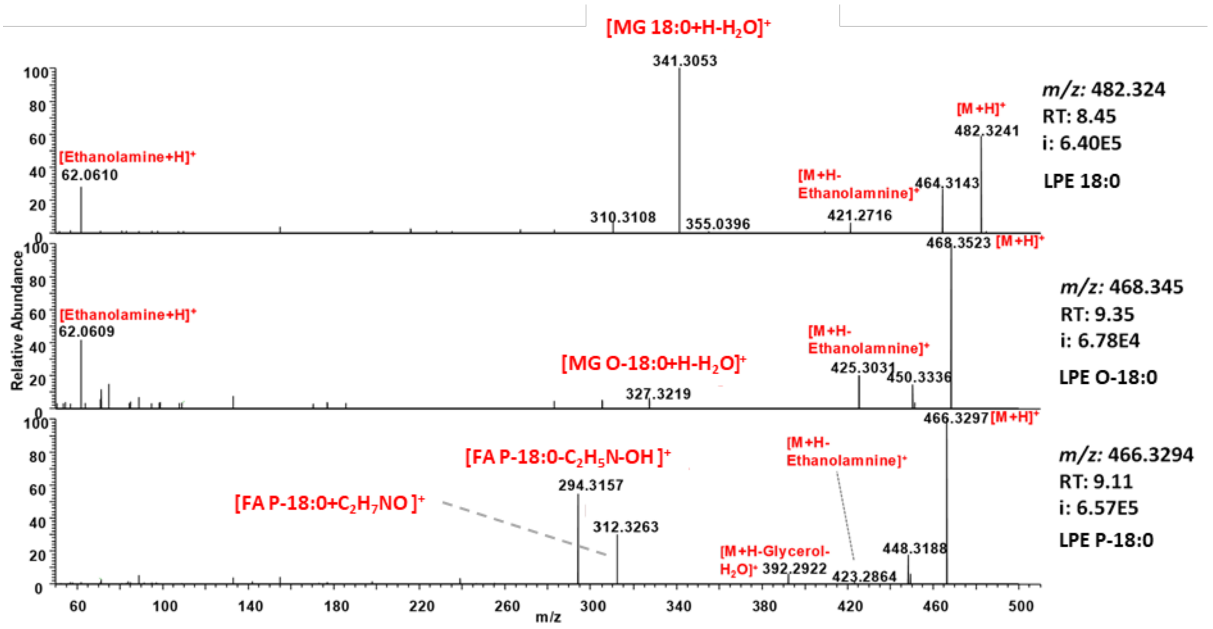


**Figure 2.1.** Representative HCD spectra of protonated LPE 18:0 (top), LPE O-18:0 (middle), and LPE P-18:0 (bottom).

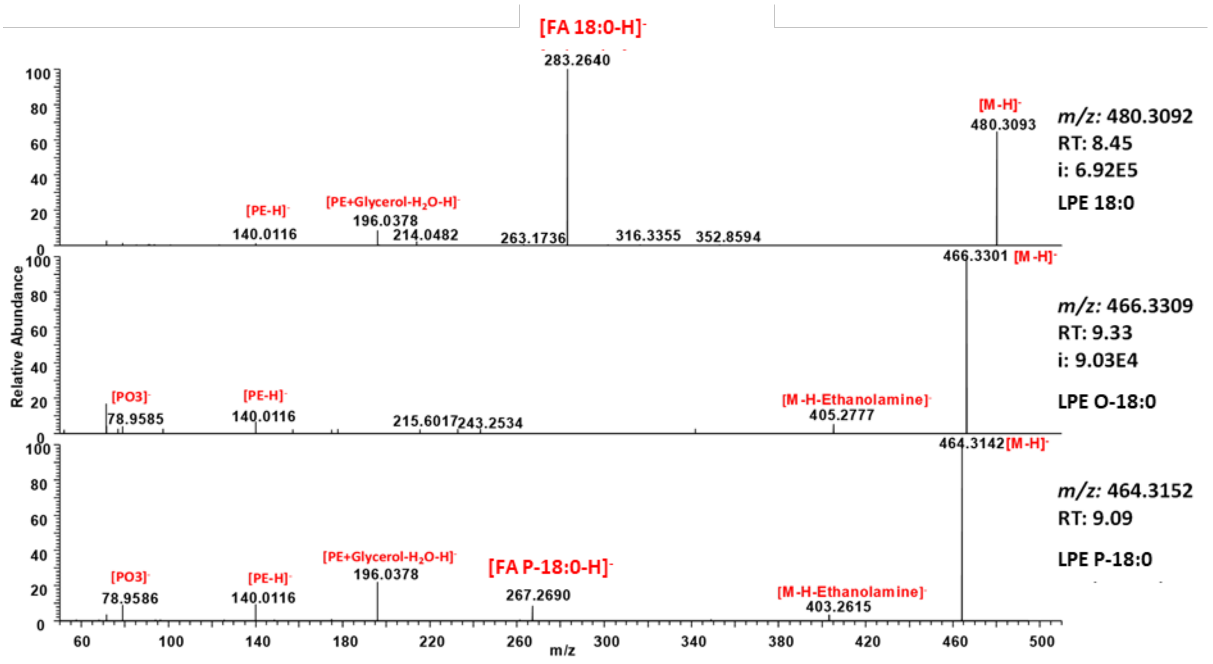


**Figure 2.2.** Representative HCD spectra of deprotonated LPE 18:0 (top), LPE O-18:0 (middle), and LPE P-18:0 (bottom).

**Table 2.1.** Summary of LPE specific neutral loss and fragment ions obtained by positive and negative ion modes HCD.

| NL/fragment ion | A-LPE | O-LPE | P-LPE |
|---|---|---|---|
| **Positive ion mode HCD** | | | |
| NL: -water (-18) | + | + | + |
| NL: -43 (ethanolamine) | | + | + |
| NL: -61 (ethanolamine + water) | + | | |
| NL: -141 (phosphoethanolamine) | + | + | |
| NL: -74 (dehydroglycerol) | | | + |
| NL: -154(dehydroglycerol phosphate) | | | + |
| NL: -172 (dehydroglycerol phosphate+water) | + | | + |
| FI: 62 (ethanolamine + water) | + | + | |
| **Negative ion mode HCD** | | | |
| NL: -61 (ethanolamine + water) | | + | + |
| FI: fatty acyl anion | + | | |
| FI: alkenyl anion | | | + |
| FI: 196 (dehydroglycerol phosphoethanolamine) | + | | + |
| FI: 140 (phosphoethanolamine) | + | | + |
| FI: 79.9 (metaphosphoric acid) | | + | + |

**Positive ion mode HCD:**

- NL of 141 (phosphoethanolamine) observed only for A- and O-LPE
- Consecutive NLs of -74 (dehydroglycerol), -154 (dehydroglycerol phosphate)172 (dehydroglycerol phosphate+water) are characteristic for P-LPE
- NL of -172 (dehydroglycerol phosphate+water) can be also detected for A-LPE (low intensity)

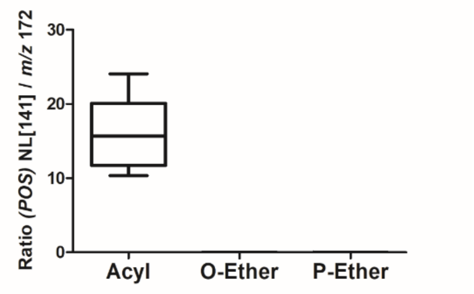


**Figure 2.3**. Fragment ions intensities ratio for NL[141]/172 in the positive ion mode calculated using 16 LPC lipids identified in the study.

**Negative ion mode HCD:**

- Fatty acyl anion is characteristic only for A-LPE
- Alkenyl anion is characteristic only for P-LPE

- The ratio of head group derived fragments intensities (196 to 140) can be used to differentiate between different LPE subclasses as illustrated at Figure 2.4.

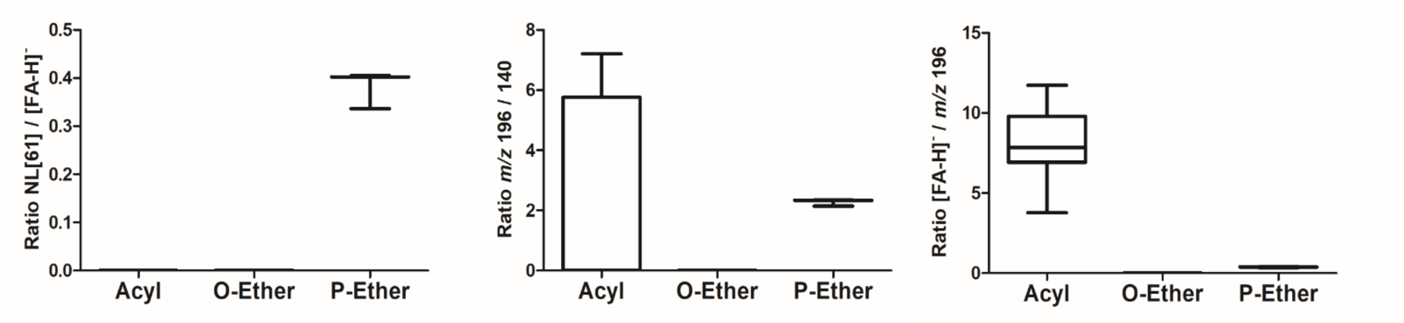


**Figure 2.4.** Ratio of fragment ions intensities (196 to 140) formed by negative ion mode HCD calculated using 12 LPE lipids identified in the study.

Observed fragmentation pattern and fragment ions intensities ratio are in agreement with previously published tandem MS studies of A-, O- and P-LPEs.[4]

##### 2.3. RT mapping

To establish retention time rules, molecular LPE species with single proposed structure (no possible isomers between A-LPE, O- and P-LPEs) were used. Here four lipid species representing A-LPE and O-LPE without double bounds in carbohydrate chains (marked in green in Table 1.2) were used as “RT anchors” which allowed to establish that Δ RT between A- and O-LPC pairs carrying the same carbohydrate chains correspond to 0.9-1.0 min.

For possible isomeric LPC with two proposed molecular species (e.g. LPE P-16:0 vs LPE O-16:1), Δ RT to corresponding A-LPC (e.g. LPC 16:0) was calculated, and if ΔRT was above 1.0 min or had a negative value, that ID was considered as a false positive (red values in the Table 1.2). Thus, ΔRT between A- and O-LPC and A- and P-LPC with the same carbon number were identified as 0.9-1.0 and 0.7-0.8 min, respectively.

**Table 2.2.** Calculated RT differences between A-, O- and P-LPC lipids used to identify true and false positive IDs.

| Bulk | [M+H]+ | RT_C18 | O-A | P-A | O-P |
|---|---|---|---|---|---|
| LPE 16:0 | 454.2928 | 6.11 | | | |
| LPE O-16:0 | 440.3135 | 7.04 | 0.93 | | 0.16 |
| LPE P-16:0 | 438.2979 | 6.88 | | 0.77 | |
| LPE O-16:1 | | 6.88 | | | |
| LPE 18:0 | 482.3241 | 8.36 | | | |
| LPE O-18:0 | 468.3448 | 9.34 | 0.98 | | 0.17 |
| LPE P-18:0 | 466.3292 | 9.17 | | 0.81 | |
| LPE O-18:1 | | 9.17 | **2.72** | | |
| LPE 18:1 | 480.3084 | 6.45 | | | |
| LPE P-18:1 | 464.3135 | 7.15 | | 0.7 | |
| LPE O-18:2 | | 7.15 | **2.16** | | |
| LPE 18:2 | 478.2928 | 4.99 | | | |

Finally, structure – RT relationships for all identified LPE species were visualized by plotting Kendrick mass defect by hydrogen (KMD(H)) vs RT plot to control identification accuracy (Figure 2.5). All species falling out of the diagonal (different carbon number but the same DBE) and horizonal (same carbon number but different DBE) trend lines were excluded.

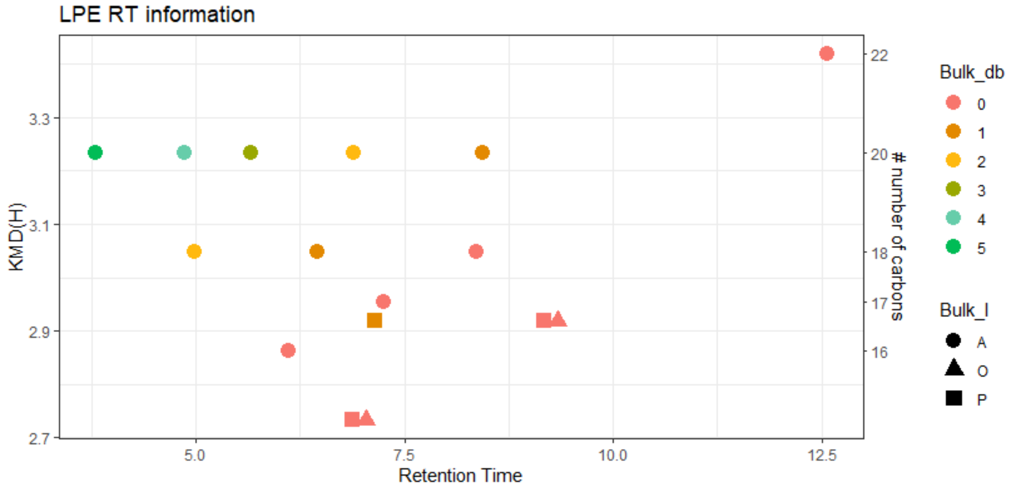


**Figure 2.5**. KMD(H), LPE carbohydrate chain carbon number vs RT plot for all identified LPE species. Symbols color represent the number of double bounds in carbohydrate chains. Circles – acyl LPE (A-LPE), triangles – alkyl LPE (O-LPE), and square – alkenyl ether LPE (P-LPE).

##### 3. Manual annotation of phosphatidylcholine (PC) lipids

Manual confirmation to assure accurate identification of PC subclasses (A- vs O- vs P-PC) based on specific fragment ions, their ratio and retention time mapping.

##### 3.1. Monitored adducts

PC were monitored as protonated or format adducts in positive and negative ion modes, respectively. $[M\text{-}CH_3]^-$ ions, usually formed as the result of in source fragmentation (ISF) by the collisional decomposition of anionic format adducts with the loss of neutral methyl format at the intermediate region of the instrument between the atmospheric pressure of the ESI source and the vacuum parts of the analyzer, were below 1% relative to the format adducts even for the most abundant LPC lipids and thus were not considered for the identification. However, $[M\text{-}CH_3]^-$ ions might be more prominent on the QTOF instruments operating with a higher pressure at the front end of the instrument or Orbitrap-based platforms with ion funnel configuration of the transmission devices [1–3].

##### 3.2. Fragmentation patterns

General fragmentation pattern and representative MS/MS spectra for PC ionized in positive and negative modes are illustrated below:

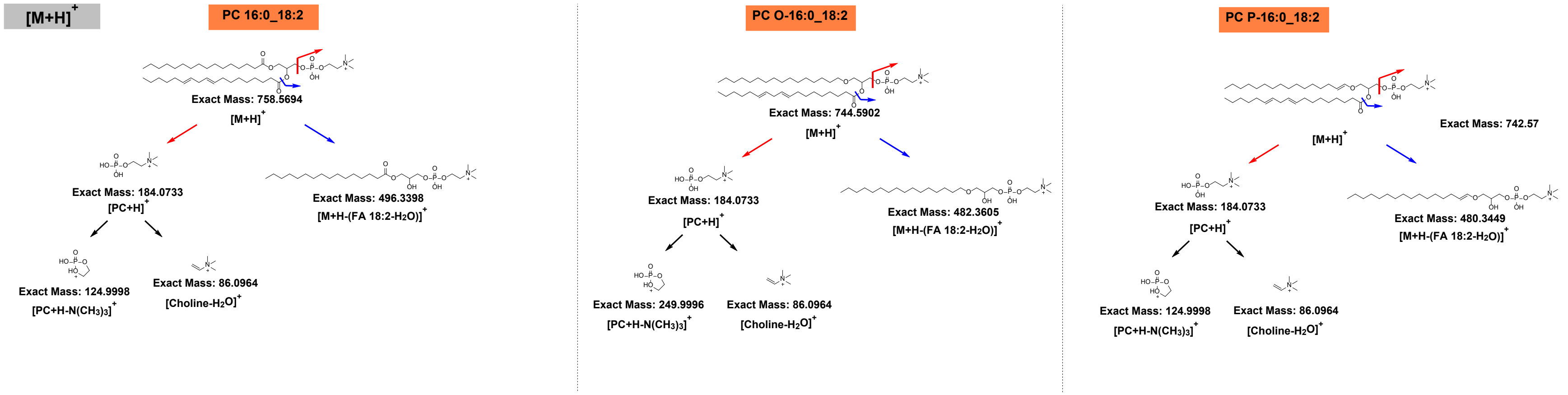


**Scheme 3.1**. General HCD fragmentation pattern of protonated PC $[M+H]^{+}$.

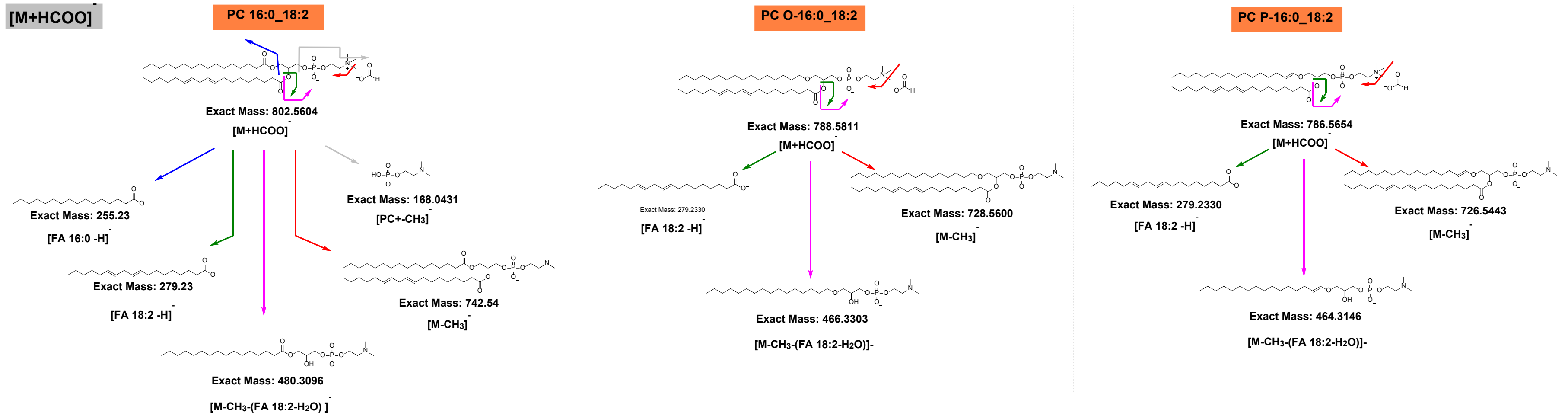


**Scheme 3.2.** General HCD fragmentation pattern of PC formate adduct $[M+HCOO]^{-}$.

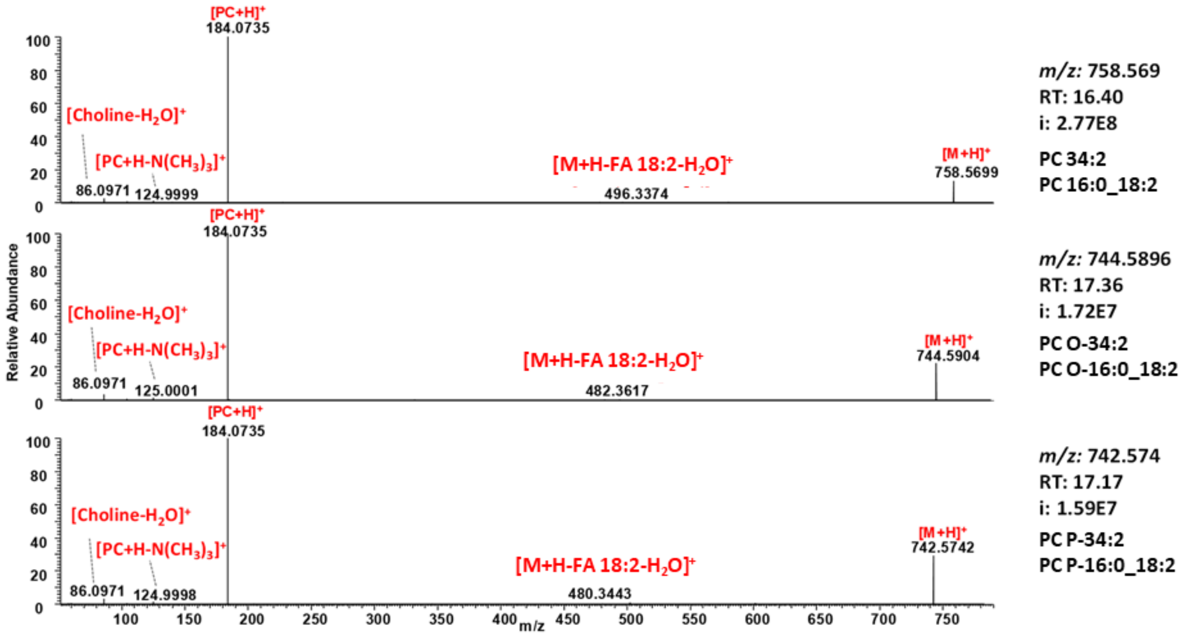


**Figure 3.1.** Representative HCD spectra of protonated PC 16:0_18:2 (top), PC O-16:0_18:2 (middle), and PC P-16:0_18:2 (bottom).

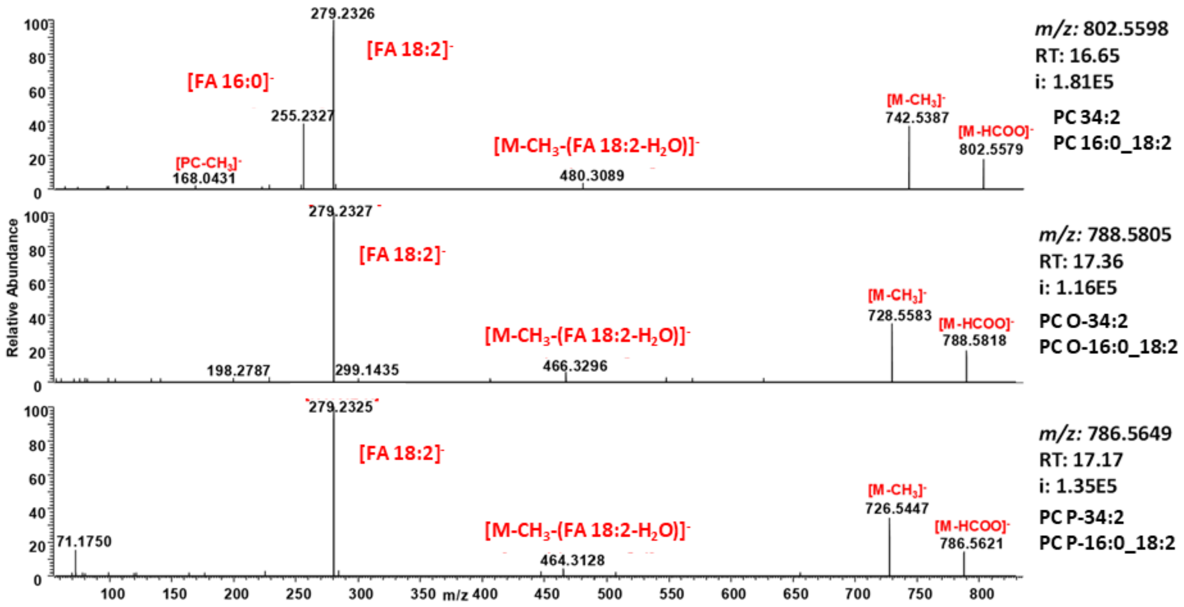


**Figure 3.2.** Representative HCD spectra of format adducts of PC 16:0_18:2 (top), PC O-16:0_18:2 (middle), and PC P-16:0_18:2 (bottom).

**Table 3.1.** Summary of PC specific neutral loss and fragment ions obtained by positive and negative ion modes HCD.

| NL/fragment ion | A-PC | O-PC | P-PC |
|---|---|---|---|
| **Positive ion mode HCD** | | | |
| NL: - (fatty acyl + water) | + | + | + |
| FI: 184 (phosphocholine) | + | + | + |
| FI: 125 (phosphocholine - trimethylamine) | + | + | + |
| FI: 86 (choline – water) | + | + | + |
| **Negative ion mode HCD** | | | |
| NL: - 60 (methyl format) | + | + | + |

| | | | |
|---|---|---|---|
| NL: - (fatty acyl + water + methyl) | + | + | + |
| FI: fatty acyl anion 1 | + | + | + |
| FI: fatty acyl anion 2 | + | | |
| FI: 224 (dehydroglycerol phosphocholine) | + | + | + |
| FI: 168 (demethylated phosphocholine) | + | + | + |

**Positive ion mode HCD:**

- All NLs (except second fatty acyl NL for A-PC, if present) and fragment ions were common for A-, O- and P-PC subclasses

**Negative ion mode HCD:**

- Presence of second fatty acyl anion is characteristic only for A-PC
- Other NL and fragment ions were common for A-, O- and P-PC subclasses

##### 3.3. RT mapping

To establish retention time rules for various PCs, molecular species with single proposed structure (no possible isomers between A-PC, O- and P-PCs) were used in the approach described for LPC and LPE lipids above (Table 3.2). ΔRT between A- and O-PC and A- and P-PC with the same carbon number were determined as 0.9 and 0.7 min, respectively, and were further used to filter out false positive identifications (shown in red).

**Table 3.2.** Calculated RT differences between A-, O- and P-PC lipids used to identify true and false positive IDs.

| Bulk | Discrete | [M+H]+ | [M+HCOO]- | RT_C18 | O-A | P-A | O-P |
|---|---|---|---|---|---|---|---|
| PC 30:0 | PC 14:0_16:0 | 706.5381 | 750.529 | 16.02 | | | |
| PC O-30:0 | PC O-16:0_14:0 | 692.5588 | 736.5497 | 16.94 | 0.92 | | 0.27 |
| PC P-30:0 | PC P-16:0_14:0 | 690.5432 | 734.5341 | 16.67 | | 0.65 | |
| PC O-30:1 | PC O-16:1_14:0 | | | | 2.15 | | |
| PC 30:1 | PC 14:0_16:1 | 704.5224 | 748.5134 | 14.52 | | | |
| PC 31:0 | PC 15:0_16:0 | 720.5537 | 764.5447 | 16.75 | | | |
| PC 31:1 | PC 15:0_16:1 | 718.5381 | 762.529 | 15.63 | | | |
| PC 32:0 | PC 16:0_16:0 | 734.5694 | 778.5603 | 17.42 | | | |
| PC O-32:0 | PC O-16:0_16:0 | 720.5901 | 764.581 | 18.35 | 0.93 | | 0.25 |
| PC P-32:0 | PC P-16:0_16:0 | 718.5745 | 762.5654 | 18.1 | | 0.68 | |
| PC O-32:1 | PC O-16:1_16:0 | | | | 2.01 | | |
| PC 32:1 | PC 16:0_16:1 | 732.5537 | 776.5447 | 16.09 | | | |
| PC O-32:1 | PC O-16:0_16:1 | 718.5745 | 762.5654 | 17 | 0.91 | | 0.22 |
| PC P-32:1 | PC P-16:0_16:1 | 716.5588 | 760.5497 | 16.78 | | 0.69 | |
| PC O-32:2 | PC O-16:1_16:1 | | | | 2.12 | | |
| PC P-32:1 | PC P-16:1_16:0 | 716.5588 | 760.5497 | 16.78 | | 0.69 | |
| PC 32:2 | PC 16:1_16:1 | 730.5381 | 774.529 | 14.66 | | | |
| PC 34:0 | PC 16:0_18:0 | 762.6007 | 806.5916 | 18.79 | | | |
| PC O-34:0 | PC O-18:0_16:0 | 748.6214 | 792.6123 | 19.68 | 0.89 | | |
| PC 34:1 | PC 16:0_18:1 | 760.585 | 804.576 | 17.46 | | | |
| PC O-34:1 | PC O-16:0_18:1 | 746.6058 | 790.5967 | 18.33 | 0.87 | | 0.22 |

| | | | | | | | |
|---|---|---|---|---|---|---|---|
| PC P-34:1 | PC P-18:1_16:0 | 746.6058 | 790.5967 | 18.33 | | -0.46 | |
| PC O-34:1 | PC O-18:1_16:0 | | | | 0.87 | | 0.22 |
| PC P-34:1 | PC P-16:0_18:1 | 744.5901 | 788.581 | 18.11 | | 0.65 | |
| PC O-34:2 | PC O-16:1_18:1 | | | | 1.97 | | |
| PC P-34:1 | PC P-16:1_18:0 | 744.5901 | 788.581 | 18.11 | | 0.65 | |
| PC O-34:2 | PC O-16:2_18:0 | | | | 1.97 | | |
| PC P-34:1 | PC P-18:0_16:1 | 744.5901 | 788.581 | 18.11 | | 0.65 | |
| PC O-34:2 | PC O-18:1_16:1 | | | | 1.97 | | |
| PC P-34:1 | PC P-18:1_16:0 | 744.5901 | 788.581 | 18.11 | | 0.65 | |
| PC O-34:2 | PC O-18:2_16:0 | | | | 1.74 | | |
| PC 34:2 | PC 16:1_18:1 | 758.5694 | 802.5603 | 16.14 | | | |
| PC 34:2 | PC 16:0_18:2 | 758.5694 | 802.5603 | 16.37 | | | |
| PC O-34:2 | PC O-16:0_18:2 | 744.5901 | 788.581 | 17.31 | 0.94 | | 0.27 |
| PC P-34:2 | PC P-16:0_18:2 | 742.5745 | 786.5654 | 17.04 | | 0.67 | |
| PC O-34:3 | PC O-16:1_18:2 | | | | 2.04 | | |
| PC 34:3 | PC 16:0_18:3 | 756.5537 | 800.5447 | 15.45 | | | |
| PC 34:3 | PC 16:1_18:2 | 756.5537 | 800.5447 | 15 | | | |
| PC P-34:3 | PC P-16:0_18:3 | 740.5588 | 784.5497 | 16.09 | | 0.64 | |
| PC O-34:4 | PC O-16:1_18:3 | | | | 1.46 | | |
| PC P-34:3 | PC P-16:1_18:2 | 740.5588 | 784.5497 | 15.68 | | 0.68 | |
| PC 34:4 | PC 14:0_20:4 | 754.5381 | 798.529 | 14.63 | | | |
| PC 35:2 | PC 17:0_18:2 | 772.585 | 816.576 | 17.16 | | | |
| PC P-35:2 | PC P-17:0_18:2 | 756.5901 | 800.581 | 17.74 | | 0.58 | |
| PC 36:1 | PC 18:0_18:1 | 788.6163 | 832.6073 | 18.79 | | | |
| PC O-36:1 | PC O-18:0_18:1 | 774.6371 | 818.628 | 19.64 | 0.85 | | 0.22 |
| PC P-36:1 | PC P-18:0_18:1 | 772.6214 | 816.6123 | 19.42 | | 0.63 | |
| PC O-36:2 | PC O-18:1_18:1 | | | | 1.95 | | |
| PC 36:2 | PC 16:0_20:2 | 786.6007 | 830.5916 | 17.47 | | | |
| PC P-36:2 | PC P-16:0_20:2 | 770.6058 | 814.5967 | 18.11 | | 0.64 | |
| PC O-36:3 | PC O-16:1_20:2 | | | | 1.36 | | |
| PC 36:2 | PC 18:0_18:2 | 786.6007 | 830.5916 | 17.81 | | | |
| PC O-36:2 | PC O-18:0_18:2 | 772.6214 | 816.6123 | 18.68 | 0.87 | | 0.26 |
| PC P-36:2 | PC P-18:0_18:2 | 770.6058 | 814.5967 | 18.42 | | 0.61 | |
| PC O-36:3 | PC O-18:1_18:2 | | | | 2 | | |
| PC 36:2 | PC 18:1_18:1 | 786.6007 | 830.5916 | 17.47 | | | |
| PC P-36:1 | PC P-18:0_18:1 | 772.6214 | 816.6123 | 18.26 | | -0.53 | |
| PC O-36:2 | PC O-18:1_18:1 | | | | 0.79 | | 0.15 |
| PC P-36:2 | PC P-18:1_18:1 | 770.6058 | 814.5967 | 18.11 | | 0.64 | |
| PC O-36:3 | PC O-18:2_18:1 | | | | 1.69 | | |
| PC 36:3 | PC 16:0_20:3 | 784.585 | 828.576 | 16.75 | | | |
| PC O-36:3 | PC O-16:0_20:3 | 770.6058 | 814.5967 | 17.63 | 0.88 | | 0.25 |
| PC P-36:3 | PC P-16:0_20:3 | 768.5901 | 812.581 | 17.38 | | 0.63 | |
| PC O-36:4 | PC O-16:1_20:3 | | | | 2.07 | | |
| PC 36:3 | PC 18:1_18:2 | 784.585 | 828.576 | 16.42 | | | |

| | | | | | | | |
|---|---|---|---|---|---|---|---|
| PC P-36:2 | PC P-18:0_18:2 | 770.6058 | 814.5967 | 17.24 | | -0.57 | |
| PC O-36:3 | PC O-18:1_18:2 | | | | 0.82 | | 0.18 |
| PC P-36:3 | PC P-18:1_18:2 | 768.5901 | 812.581 | 17.06 | | 0.64 | |
| PC O-36:4 | PC O-18:2_18:2 | | | | 1.6 | | |
| PC 36:4 | PC 16:0_20:4 | 782.5694 | 826.5603 | 16.17 | | | |
| PC O-36:4 | PC O-16:0_20:4 | 768.5901 | 812.581 | 17.06 | 0.89 | | 0.27 |
| PC P-36:4 | PC P-16:0_20:4 | 766.5745 | 810.5654 | 16.79 | | 0.62 | |
| PC O-36:5 | PC O-16:1_20:4 | | | | 2.01 | | |
| PC 36:4 | PC 18:1_18:3 | 782.5694 | 826.5603 | 15.46 | | | |
| PC P-36:4 | PC P-18:1_18:3 | 766.5745 | 810.5654 | 16.13 | | 0.67 | |
| PC O-36:5 | PC O-18:2_18:3 | | | | 1.35 | | |
| PC 36:4 | PC 18:2_18:2 | 782.5694 | 826.5603 | 15.46 | | | |
| PC 36:4 | PC 16:1_20:3 | 782.5694 | 826.5603 | 15.31 | | | |
| PC 36:5 | PC 16:0_20:5 | 780.5537 | 824.5447 | 15.26 | | | |
| PC O-36:5 | PC O-16:0_20:5 | 766.5745 | 810.5654 | 16.12 | 0.86 | | 0.25 |
| PC P-36:5 | PC P-16:0_20:5 | 764.5588 | 808.5497 | 15.87 | | 0.61 | |
| PC O-36:6 | PC O-16:1_20:5 | | | | | | |
| PC 36:5 | PC 16:1_20:4 | 780.5537 | 824.5447 | 14.78 | | | |
| PC P-36:5 | PC P-16:1_20:4 | 764.5588 | 808.5497 | 15.41 | | 0.63 | |
| PC 37:4 | PC 17:0_20:4 | 796.585 | 840.576 | 16.92 | | | |
| PC O-37:4 | PC O-17:0_20:4 | 782.605 | 826.5967 | 17.73 | 0.81 | | |
| PC 38:3 | PC 18:0_20:3 | 812.6163 | 856.6073 | 18.15 | | | |
| PC O-38:3 | PC O-18:0_20:3 | 798.6371 | 842.628 | 18.96 | 0.81 | | |
| PC P-38:3 | PC P-18:0_20:3 | 796.6214 | 840.6123 | 18.71 | | 0.56 | |
| PC O-38:4 | PC O-18:1_20:3 | | | | 1.92 | | |
| PC 38:4 | PC 16:0_22:4 | 810.6007 | 854.5916 | 17.16 | | | |
| PC P-38:4 | PC P-16:0_22:4 | 794.6058 | 838.5967 | 17.74 | | 0.58 | |
| PC O-38:5 | PC O-16:1_22:4 | | | | 1.28 | | |
| PC 38:4 | PC 18:0_20:4 | 810.6007 | 854.5916 | 17.6 | | | |
| PC O-38:4 | PC O-18:0_20:4 | 796.6214 | 840.6123 | 18.42 | 0.82 | | 0.23 |
| PC P-38:4 | PC P-18:0_20:4 | 794.6058 | 838.5967 | 18.19 | | 0.59 | |
| PC O-38:5 | PC O-18:1_20:4 | | | | 1.97 | | |
| PC 38:4 | PC 18:1_20:3 | 810.6007 | 854.5916 | 16.79 | | | |
| PC P-38:3 | PC P-18:0_20:3 | 796.6214 | 840.6123 | 17.56 | | -0.59 | |
| PC O-38:4 | PC O-18:1_20:3 | | | | 0.77 | | |
| PC 38:5 | PC 16:0_22:5 | 808.585 | 852.576 | 16.46 | | | |
| PC P-38:5 | PC P-16:0_22:5 | 792.5901 | 836.581 | 17.1 | | 0.64 | |
| PC O-38:6 | PC O-16:1_22:5 | | | | 1.3 | | |
| PC 38:5 | PC 18:1_20:4 | 808.585 | 852.576 | 16.22 | | | |
| PC P-38:4 | PC P-18:0_20:4 | 794.6058 | 838.5967 | 17 | | -0.6 | |
| PC O-38:5 | PC O-18:1_20:4 | | | | 0.78 | | |
| PC P-38:5 | PC P-18:1_20:4 | 792.5901 | 836.581 | 16.83 | | 0.61 | |
| PC O-38:6 | PC O-18:2_20:4 | | | | 1.76 | | |
| PC 38:5 | PC 18:0_20:5 | 808.585 | 852.576 | 16.71 | | | |

| | | | | | | | |
|---|---|---|---|---|---|---|---|
| **PC P-38:5** | **PC P-18:0_20:5** | **792.5901** | **836.581** | **17.3** | | **0.59** | |
| **PC O-38:6** | **PC O-18:1_20:5** | | | | **2.02** | | |
| **PC 38:6** | **PC 16:0_22:6** | 806.5694 | 850.5603 | 15.8 | | | |
| **PC O-38:6** | **PC O-16:0_22:6** | 792.5901 | 836.581 | 16.83 | **1.03** | | **0.43** |
| **PC P-38:6** | **PC P-16:0_22:6** | **790.5745** | **834.5654** | **16.4** | | **0.6** | |
| **PC O-38:7** | **PC O-16:1_22:6** | | | | | | |
| **PC 38:6** | **PC 18:1_20:5** | 806.5694 | 850.5603 | 15.28 | | | |
| **PC P-38:6** | **PC P-18:1_20:5** | **790.5745** | **834.5654** | **15.89** | | **0.61** | |
| **PC O-38:7** | **PC O-18:2_20:5** | | | | | | |
| **PC 38:6** | **PC 18:2_20:4** | 806.5694 | 850.5603 | 15.07 | | | |
| **PC P-38:6** | **PC P-18:2_20:4** | **790.5745** | **834.5654** | **15.7** | | **0.63** | |
| **PC O-38:7** | **PC O-18:3_20:4** | | | | | | |

Finally, structure – RT relationships for all identified LPE species were visualized by plotting Kendrick mass defect by hydrogen (KMD(H)) vs RT plot to control identification accuracy (Figure 3.3 and 3.4). All species falling out of the diagonal (different carbon number but the same DBE) and horizonal (same carbon number but different DBE) trend lines were excluded. To simplify the visualization, A-PC were plotted separately form O-/P-linked PC lipids.

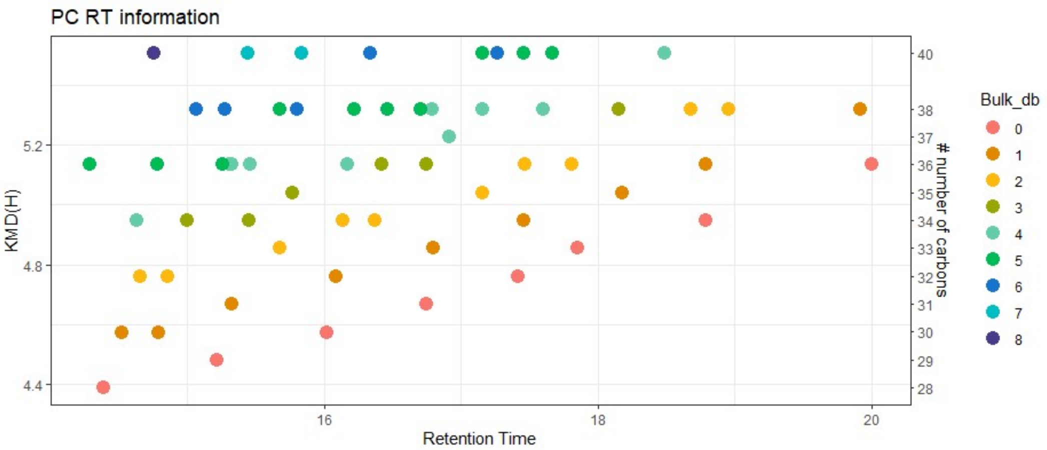


**Figure 3.3**. KMD(H), A-PC carbohydrate chain carbon number vs RT plot for all identified A-PC species. Symbols color represents the number of double bounds in carbohydrate chains.

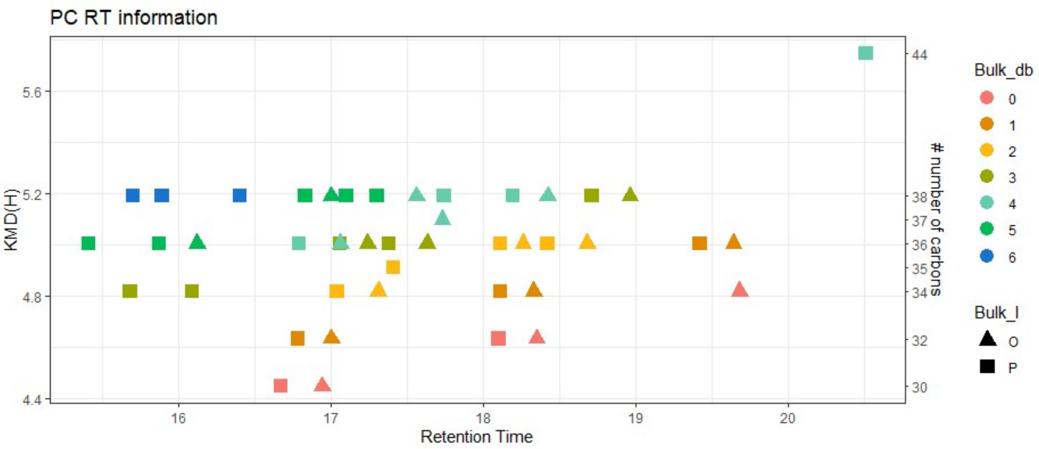

**Figure 3.4**. KMD(H), PC carbohydrate chain carbon number vs RT plot for all identified O-/P-PC species. Symbols color represents the number of double bounds in carbohydrate chains. Triangles – alkyl PC (O-LPC), and square – alkenyl ether PC (P-LPC).

#### 4. Manual annotation of phosphatidylethanolamine (PE) lipids

Manual confirmation to assure accurate identification of PE subclasses (A- vs O- vs P-PE) was based on specific fragment ions, their ratio and retention time mapping.

##### 4.1. Monitored adducts

PE were monitored as protonated or deprotonated ions in positive and negative ion modes, respectively.

##### 4.2. Fragmentation patterns

General fragmentation pattern and representative MS/MS spectra for PE ionized in positive and negative modes are illustrated below:

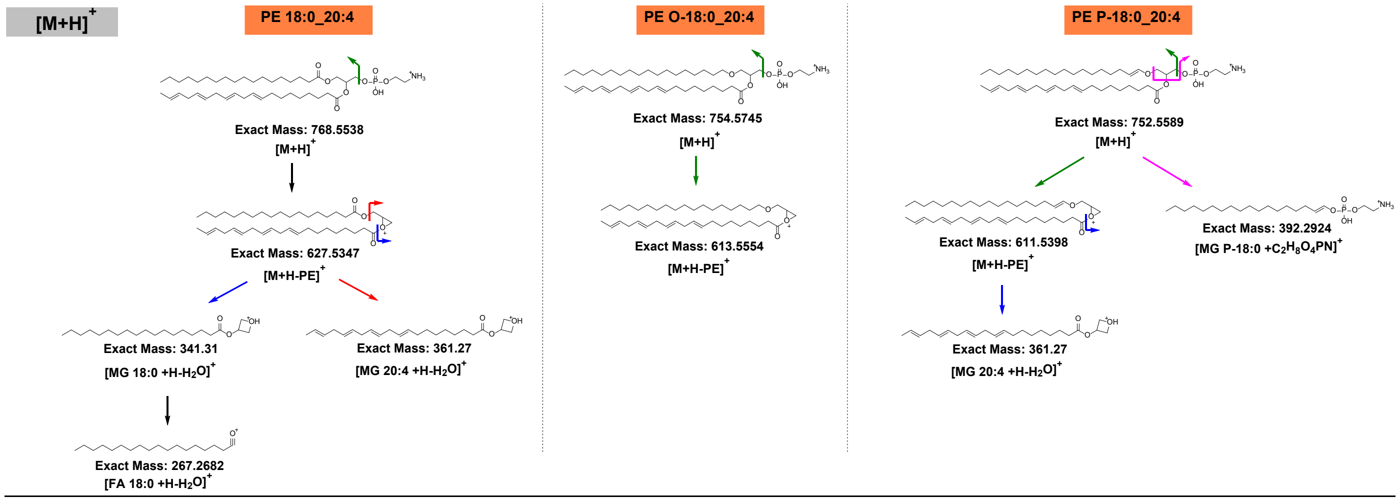


**Scheme 4.1.** General HCD fragmentation pattern of protonated acyl-, alkyl and alkenyl-chain PE [M+H]+.

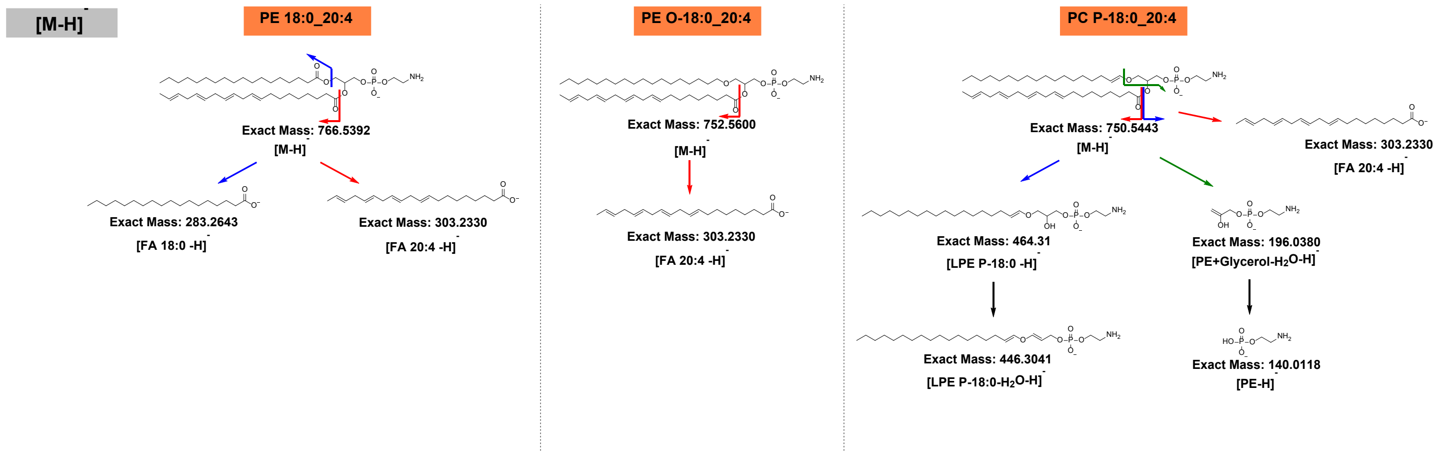


**Scheme 4.2.** General HCD fragmentation pattern of deprotonated acyl-, alkyl and alkenyl-chain PE [M-H]-

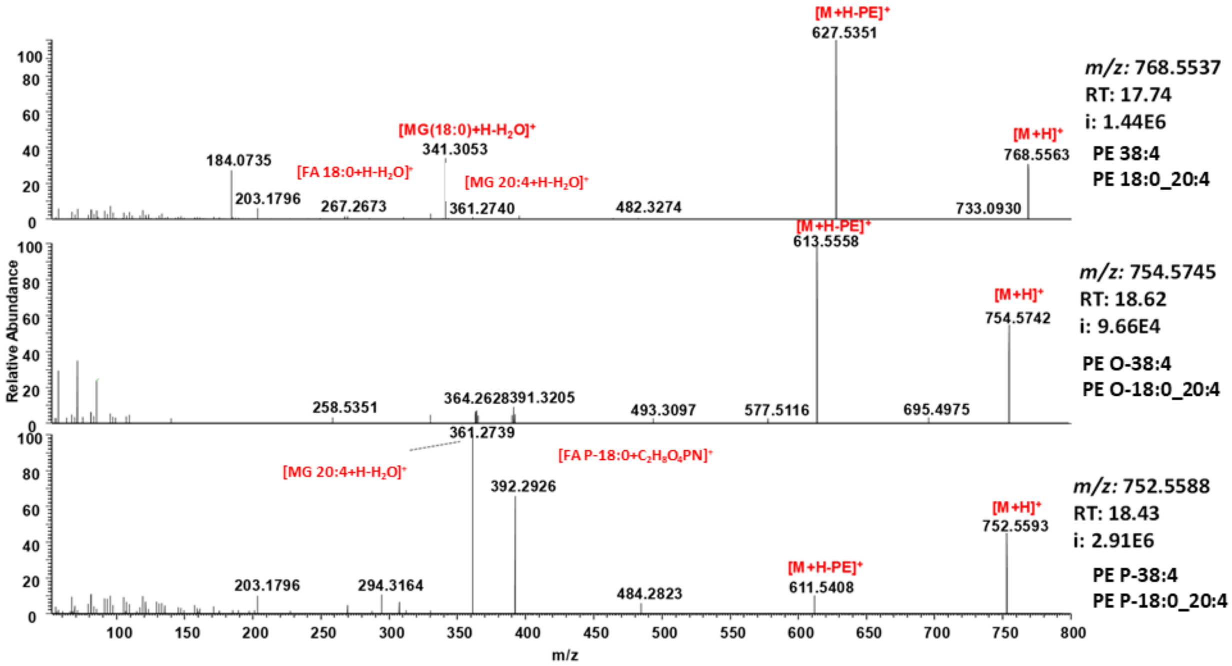


**Figure 4.1.** Representative HCD spectra of protonated adduct $[M+H]^+$ of PE 18:0_20:4 (top), PE O-18:0_20:4 (middle) and PE P-18:0_20:4 (bottom).

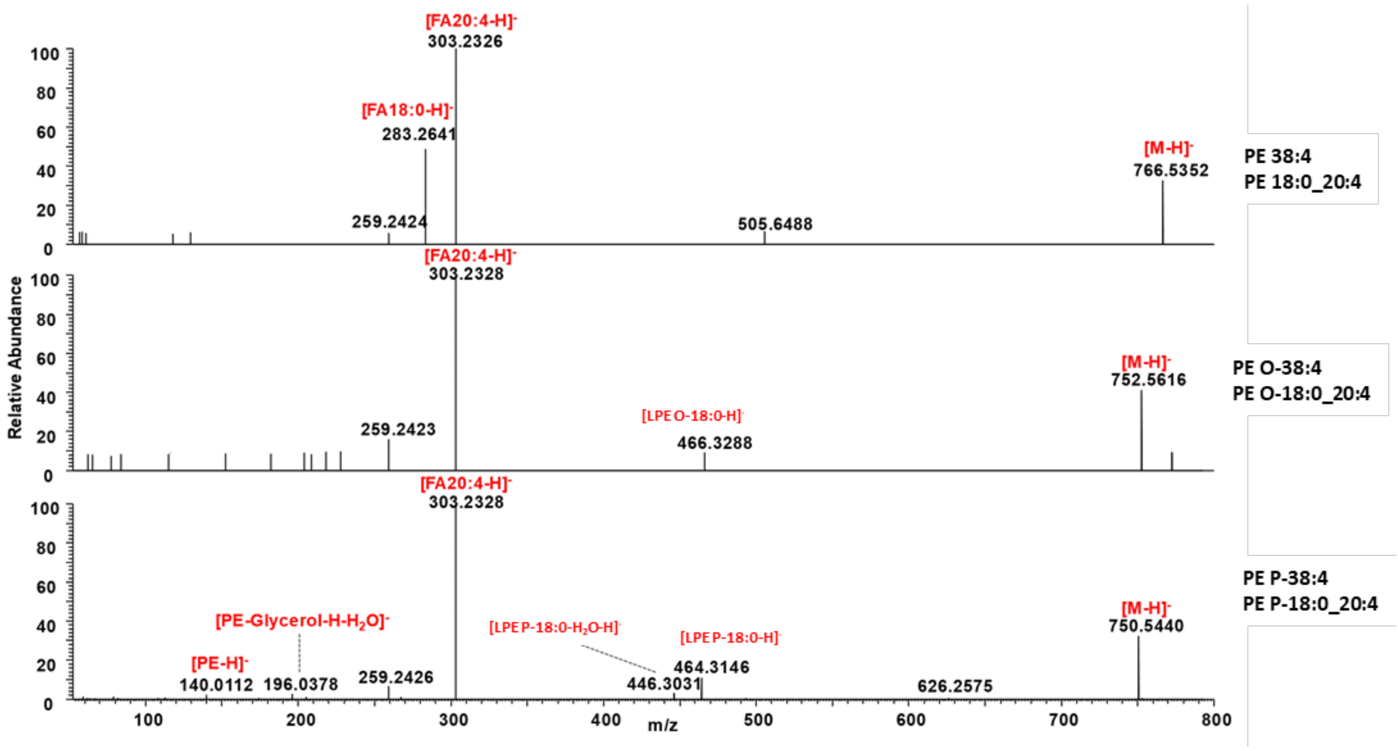


**Figure 4.2.** Representative HCD spectra of deprotonated adducts $[M-H]^-$ of PE 18:0_20:4 (top), PE O-18:0_20:4 (middle) and PE P-18:0_20:4 (bottom).

**Table 4.1.** Summary of PE specific neutral loss and fragment ions obtained by positive and negative ion modes HCD.

| NL/fragment ion | A-PE | O-PE | P-PE |
|---|---|---|---|
| **Positive ion mode HCD** | | | |
| NL: -141 (phosphoethanolamine) | + | + | + |
| FI: acyl chain 1 specific | + | | + |
| FI: acyl chain 2 specific | + | | |
| FI: alkenyl chain specific | | | + |
| **Negative ion mode HCD** | | | |
| FI: O-/P-LPE | | + | + |
| FI: fatty acyl 1 anion | + | + | + |
| FI: fatty acyl 2 anion | + | | |
| FI: 196 (dehydroglycerol phosphatidylethanolamine) | + | + | + |
| FI: 140 (phosphatidylethanolamine) | + | + | + |

**Positive ion mode HCD:**

- NL of 141 (phosphoethanolamine) observed only for A- and P-PE
- High ratio of MAG[FA1-$H_2O$]$^+$ and MAG[FA2-$H_2O$]$^+$ to NL [141] is highly diagnostic for P-PE

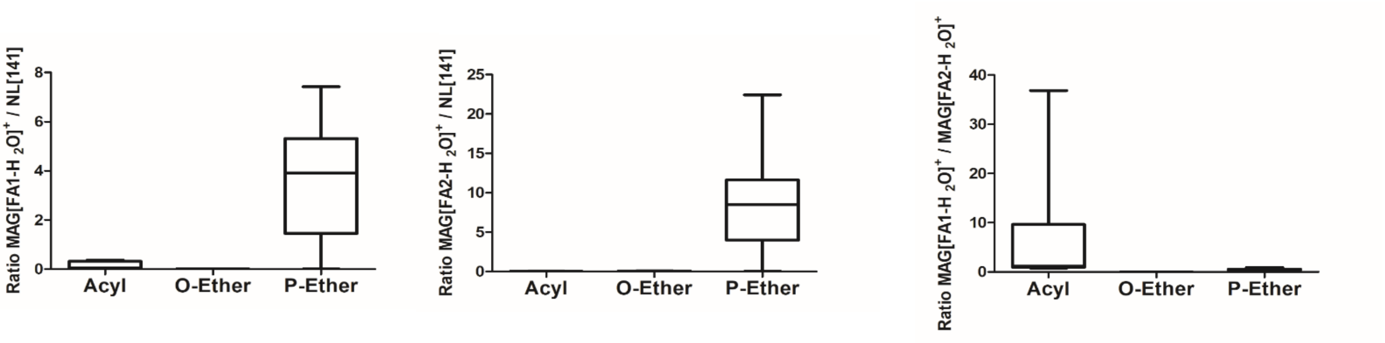


**Figure 4.3.** Ratio of fragment ions intensities formed by positive ion mode HCD calculated using PE lipids identified in the study.

Observed fragmentation pattern and fragment ions intensities ratio are in agreement with previously published tandem MS studies of A-, O- and P-PEs.[4]

##### 4.3. RT mapping

To establish retention time rules for various PEs, molecular species with single proposed structure (no possible isomers between A-PE, O- and P-PEs) were used in the approach described for PC and PE lipids above (Table 4.2). ΔRT between A- and O-PE and A- and P-PE with the same carbon number were determined as 0.9 and 0.7 min, respectively, and were further used to filter out false positive identifications (shown in red).

**Table 4.2.** Calculated RT differences between A-, O- and P-PE lipids used to identify true and false positive IDs.

| Bulk | Discrete | [M+H]+ | [M-H]- | RT_Leipzig | O-A | P-A | O-P |
|---|---|---|---|---|---|---|---|
| PE 32:0 | PE 16:0_16:0 | 692.5224 | 690.5079 | 17.65 | | | |
| PE P-32:0 | PE P-16:0_16:0 | 676.5275 | 674.513 | 18.33 | | 0.68 | |
| PE O-32:1 | PE O-16:1_16:0 | | | | 1.99 | | |
| PE 32:1 | PE 16:0_16:1 | 690.5068 | 688.4922 | 16.34 | | | |

| | | | | | | | |
|---|---|---|---|---|---|---|---|
| PE P-32:1 | PE P-16:0_16:1 | 674.5119 | 672.4973 | 17.04 | | 0.7 | |
| PE O-32:1 | PE O-16:1_16:1 | | | | 2.14 | | |
| PE 32:2 | PE 16:1_16:1 | 688.4911 | 686.4766 | 14.9 | | | |
| PE P-33:1 | PE P-15:0_18:1 | 688.5275 | 686.513 | 17.7 | | | |
| PE P-33:1 | PE P-16:0_17:1 | 688.5275 | 686.513 | 17.7 | | | |
| PE P-33:1 | PE P-17:0_16:1 | 688.5275 | 686.513 | 17.7 | | | |
| PE P-33:2 | PE P-15:0_18:2 | 686.5119 | 684.4973 | 16.56 | | | |
| PE 34:0 | PE 16:0_18:0 | 720.5537 | 718.5392 | 18.95 | | | |
| PE P-34:0 | PE P-16:0_18:0 | 704.5588 | 702.5443 | 19.54 | | 0.59 | |
| PE O-34:1 | PE O-16:1_18:0 | | | | 1.86 | | |
| PE P-34:0 | PE P-18:0_16:0 | 704.5588 | 702.5443 | 19.54 | | 0.59 | |
| PE O-34:1 | PE O-18:1_16:0 | | | | 1.86 | | |
| PE 34:1 | PE 16:0_18:1 | 718.5381 | 716.5235 | 17.68 | | | |
| PE O-34:1 | PE O-16:0_18:1 | 704.5588 | 702.5443 | 18.53 | 0.85 | | 0.23 |
| PE P-34:1 | PE P-16:0_18:1 | 702.5432 | 700.5286 | 18.3 | | 0.62 | |
| PE O-34:2 | PE O-16:1_18:1 | | | | 1.93 | | |
| PE P-34:1 | PE P-18:1_16:0 | 702.5432 | 700.5286 | 18.3 | | 0.62 | |
| PE O-34:2 | PE O-18:2_16:0 | | | | 1.66 | | |
| PE 34:1 | PE 16:1_18:0 | 718.5381 | 716.5235 | 17.68 | | | |
| PE P-34:1 | PE P-18:0_16:1 | 702.5432 | 700.5286 | 18.3 | | 0.62 | |
| PE O-34:2 | PE O-18:1_16:1 | | | | 1.93 | | |
| PE 34:2 | PE 16:0_18:2 | 716.5224 | 714.5079 | 16.64 | | | |
| PE P-34:2 | PE P-16:0_18:2 | 700.5275 | 698.513 | 17.29 | | 0.65 | |
| PE O-34:3 | PE O-16:1_18:2 | | | | 2.04 | | |
| PE 34:2 | PE 16:1_18:1 | 716.5224 | 714.5079 | 16.37 | | | |
| PE P-34:2 | PE P-18:1_16:1 | 700.5275 | 698.513 | 17.04 | | 0.67 | |
| PE O-34:3 | PE O-18:2_16:1 | | | | 1.79 | | |
| PE 34:3 | PE 16:0_18:3 | 714.5068 | 712.4922 | 15.68 | | | |
| PE P-34:3 | PE P-16:0_18:3 | 698.5119 | 696.4973 | 16.35 | | 0.67 | |
| PE O-34:4 | PE O-16:1_18:3 | | | | 1.45 | | |
| PE 34:3 | PE 16:1_18:2 | 714.5068 | 712.4922 | 15.25 | | | |
| PE P-34:3 | PE P-16:1_18:2 | 698.5119 | 696.4973 | 15.94 | | 0.69 | |
| PE 34:4 | PE 14:0_20:4 | 712.4911 | 710.4766 | 14.9 | | | |
| PE P-34:4 | PE P-14:0_20:4 | 696.4962 | 694.4817 | 15.55 | | 0.65 | |
| PE 35:1 | PE 17:0_18:1 | 732.5537 | 730.5392 | 18.33 | | | |
| PE P-35:1 | PE P-17:0_18:1 | 716.5588 | 714.5443 | 18.95 | | 0.62 | |
| PE P-35:1 | PE P-18:1_17:0 | 716.5588 | 714.5443 | 18.68 | | | |
| PE O-35:2 | PE O-18:2_17:0 | | | | 1.34 | | |
| PE P-35:1 | PE P-16:0_19:1 | 716.5588 | 714.5443 | 18.95 | | 0.62 | |
| PE O-35:2 | PE O-16:1_19:1 | | | | 1.61 | | |
| PE 35:2 | PE 17:0_18:2 | 730.5381 | 728.5235 | 17.34 | | | |
| PE P-35:2 | PE P-17:0_18:2 | 714.5432 | 712.5286 | 17.95 | | 0.61 | |
| PE P-35:2 | PE P-18:1_17:1 | 714.5432 | 712.5286 | 17.65 | | | |
| PE O-35:3 | PE O-18:2_17:1 | | | | | | |

| | | | | | | | |
|---|---|---|---|---|---|---|---|
| PE 36:1 | PE 16:0_20:1 | 746.5694 | 744.5548 | 18.94 | | | |
| PE P-36:1 | PE P-16:0_20:1 | 730.5745 | 728.5599 | 19.39 | | 0.45 | |
| PE O-36:2 | PE O-16:1_20:1 | | | | 1.69 | | |
| PE P-36:1 | PE P-20:1_16:0 | 730.5745 | 728.5599 | 19.39 | | | |
| PE O-36:2 | PE O-20:2_16:0 | | | | 1.69 | | |
| PE 36:1 | PE 18:0_18:1 | 746.5694 | 744.5548 | 18.94 | | | |
| PE O-36:1 | PE O-18:0_18:1 | 732.5901 | 730.5756 | 19.69 | 0.75 | | 0.16 |
| PE P-36:1 | PE P-18:0_18:1 | 730.5745 | 728.5599 | 19.53 | | 0.59 | |
| PE O-36:2 | PE O-18:1_18:1 | | | | 1.83 | | |
| PE 36:2 | PE 18:0_18:2 | 744.5537 | 742.5392 | 18 | | | |
| PE P-36:2 | PE P-18:0_18:2 | 728.5588 | 726.5443 | 18.6 | | 0.6 | |
| PE O-36:3 | PE O-18:1_18:2 | | | | 1.96 | | |
| PE 36:2 | PE 18:1_18:1 | 744.5537 | 742.5392 | 17.7 | | | |
| PE P-36:2 | PE P-18:1_18:1 | 728.5588 | 726.5443 | 18.3 | | 0.6 | |
| PE O-36:3 | PE O-18:2_18:1 | | | | 1.66 | | |
| PE P-36:2 | PE P-16:0_20:2 | 728.5588 | 726.5443 | 18.3 | | | |
| PE O-36:3 | PE O-16:1_20:2 | | | | 1.34 | | |
| PE 36:3 | PE 16:0_20:3 | 742.5381 | 740.5235 | 16.96 | | | |
| PE P-36:3 | PE P-16:0_20:3 | 726.5432 | 724.5286 | 17.6 | | 0.64 | |
| PE O-36:4 | PE O-16:1_20:3 | | | | 1.18 | | |
| PE 36:3 | PE 18:0_18:3 | 742.5381 | 740.5235 | 17.13 | | | |
| PE P-36:3 | PE P-18:0_18:3 | 726.5432 | 724.5286 | 17.77 | | 0.64 | |
| PE O-36:4 | PE O-18:1_18:3 | | | | 2.22 | | |
| PE 36:3 | PE 18:1_18:2 | 742.5381 | 740.5235 | 16.64 | | | |
| PE P-36:3 | PE P-18:1_18:2 | 726.5432 | 724.5286 | 17.31 | | 0.67 | |
| PE O-36:4 | PE O-18:2_18:2 | | | | 1.76 | | |
| PE 36:4 | PE 16:0_20:4 | 740.5224 | 738.5079 | 16.42 | | | |
| PE P-36:4 | PE P-16:0_20:4 | 724.5275 | 722.513 | 17.04 | | 0.62 | |
| PE O-36:5 | PE O-16:1_20:4 | | | | 2.04 | | |
| PE 36:4 | PE 18:2_18:2 | 740.5224 | 738.5079 | 15.55 | | | |
| PE P-36:4 | PE P-18:1_18:3 | 724.5275 | 722.5541 | 16.7 | | | |
| PE O-36:5 | PE O-18:2_18:3 | | | | 1.7 | | |
| PE 36:5 | PE 16:0_20:5 | 738.5068 | 736.4922 | 15.52 | | | |
| PE P-36:5 | PE P-16:0_20:5 | 722.5119 | 720.4973 | 16.15 | | 0.63 | |
| PE O-36:6 | PE O-16:1_20:5 | | | | | | |
| PE 36:5 | PE 16:1_20:4 | 738.5068 | 736.4922 | 15 | | | |
| PE P-36:5 | PE P-16:1_20:4 | 722.5119 | 720.4973 | 15.68 | | 0.68 | |
| PE P-37:3 | PE P-17:0_20:3 | 740.5588 | 738.5443 | 18.26 | | | |
| PE 37:4 | PE 17:0_20:4 | 754.5381 | 752.5235 | 17.13 | | | |
| PE P-37:4 | PE P-17:0_20:4 | 738.5432 | 736.5286 | 17.74 | | 0.61 | |
| PE 37:5 | PE 17:1_20:4 | 752.5224 | 750.5079 | 15.81 | | | |
| PE P-37:5 | PE P-17:0_20:5 | 736.5275 | 734.513 | 16.6 | | 0.79 | |
| PE 38:1 | PE 18:0_20:1 | 774.6007 | 772.5861 | 19.96 | | | |
| PE P-38:1 | PE P-18:0_20:1 | 758.6058 | 756.5912 | 20.59 | | 0.63 | |

| | | | | | | | |
|---|---|---|---|---|---|---|---|
| PE O-38:2 | PE O-18:1_20:1 | | | | 1.79 | | |
| PE 38:2 | PE 18:1_20:1 | 772.585 | 770.5705 | 18.8 | | | |
| PE P-38:2 | PE P-18:1_20:1 | 756.5901 | 754.5756 | 19.39 | | 0.59 | |
| PE O-38:3 | PE O-18:2_20:1 | | | | | | |
| PE P-38:2 | PE P-20:1_18:1 | 756.5901 | 754.5756 | 19.39 | | 0.59 | |
| PE O-38:3 | PE O-20:2_18:1 | | | | | | |
| PE 38:3 | PE 18:0_20:3 | 770.5694 | 768.5548 | 18.3 | | | |
| PE P-38:3 | PE P-18:0_20:3 | 754.5745 | 752.5599 | 18.89 | | 0.59 | |
| PE O-38:4 | PE O-18:1_20:3 | | | | 1.89 | | |
| PE 38:4 | PE 16:0_22:4 | 768.5537 | 766.5392 | 17.35 | | | |
| PE O-38:4 | PE O-16:0_22:4 | 754.5745 | 752.5599 | 18.18 | 0.83 | | 0.23 |
| PE P-38:4 | PE P-16:0_22:4 | 752.5588 | 750.5443 | 17.95 | | 0.6 | |
| PE O-38:5 | PE O-16:1_22:4 | | | | | | |
| PE 38:4 | PE 18:0_20:4 | 768.5537 | 766.5392 | 17.8 | | | |
| PE O-38:4 | PE O-18:0_20:4 | 754.5745 | 752.5599 | 18.57 | 0.77 | | 0.19 |
| PE P-38:4 | PE P-18:0_20:4 | 752.5588 | 750.5443 | 18.38 | | 0.58 | |
| PE O-38:5 | PE O-18:1_20:4 | | | | 1.92 | | |
| PE 38:4 | PE 18:1_20:3 | 768.5537 | 766.5392 | 17 | | | |
| PE P-38:4 | PE P-18:1_20:3 | 752.5588 | 750.5443 | 17.63 | | 0.63 | |
| PE O-38:5 | PE O-18:2_20:3 | | | | 1.17 | | |
| PE 38:4 | PE 18:2_20:2 | 768.5537 | 766.5392 | 17.3 | | | |
| PE O-38:4 | PE O-16:1_22:3 | 754.5745 | 752.5599 | 18.57 | | | |
| PE 38:5 | PE 16:0_22:5 | 766.5381 | 764.5235 | 16.71 | | | |
| PE P-38:5 | PE P-16:0_22:5 | 750.5432 | 748.5286 | 17.29 | | 0.58 | |
| PE O-38:6 | PE O-16:1_22:5 | | | | 1.2 | | |
| PE 38:5 | PE 18:0_20:5 | 766.5381 | 764.5235 | 16.97 | | | |
| PE P-38:5 | PE P-18:0_20:5 | 750.5432 | 748.5286 | 17.56 | | 0.59 | |
| PE O-38:6 | PE O-18:1_20:5 | | | | 2.01 | | |
| PE 38:5 | PE 18:1_20:4 | 766.5381 | 764.5235 | 16.46 | | | |
| PE P-38:5 | PE P-18:1_20:4 | 750.5432 | 748.5286 | 17.08 | | 0.62 | |
| PE O-38:6 | PE O-18:2_20:4 | | | | 1.75 | | |
| PE 38:6 | PE 16:0_22:6 | 764.5224 | 762.5079 | 16.09 | | | |
| PE P-38:6 | PE P-16:0_22:6 | 748.5275 | 746.513 | 16.67 | | 0.58 | |
| PE O-38:7 | PE O-16:1_22:6 | | | | | | |
| PE 38:6 | PE 18:1_20:5 | 764.5224 | 762.5079 | 15.55 | | | |
| PE P-38:6 | PE P-18:1_20:5 | 748.5275 | 746.513 | 16.17 | | 0.62 | |
| PE O-38:7 | PE O-18:2_20:5 | | | | | | |
| PE 38:6 | PE 18:2_20:4 | 764.5224 | 762.5079 | 15.33 | | | |
| PE P-38:6 | PE P-18:2_20:4 | 748.5275 | 746.513 | 15.96 | | 0.63 | |
| PE O-38:7 | PE O-18:3_20:4 | | | | | | |
| PE 39:4 | PE 19:0_20:4 | 782.5694 | 780.5548 | 18.41 | | | |
| PE P-39:4 | PE P-17:0_22:4 | 766.5745 | 764.5599 | 18.6 | | | |
| PE P-39:5 | PE P-17:0_22:5 | 764.5588 | 762.5443 | 17.78 | | | |
| PE 40:1 | PE 18:0_22:1 | 802.632 | 800.6174 | 21.05 | | | |

| | | | | | | | |
|---|---|---|---|---|---|---|---|
| **PE 40:2** | **PE 18:1_22:1** | 800.6163 | 798.6018 | 19.88 | | | |
| **PE P-40:2** | **PE P-18:1_22:1** | **784.6214** | **782.6069** | **20.4** | | **0.52** | |
| **PE O-40:3** | **PE O-18:2_22:1** | | | | | | |
| **PE P-40:3** | **PE P-18:0_22:3** | 782.6058 | 780.5912 | 19.99 | | | |
| **PE P-40:3** | **PE P-20:0_20:3** | 782.6058 | 780.5912 | 19.99 | | | |
| **PE 40:4** | **PE 18:0_22:4** | 796.585 | 794.5705 | 18.64 | | | |
| **PE O-40:4** | **PE O-18:0_22:4** | 782.6058 | 780.5912 | 19.38 | **0.74** | | **0.17** |
| **PE P-40:4** | **PE P-18:0_22:4** | **780.5901** | **778.5756** | **19.21** | | **0.57** | |
| **PE O-40:5** | **PE O-18:1_22:4** | | | | **1.86** | | |
| **PE 40:4** | **PE 20:0_20:4** | 796.585 | 794.5705 | 19.01 | | | |
| **PE P-40:4** | **PE P-20:0_20:4** | **780.5901** | **778.5756** | **19.53** | | **0.52** | |
| **PE O-40:5** | **PE O-20:1_20:4** | | | | **1.85** | | |
| **PE 40:5** | **PE 18:0_22:5** | 794.5694 | 792.5548 | 17.85 | | | |
| **PE O-40:5** | **PE O-18:0_22:5** | 780.5901 | 778.5756 | 18.6 | **0.75** | | **0.2** |
| **PE P-40:5** | **PE P-18:0_22:5** | **778.5745** | **776.5599** | **18.4** | | **0.55** | |
| **PE O-40:6** | **PE O-18:1_22:5** | | | | **1.87** | | |
| **PE 40:5** | **PE 18:1_22:4** | 794.5694 | 792.5548 | 17.35 | | | |
| **PE P-40:5** | **PE P-18:1_22:4** | **778.5745** | **776.5599** | **17.95** | | **0.6** | |
| **PE O-40:6** | **PE O-18:2_22:4** | | | | **1.42** | | |
| **PE 40:5** | **PE 20:1_20:4** | 794.5694 | 792.5548 | 17.68 | | | |
| **PE P-40:5** | **PE P-20:1_20:4** | **778.5745** | **776.5599** | **18.25** | | **0.57** | |
| **PE O-40:6** | **PE O-20:2_20:4** | | | | **1.72** | | |
| **PE 40:6** | **PE 18:0_22:6** | 792.5537 | 790.5392 | 17.47 | | | |
| **PE P-40:6** | **PE P-18:0_22:6** | **776.558** | **774.5443** | **18.02** | | **0.55** | |
| **PE O-40:7** | **PE O-18:1_22:6** | | | | **1.93** | | |
| **PE 40:6** | **PE 18:1_22:5** | 792.5537 | 790.5392 | 16.53 | | | |
| **PE P-40:6** | **PE P-18:1_22:5** | **776.558** | **774.5443** | **17.11** | | **0.58** | |
| **PE O-40:7** | **PE O-18:2_22:5** | | | | | | |
| **PE 40:7** | **PE 18:1_22:6** | 790.5381 | 788.5235 | 16.09 | | | |
| **PE P-40:7** | **PE P-18:1_22:6** | **774.5432** | **772.5286** | **16.71** | | **0.62** | |
| **PE O-40:8** | **PE O-18:2_22:6** | | | | | | |

Finally, structure – RT relationships for all identified PE species were visualized by plotting Kendrick mass defect by hydrogen (KMD(H)) vs RT plot to control identification accuracy (Figure 4.3 and 4.4). All species falling out of the diagonal (different carbon number but the same DBE) and horizontal (same carbon number but different DBE) trend lines were excluded. To simplify the visualization, A-PE were plotted separately form O-/P-linked PE lipids.

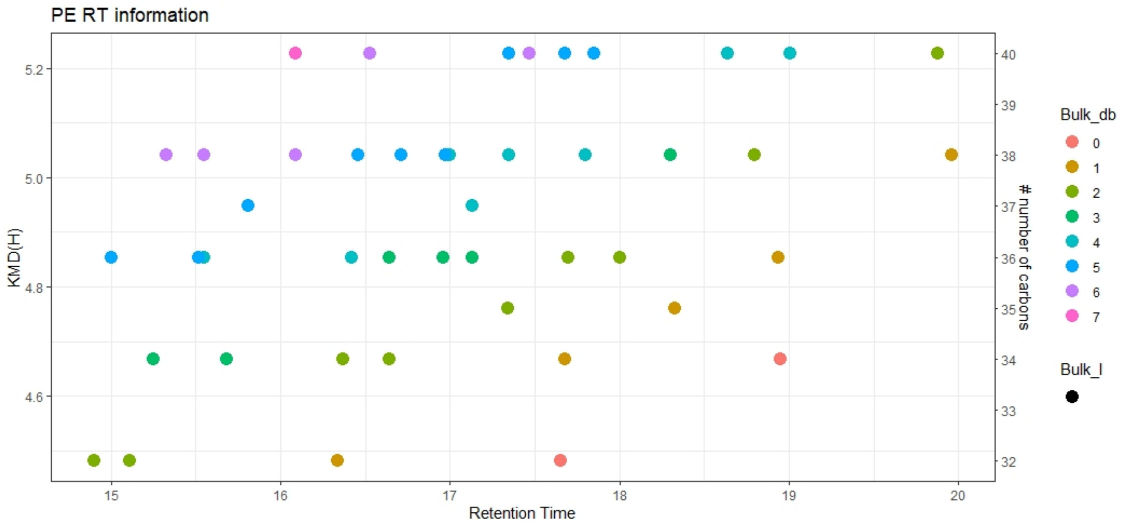


**Figure 4.3**. KMD(H), A-PE carbohydrate chain carbon number vs RT plot for all identified A-PE species. Symbols color represents the number of double bounds in carbohydrate chains.

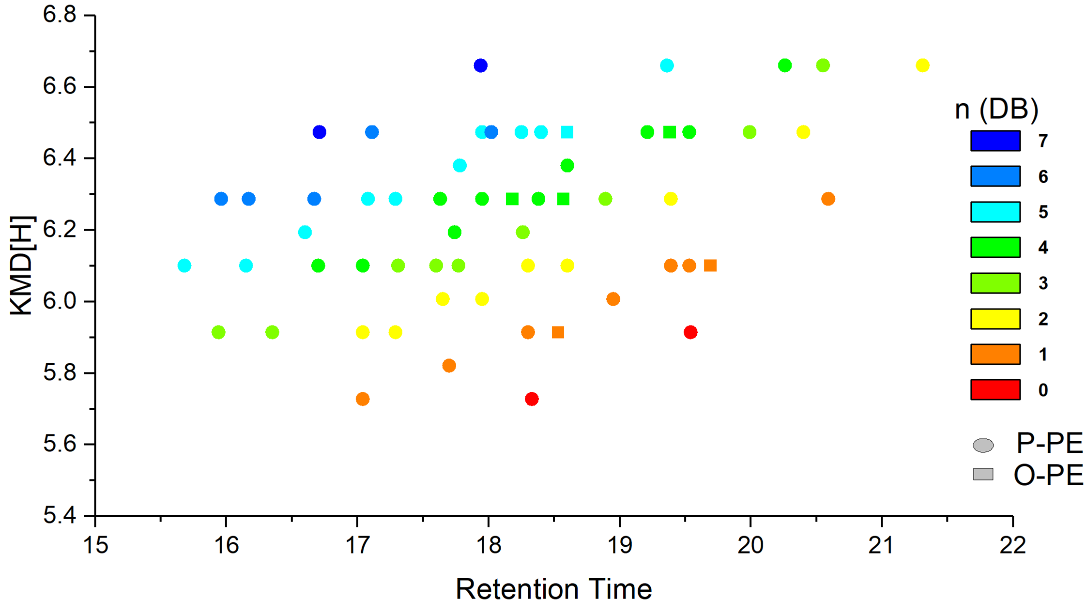


**Figure 4.4**. KMD(H), PE carbohydrate chain carbon number vs RT plot for all identified O-/P-PE species. Symbols color represents the number of double bounds in carbohydrate chains. Triangles – alkyl LPE (O-PE), and square – alkenyl ether PE (P-LPE).

#### 5. Manual confirmation of identities for PI lipid molecular species.

Software assisted identification of PI molecular species required further manual confirmation to assure accurate identification based on specific fragment ions and retention time mapping.

##### 5.1. Monitored adducts

PI were monitored as protonated or deprotonated ions in positive and negative ion modes, respectively.

##### 5.2. Fragmentation patterns

General fragmentation pattern and representative MS/MS spectra for PI ionized in positive and negative modes are illustrated below:

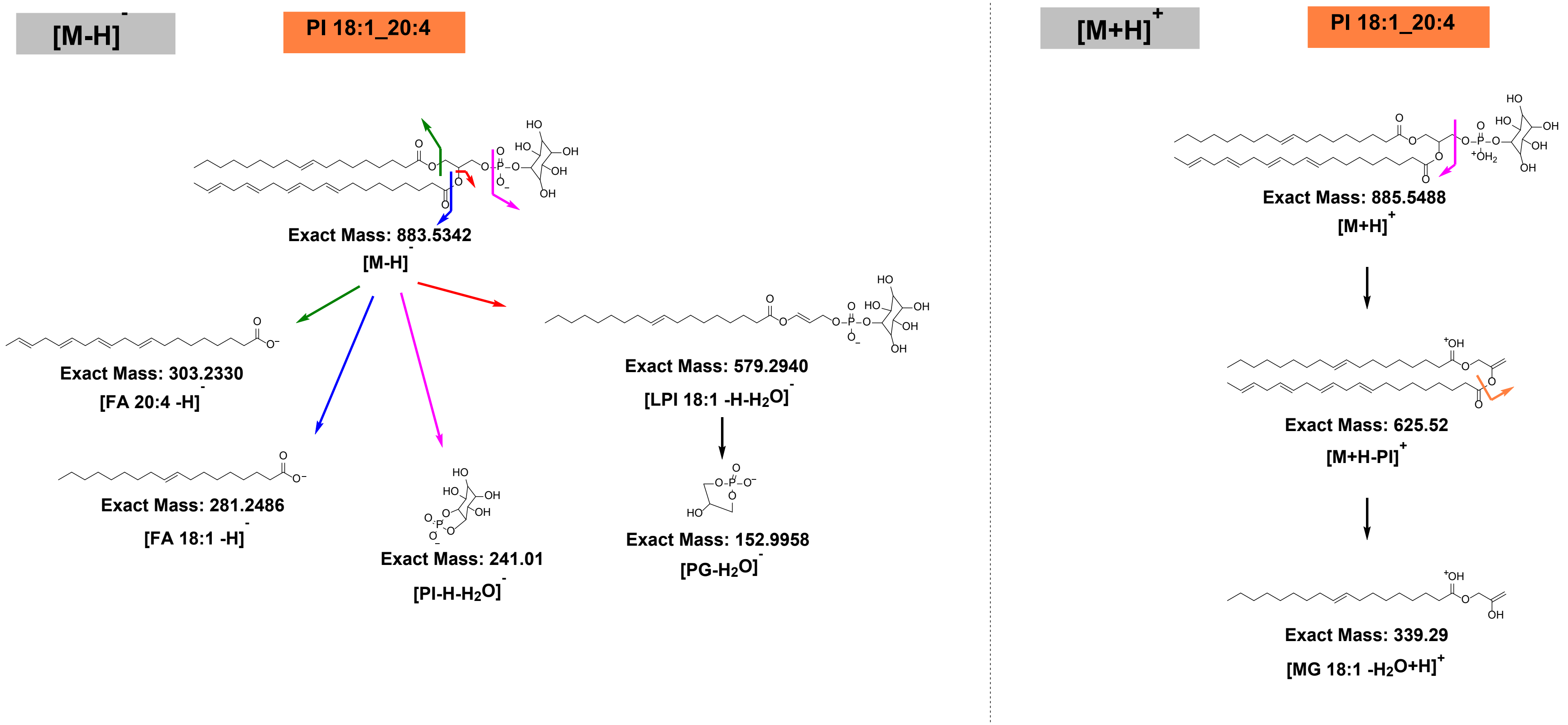


**Scheme 5.1.** General HCD fragmentation pattern of deprotonated [M-H]- and protonated [M+H]+ adducts of PI.

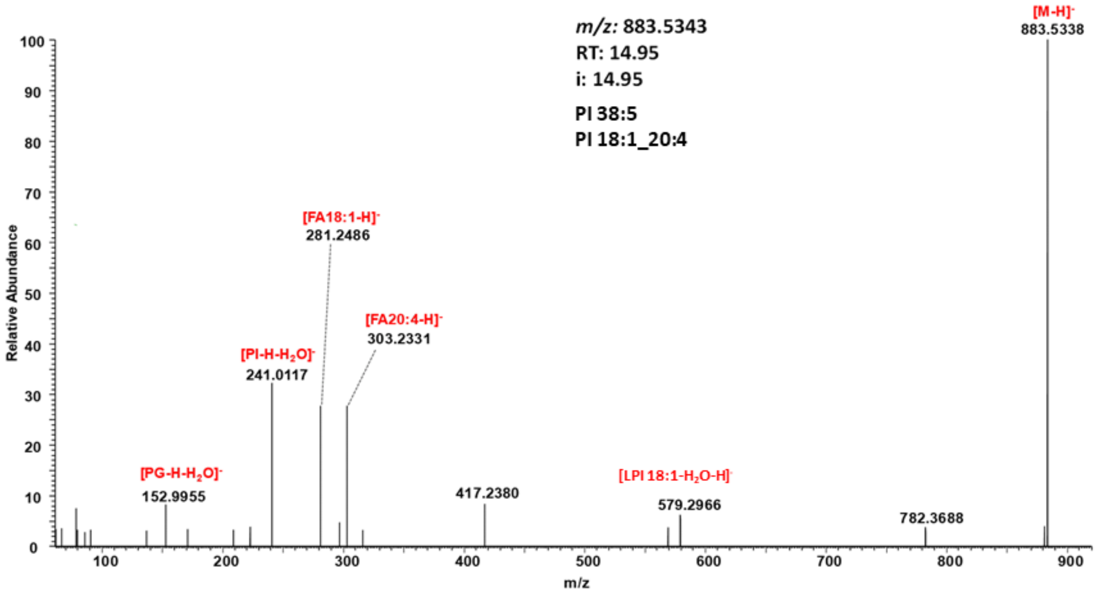


**Figure 5.1.** Representative HCD spectrum of the deprotonated adduct $[M-H]^-$ of PI 18:1_20:4.

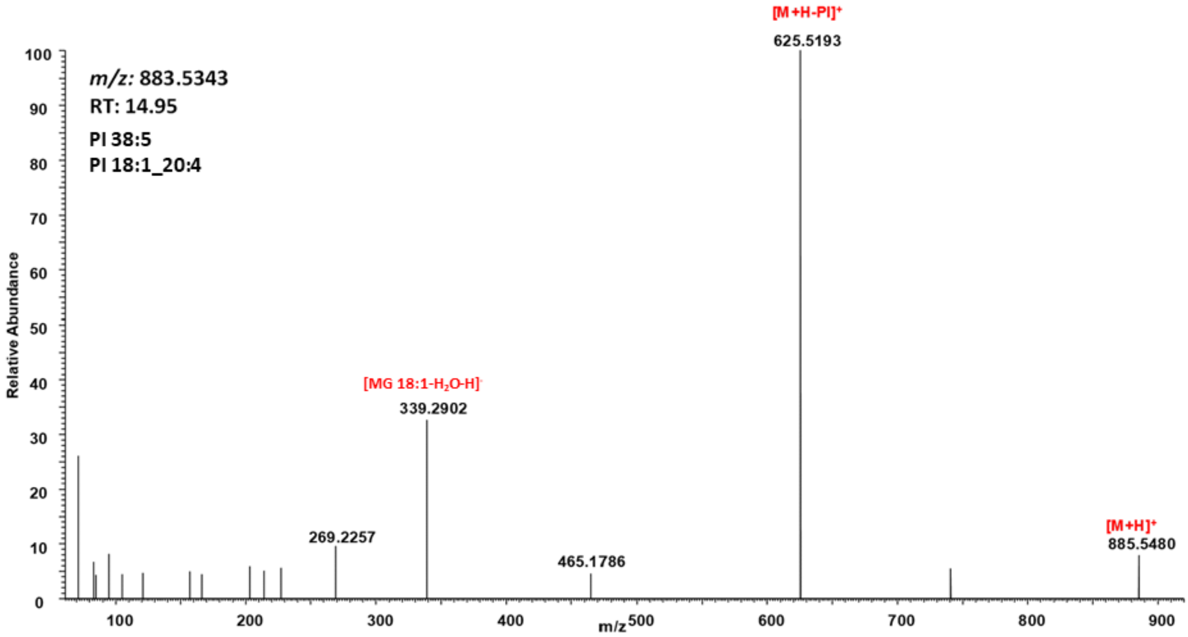


**Figure 5.2.** Representative HCD spectrum of the protonated adduct $[M+H]^+$ of PI 18:1_20:4.

**Table 5.1.** Summary of PI specific neutral loss and fragment ions obtained by positive and negative ion modes HCD.

| NL/Fragment ion | PI |
|---|---|
| **Negative ion mode HCD** | |
| FI: fatty acyl 1 anion | + |
| FI: fatty acyl 2 anion | + |
| FI: LPI fatty acyl 1 anion | + |
| FI: 241 $[PI-H-H2O]^-$ | + |
| FI: 152 $[PG-H_2O]^-$ | + |

| Positive ion mode HCD | |
|---|---|
| NL: 260 (phosphoinositol) | + |
| FI: fatty acyl monoglycerol $[MG-H_2O+H]^+$ | + |

##### 5.3. RT mapping

To establish retention time rules for various PIs, molecular species were used (Table 5.2).

**Table 5.2.** Calculated RT differences between PI lipids used to identify true and false positive IDs.

| Bulk | Discrete | RT | Δ PC-PI | RT | Discrete | Bulk |
|---|---|---|---|---|---|---|
| **PI 32:0** | **PI 16:0_16:0** | **15.98** | 1.44 | 17.42 | PC 16:0_16:0 | PC 32:0 |
| **PI 34:1** | **PI 16:0_18:1** | **16.06** | 1.4 | 17.46 | PC 16:0_18:1 | PC 34:1 |
| **PI 34:2** | **PI 16:1_18:1** | **14.71** | 1.43 | 16.14 | PC 16:1_18:1 | PC 34:2 |
| **PI 34:2** | **PI 16:0_18:2** | **14.99** | 1.38 | 16.37 | PC 16:0_18:2 | PC 34:2 |
| **PI 36:1** | **PI 18:0_18:1** | **17.38** | 1.41 | 18.79 | PC 18:0_18:1 | PC 36:1 |
| **PI 36:2** | **PI 18:1_18:1** | **16.09** | 1.38 | 17.47 | PC 18:1_18:1 | PC 36:2 |
| **PI 36:2** | **PI 18:0_18:2** | **16.42** | 1.39 | 17.81 | PC 18:0_18:2 | PC 36:2 |
| **PI 36:3** | **PI 18:1_18:2** | **15.07** | 1.35 | 16.42 | PC 18:1_18:2 | PC 36:3 |
| **PI 36:3** | **PI 16:0_20:3** | **15.38** | 1.37 | 16.75 | PC 16:0_20:3 | PC 36:3 |
| **PI 36:4** | **PI 16:0_20:4** | **14.83** | 1.34 | 16.17 | PC 16:0_20:4 | PC 36:4 |
| **PI 37:4** | **PI 17:0_20:4** | **15.55** | 1.37 | 16.92 | PC 17:0_20:4 | PC 37:4 |
| **PI 38:3** | **PI 18:0_20:3** | **16.75** | 1.4 | 18.15 | PC 18:0_20:3 | PC 38:3 |
| **PI 38:4** | **PI 18:1_20:3** | **15.88** | 0.91 | 16.79 | PC 18:1_20:3 | PC 38:4 |
| **PI 38:4** | **PI 18:0_20:4** | **16.23** | 1.37 | 17.6 | PC 18:0_20:4 | PC 38:4 |
| **PI 38:4** | **PI 16:0_22:4** | **15.87** | 1.29 | 17.16 | PC 16:0_22:4 | PC 38:4 |
| **PI 38:5** | **PI 18:1_20:4** | **14.86** | 1.36 | 16.22 | PC 18:1_20:4 | PC 38:5 |
| **PI 38:5** | **PI 18:0_20:5** | **15.39** | 1.32 | 16.71 | PC 18:0_20:5 | PC 38:5 |
| **PI 40:4** | **PI 18:0_22:4** | **17.13** | 1.36 | 18.49 | PC 18:0_22:4 | PC 40:4 |
| **PI 40:6** | **PI 18:0_22:6** | **15.94** | 1.33 | 17.27 | PC 18:0_22:6 | PC 40:6 |

Finally, structure – RT relationships for all identified PI species were visualized by plotting Kendrick mass defect by hydrogen (KMD(H)) vs RT plot to control identification accuracy (Figure 5.3). All species falling out of the diagonal (different carbon number but the same DBE) and horizontal (same carbon number but different DBE) trend lines were excluded.

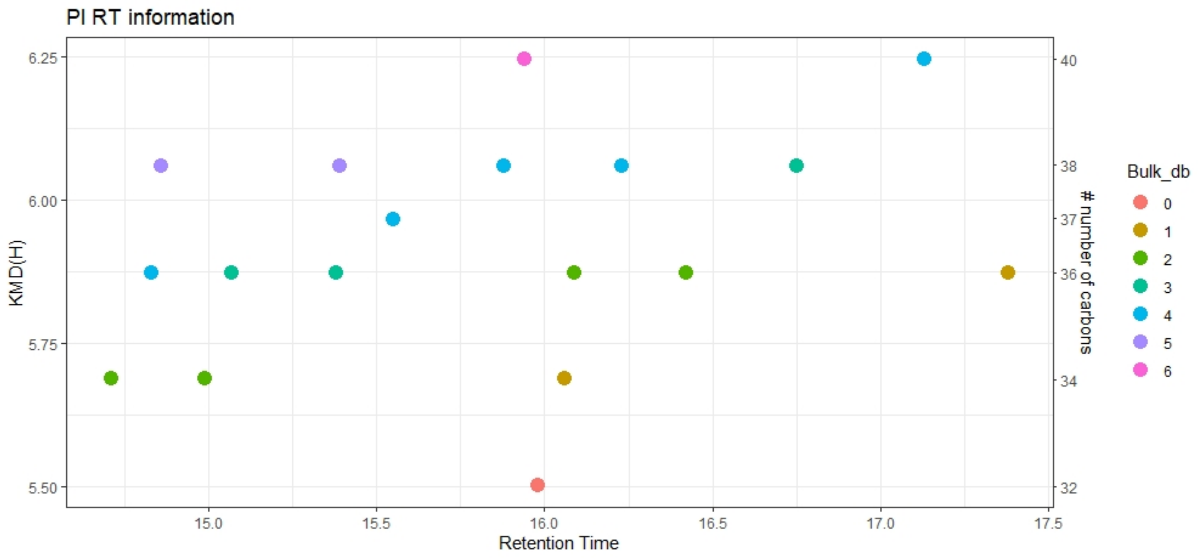


**Figure 5.3**. KMD(H), PI carbohydrate chain carbon number vs RT plot for all identified PI species. Symbols color represents the number of double bounds in carbohydrate chains.

#### 6. Manual confirmation of identities for PS lipid molecular species.

Software assisted identification of PS molecular species required further manual confirmation to assure accurate identification based on specific fragment ions and retention time mapping.

##### 6.1. Monitored adducts

PS were monitored as deprotonated ions in negative ion mode.

##### 6.2. Fragmentation patterns

General fragmentation pattern and representative MS/MS spectrum for PS ionized in negative mode are illustrated below:

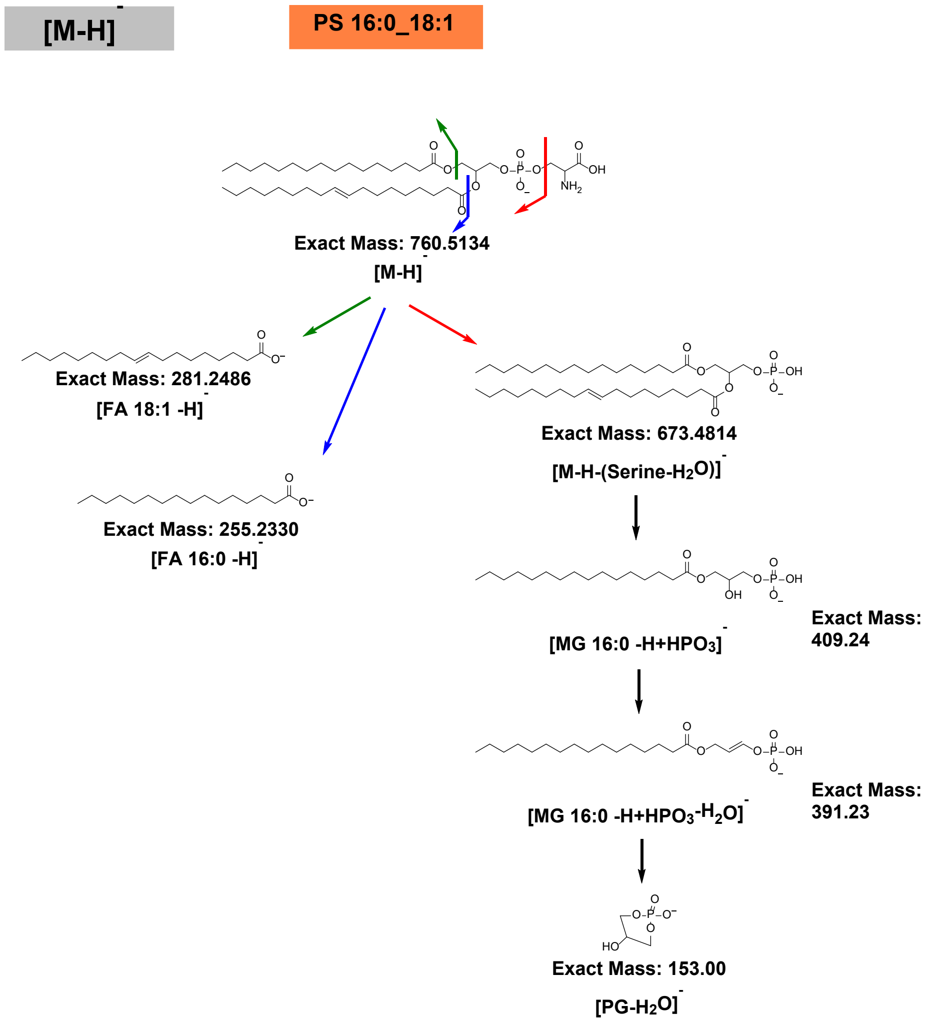


**Scheme 6.1.** General HCD fragmentation pattern of deprotonated $[M-H]^-$ adducts of PS.

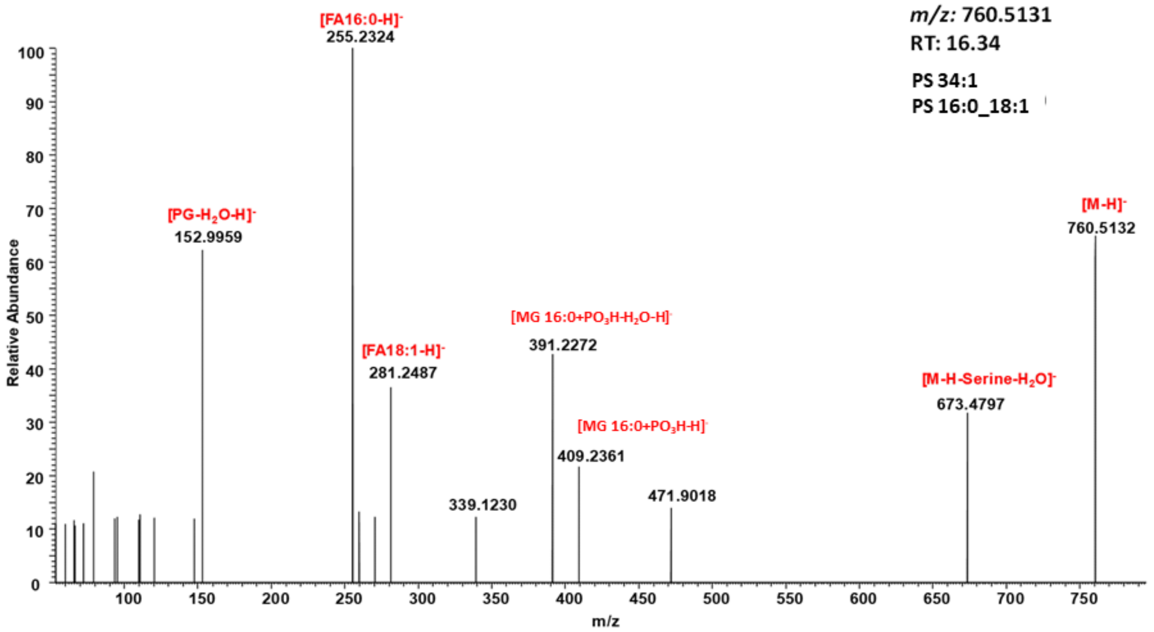


**Figure 6.1.** Representative HCD spectrum of the deprotonated adduct $[M-H]^-$ of PS 16:0_18:1.

**Table 6.1.** Summary of PS specific neutral loss and fragment ions obtained by positive and negative ion modes HCD.

| NL/Fragment ion | PS |
|---|---|
| **Positive ion mode HCD** | |
| FI: fatty acyl anion 1 | + |
| FI: fatty acyl anion 2 | + |
| NL: 87 (serine) | + |
| FI: monoglycerol + phosphate | + |
| FI: monoglycerol – $H_2O$ + phosphate | + |
| FI: phosphatidylglycerol – $H_2O$ | + |

##### 6.3. RT mapping

To establish retention time rules for various PSs, molecular species were used (Table 6.2).

**Table 6.2.** Calculated RT differences between PS lipids used to identify true and false positive IDs.

| Bulk | Discrete | RT | Δ PC-PS | RT | Discrete | Bulk |
|---|---|---|---|---|---|---|
| **PS 34:1** | **PS 16:0_18:1** | **16.34** | 1.12 | 17.46 | PC 16:0_18:1 | PC 34:1 |
| **PS 34:2** | **PS 16:0_18:2** | **15.28** | 1.09 | 16.37 | PC 16:0_18:2 | PC 34:2 |
| **PS 36:1** | **PS 18:0_18:1** | **17.7** | 1.09 | 18.79 | PC 18:0_18:1 | PC 36:1 |
| **PS 36:2** | **PS 18:0_18:2** | **16.68** | 1.13 | 17.81 | PC 18:0_18:2 | PC 36:2 |
| **PS 36:2** | **PS 18:1_18:1** | **16.34** | 1.13 | 17.47 | PC 18:1_18:1 | PC 36:2 |
| **PS 36:3** | **PS 18:1_18:2** | **15.67** | 0.75 | 16.42 | PC 18:1_18:2 | PC 36:3 |
| **PS 36:4** | **PS 18:2_18:2** | **16.62** | **-1.29** | 15.33 | PC 18:2_18:2 | PC 36:4 |
| **PS 38:3** | **PS 18:0_20:3** | **17.03** | 1.12 | 18.15 | PC 18:0_20:3 | PC 38:3 |
| **PS 38:4** | **PS 18:0_20:4** | **16.53** | 1.07 | 17.6 | PC 18:0_20:4 | PC 38:4 |

| | | | | | | |
|---|---|---|---|---|---|---|
| **PS 38:5** | **PS 18:1_20:4** | **16.67** | **-0.45** | 16.22 | PC 18:1_20:4 | PC 38:5 |
| **PS 40:6** | **PS 18:0_22:6** | **16.19** | 1.08 | 17.27 | PC 18:0_22:6 | PC 40:6 |

Finally, structure – RT relationships for all identified PS species were visualized by plotting Kendrick mass defect by hydrogen (KMD(H)) vs RT plot to control identification accuracy (Figure 6.2). All species falling out of the diagonal (different carbon number but the same DBE) and horizontal (same carbon number but different DBE) trend lines were excluded.

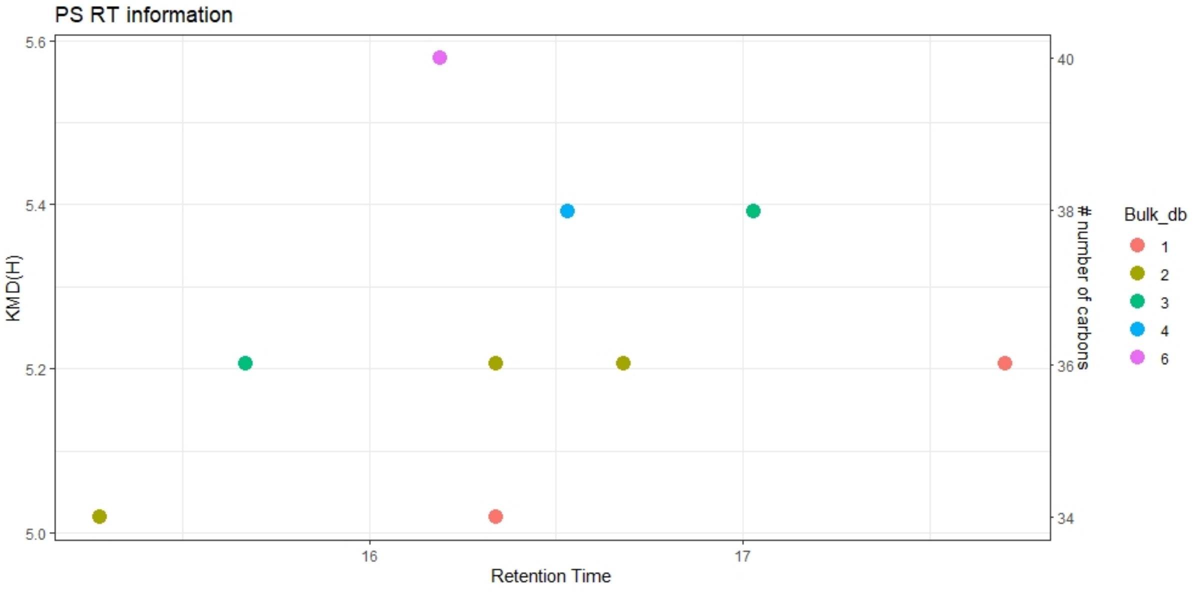


**Figure 6.2**. KMD(H), PS carbohydrate chain carbon number vs RT plot for all identified PS species. Symbols color represents the number of double bounds in carbohydrate chains

#### 7. Manual confirmation of identities for PG lipid molecular species.

Software assisted identification of PG molecular species required further manual confirmation to assure accurate identification based on specific fragment ions and retention time mapping.

##### 7.1. Monitored adducts

PG were monitored as protonated or deprotonated ions in positive and negative ion modes, respectively.

##### 7.2. Fragmentation patterns

General fragmentation pattern and representative MS/MS spectra for PG ionized in positive and negative modes are illustrated below:

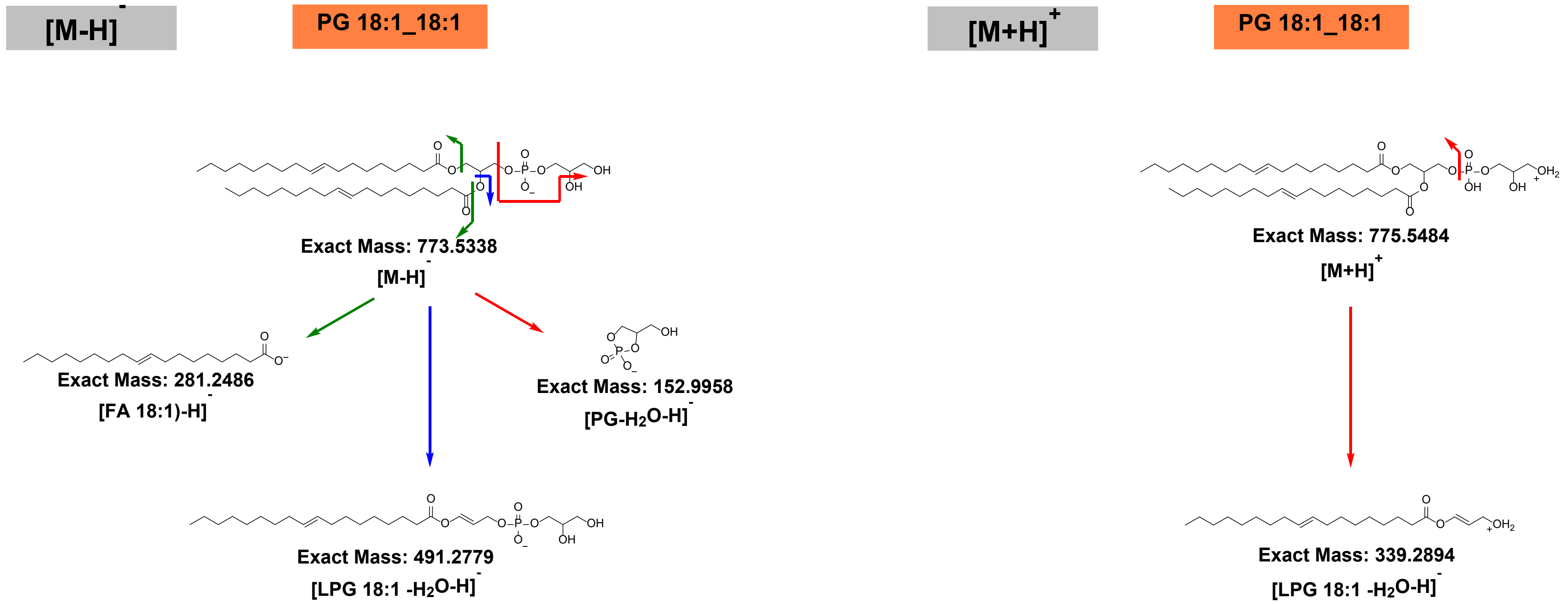


**Scheme 7.1.** General HCD fragmentation pattern of deprotonated $[M-H]^-$ and protonated $[M+H]^+$ adducts of PG.

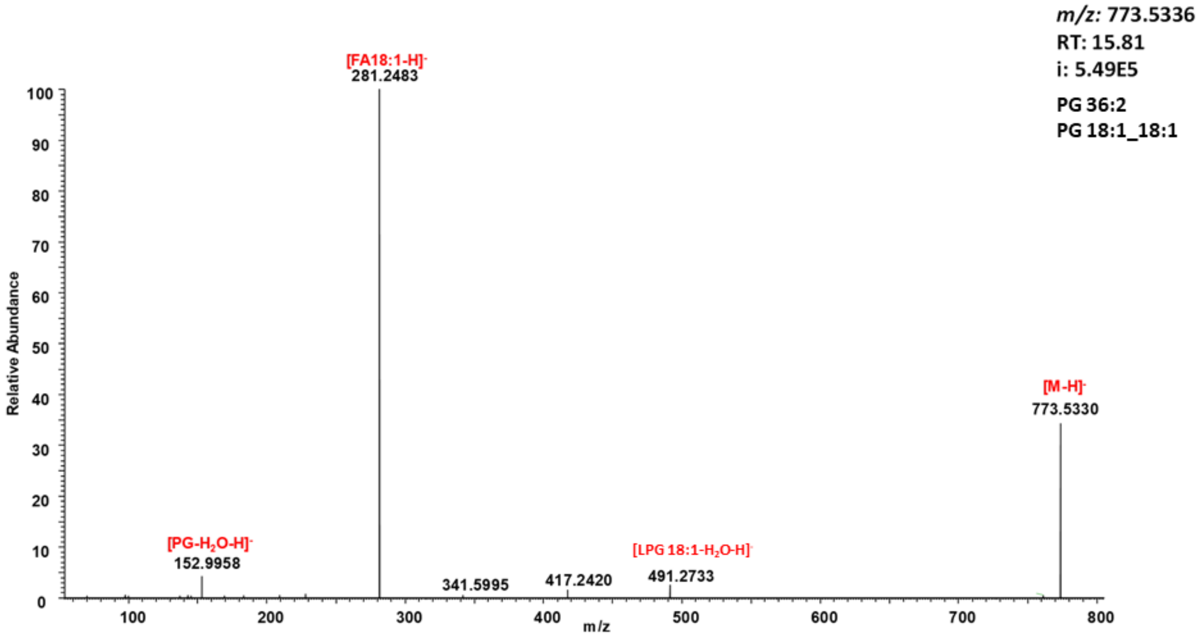


**Figure 7.1.** Representative HCD spectrum of the deprotonated adduct $[M-H]^-$ of PG 18:1_18:1.

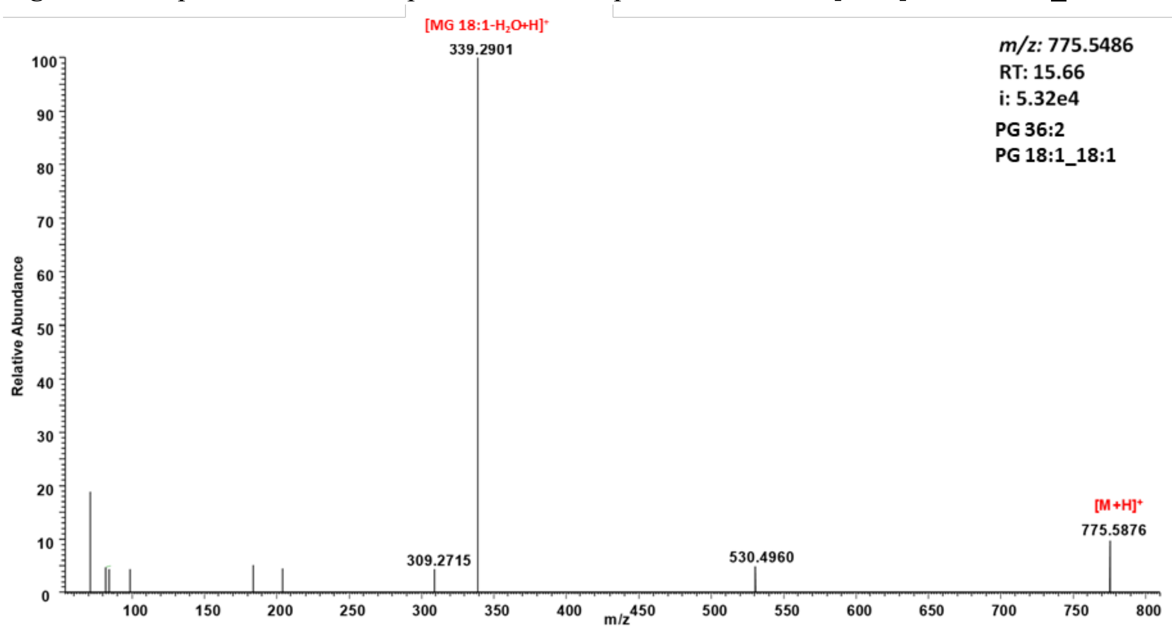


**Figure 7.2.** Representative HCD spectrum of the protonated adduct $[M+H]^+$ of PG 18:1_18:1.

**Table 7.1.** Summary of PG specific neutral loss and fragment ions obtained by positive and negative ion modes HCD.

| NL/Fragment ion | PG |
|---|---|
| **Negative ion mode HCD** | |
| FI: fatty acyl 1 anion | + |
| FI: fatty acyl 2 anion | + |
| FI: LPG fatty acyl 1 anion | + |
| FI: 152 $[PG-H_2O]^-$ | + |
| **Positive ion mode HCD** | |
| NL: 260 (phosphoinositol) | + |
| FI: lysophosphatidylglycerol – $H_2O$ $[LPG-H_2O+H]^+$ | + |

##### 7.3. RT mapping

To establish retention time rules for various PGs, molecular species were used (Table 7.2).

**Table 7.2.** Calculated RT differences between PG lipids used to identify true and false positive IDs.

| Bulk | Discrete | RT | Δ PC-PE | RT | Discrete | Bulk |
|---|---|---|---|---|---|---|
| **PG 34:2** | **PG 16:1_18:1** | **14.46** | **1.68** | 16.14 | **PC 16:1_18:1** | **PC 34:2** |
| **PG 34:1** | **PG 16:0_18:1** | **16.09** | **1.37** | 17.46 | **PC 16:0_18:1** | **PC 34:1** |
| **PG 36:4** | **PG 16:0_20:4** | **15.11** | **1.06** | 16.17 | **PC 16:0_20:4** | **PC 36:4** |
| **PG 36:4** | **PG 18:2_18:2** | **14.3** | **1.03** | 15.33 | **PC 18:2_18:2** | **PC 36:4** |
| **PG 36:2** | **PG 18:1_18:1** | **15.83** | **1.64** | 17.47 | **PC 18:1_18:1** | **PC 36:2** |
| **PG 36:1** | **PG 18:0_18:1** | **17.61** | **1.18** | 18.79 | **PC 18:0_18:1** | **PC 36:1** |
| **PG 38:5** | **PG 18:1_20:4** | **14.56** | **1.66** | 16.22 | **PC 18:1_20:4** | **PC 38:5** |
| **PG 38:4** | **PG 18:0_20:4** | **15.15** | **2.45** | 17.6 | **PC 18:0_20:4** | **PC 38:4** |

Finally, structure – RT relationships for all identified PG species were visualized by plotting Kendrick mass defect by hydrogen (KMD(H)) vs RT plot to control identification accuracy (Figure 7.3). All species falling out of the diagonal (different carbon number but the same DBE) and horizontal (same carbon number but different DBE) trend lines were excluded.

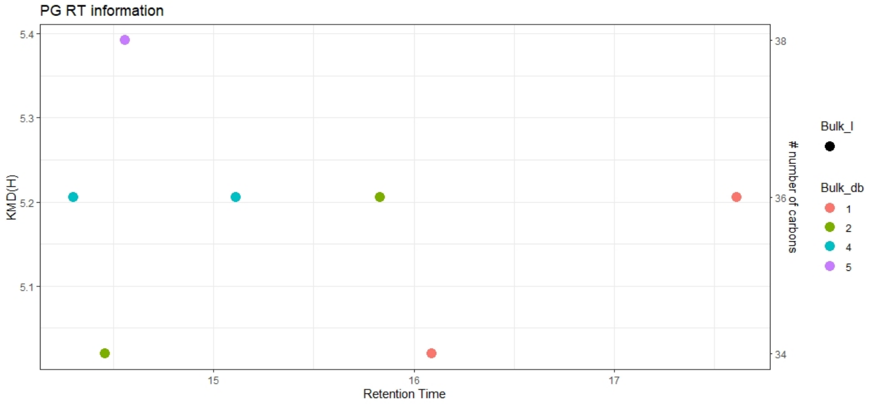


**Figure 7.3**. KMD(H), PI carbohydrate chain carbon number vs RT plot for all identified PG species. Symbols color represents the number of double bounds in carbohydrate chains.

#### Sphingolipids

##### 8. Manual confirmation of identities for SM lipid molecular species.

Software assisted identification of SM molecular species required further manual confirmation to assure accurate identification based on specific fragment ions and retention time mapping.

##### 8.1. Monitored adducts

SM were monitored as protonated ions in positive ion modes.

##### 8.2. Fragmentation patterns

General fragmentation pattern and representative MS/MS spectra for SM ionized in positive mode are illustrated below:

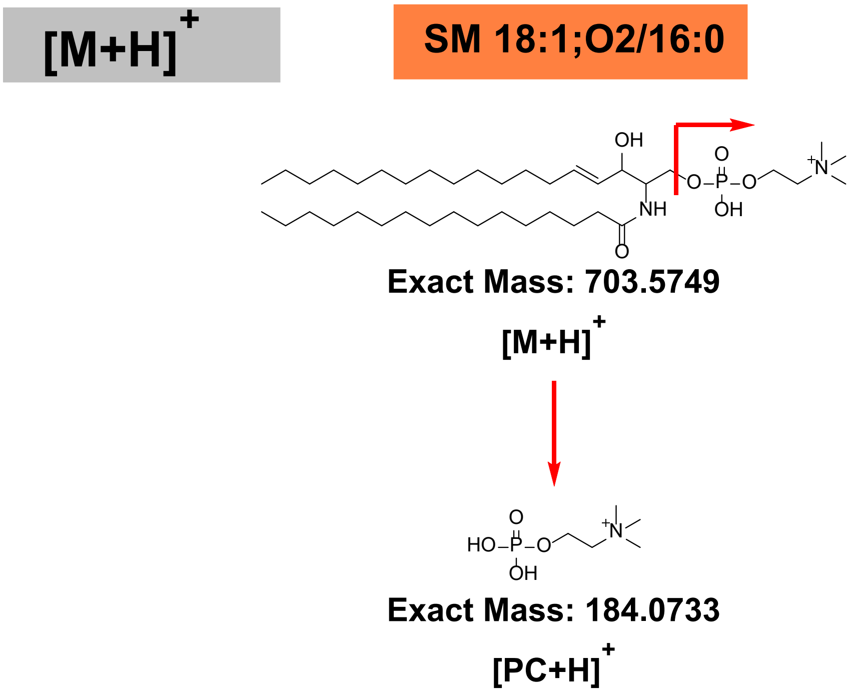


**Scheme 8.1.** General HCD fragmentation pattern of protonated [M+ H]$^{+}$ adducts of SM.

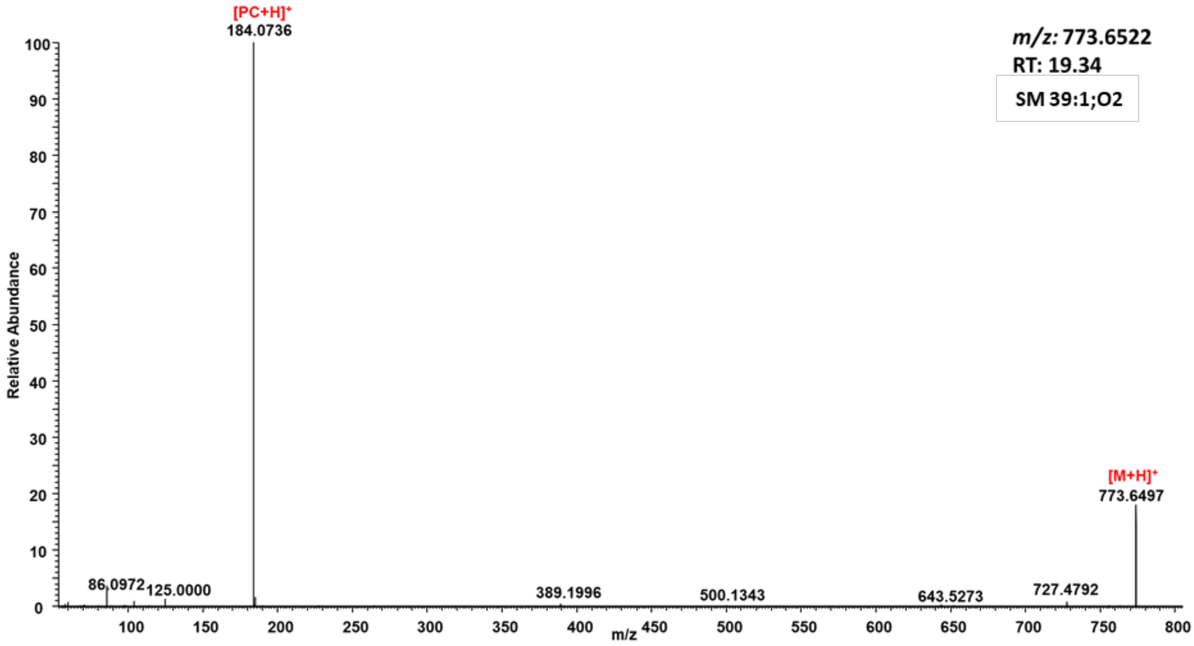


**Figure 8.1.** Representative HCD spectrum of the protonated adduct [M+H]$^{+}$ of SM 39:1;2O.

**Table 8.1.** Summary of SM specific neutral loss and fragment ions obtained by positive ion mode HCD.

| Positive ion mode HCD | |
|---|---|
| FI: 184 (phosphatidylcholine headgroup) | + |

##### 8.3. RT mapping

Finally, structure – RT relationships for all identified SM species were visualized by plotting Kendrick mass defect by hydrogen (KMD(H)) vs RT plot to control identification accuracy (Figure 8.2). All species falling out of the diagonal (different carbon number but the same DBE) and horizontal (same carbon number but different DBE) trend lines were excluded.

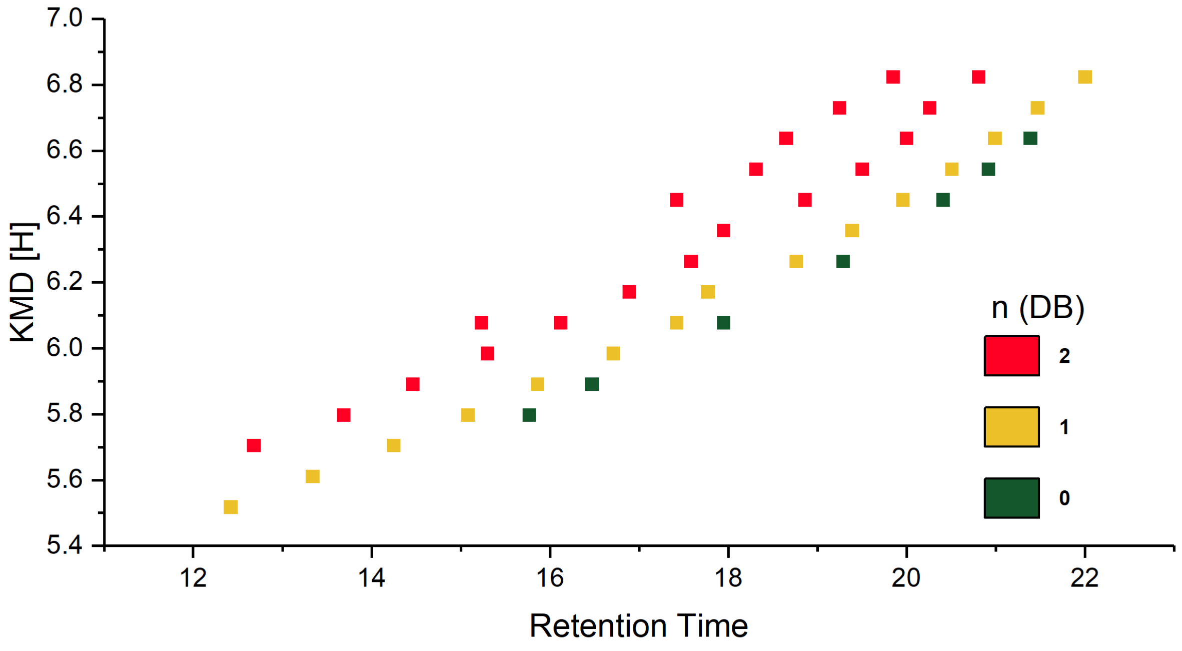


**Figure 8.2**. KMD(H), SM carbohydrate chain carbon number vs RT plot for all identified SM species. Symbols color represents the number of double bounds in carbohydrate chains.

#### 9. Manual confirmation of identities for ceramide lipid molecular species:

Software assisted identification of Cer molecular species required further manual confirmation to assure accurate identification of Cer subclasses (deoxyCer vs dihydroCer vs Cer vs phytoCer) based on specific fragment ions and retention time mapping. Cer subclasses contained sphingoid bases with varying numbers of hydroxyl groups (1 OH = deoxyCer, 2 OH = dihydroCer/Cer, 3 OH = phytoCer) and numbers of double bonds (0 DB = dihydroCer/phytoCer/deoxyDihydroCer, 1 DB = deoxyCer/Cer, 2 DB = deoxyCer/Cer)

##### 9.1. Monitored adducts

Cer were monitored as protonated adducts in positive mode.

##### 9.2. Fragmentation patterns

General fragmentation pattern and representative MS/MS spectra for Cer ionized in positive mode are illustrated below:

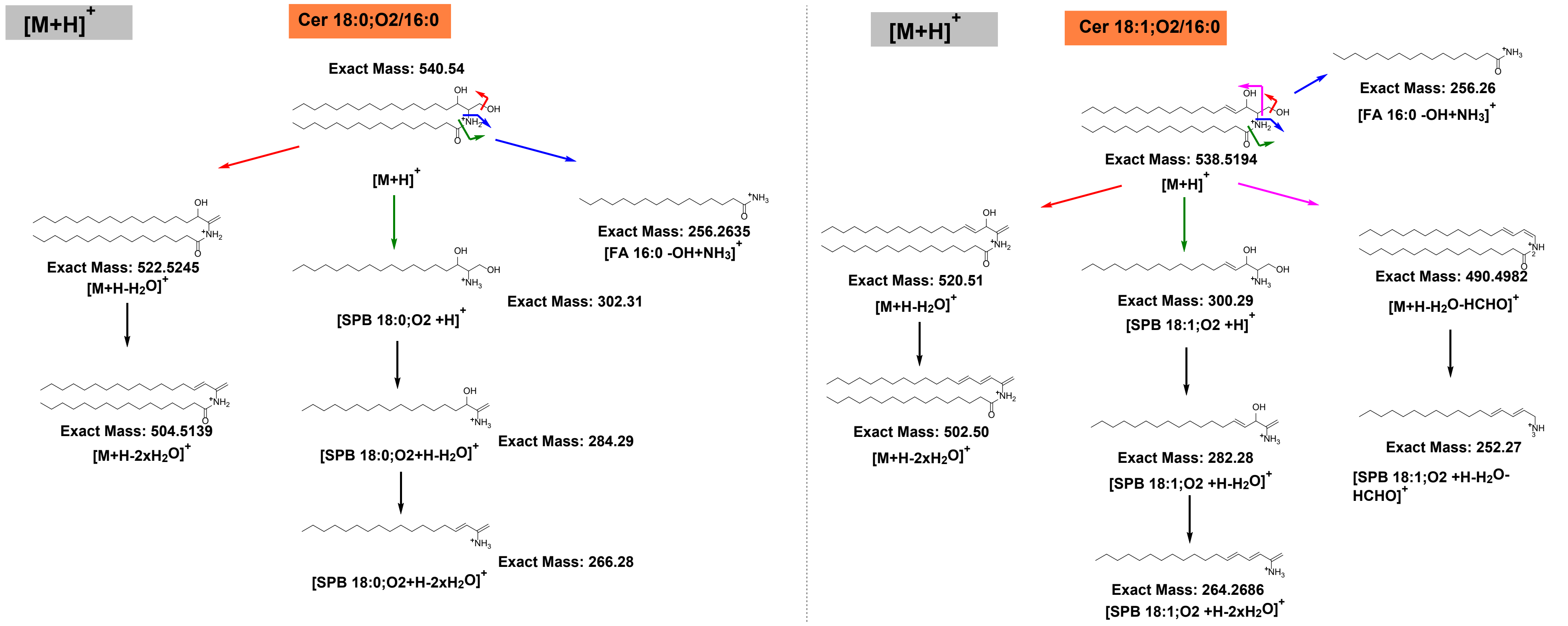


**Scheme 9.1.** General HCD fragmentation pattern of protonated dihydroCer (Cer 18:0;O2/16:0) and Cer (Cer 18:1;O2/16:0) as $[M+H]^+$.

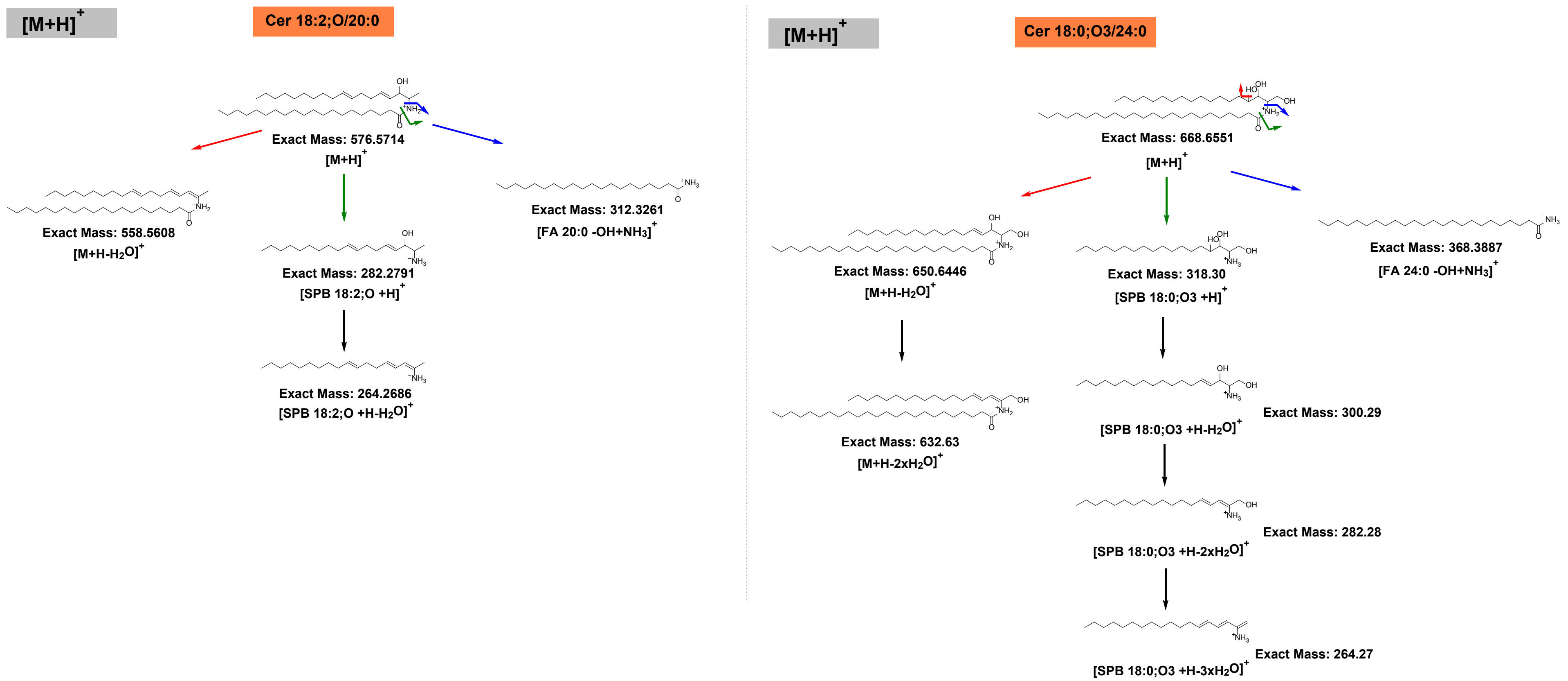


**Scheme 9.2.** General HCD fragmentation pattern of deoxyCer (Cer 18:2;O/20:0) and phytoCer (Cer 18:0;O3/24:0) protonated adducts $[M+H]^+$.

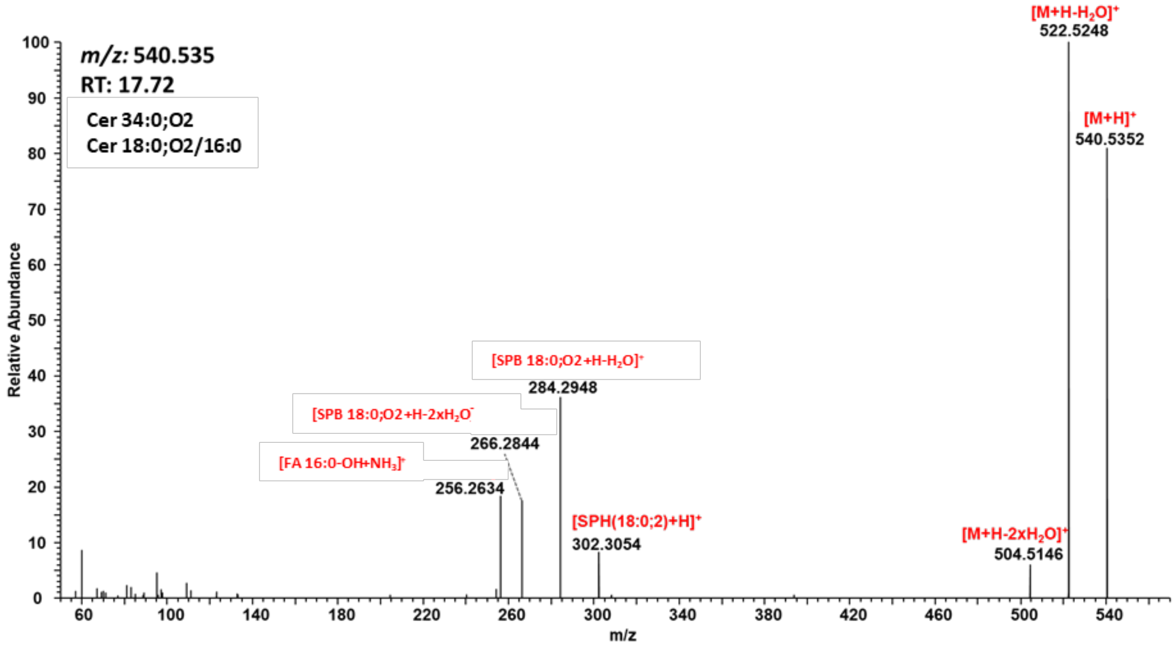


**Figure 9.1.** Representative HCD spectra of dihydroCer (Cer 18:0;O2/16:0) protonated adducts $[M+H]^+$.

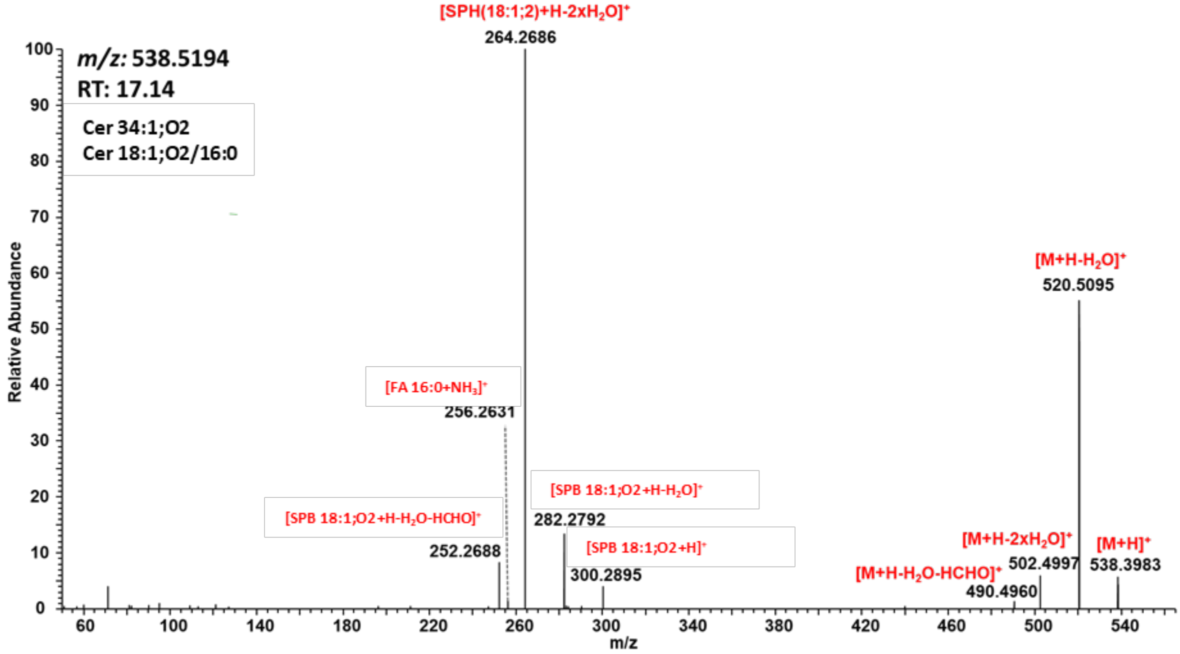


**Figure 9.2.** Representative HCD spectra of Cer (Cer 18:1;O2/16:0) protonated adduct $[M+H]^+$.

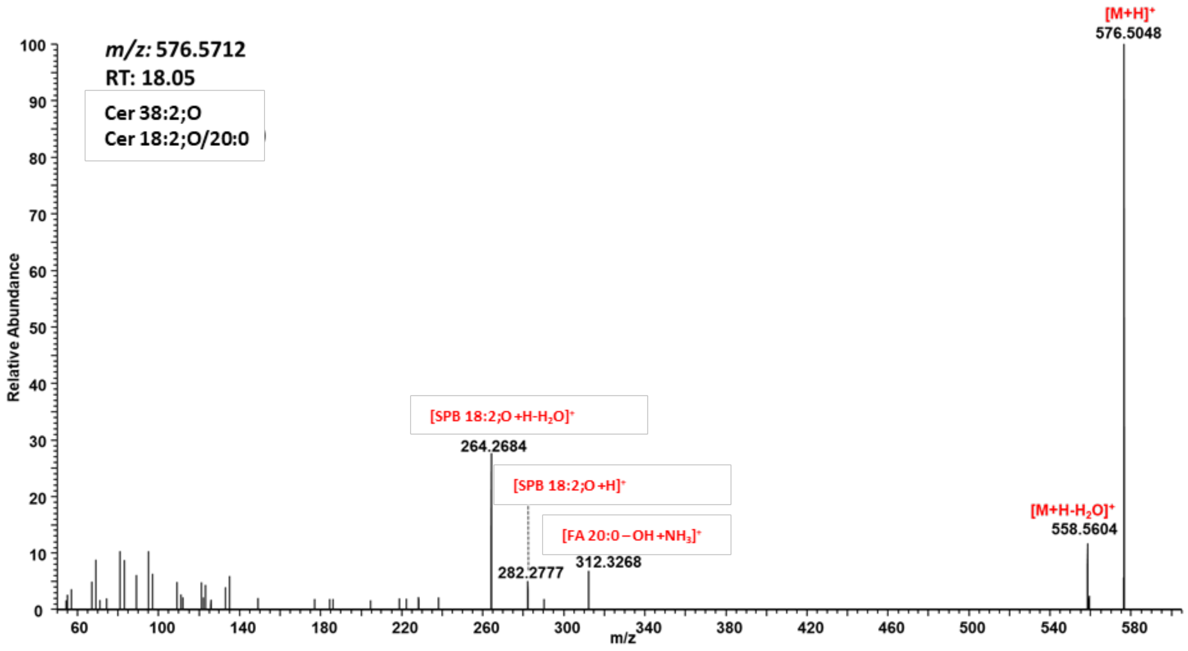


**Figure 9.3.** Representative HCD spectra of deoxyCer (Cer 18:2;O/20:0) protonated adduct $[M+H]^+$.

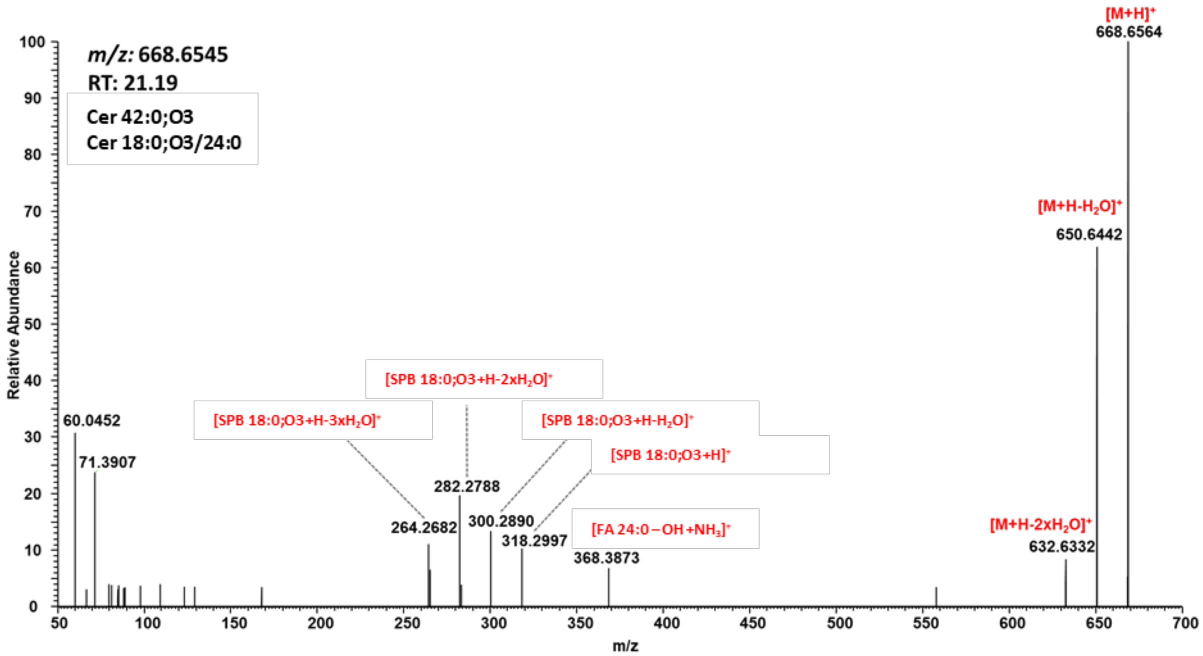


**Figure 9.4.** Representative HCD spectra of protonated adducts of phytoCer(Cer 18:0;O3/24:0) protonated adduct $[M+H]^+$.

**Table 9.1.** Summary of Cer subclass specific neutral loss (NL) and fragment ions (FI) obtained by positive HCD.

| NL/FI | DihydroCer | Cer | DeoxyCer | PhytoCer |
|---|---|---|---|---|
| NL: 18 ($H_2O$) | + | + | + | + |
| NL: 36 (2x $H_2O$) | + | + | | + |
| FI: sphinganine (18:0;O2) | + | | | |
| FI: sphinganine (18:0;O2) – $H_2O$ | + | | | |
| FI: sphinganine (18:0;O2) – $2xH_2O$ | + | | | |

| | | | | |
|---|---|---|---|---|
| FI: sphingosine (18:1,O2) | | + | | |
| FI: sphingosine (18:1,O2) – $H_2O$ | | + | | |
| FI: sphingosine (18:1,O2) – $2xH_2O$ | | + | | |
| NL: 48 ($H_2O$ + HCHO) | | + | | |
| FI: [18:1;O2–$H_2O$–HCHO+H] | | + | | |
| FI: deoxysphingosine (18:1,O) | | | + | |
| FI: deoxysphingosine (18:1,O) – $H_2O$ | | | + | |
| FI: phytosphingosine (18:0;O3) | | | | + |
| FI: phytosphingosine (18:0;O3) – $H_2O$ | | | | + |
| FI: phytosphingosine (18:0;O3) – $2xH_2O$ | | | | + |
| FI: phytosphingosine (18:0;O3) – $2xH_2O$ | | | | + |
| FI: fatty acyl amide $[FA\text{-}OH+NH_3]^+$ | + | + | + | + |

**DihydroCer:**

- Water loss from precursor
- 2 x water loss from precursor
- Protonated sphinganine base
- Protonated sphinganine base - water
- Protonated sphinganine base - 2 x water
- Fatty acyl amide fragment

**Cer**

- Water loss from precursor
- 2 x water loss from precursor
- Water loss - formaldehyde loss from precursor
- Protonated sphigosine base - water - formaldehyde
- Protonated sphingosine base
- Protonated sphingosine base - water
- Protonated sphingosine base - 2 x water
- Fatty acyl amide fragment

**DeoxyCer**

- Water loss from precursor
- Protonated deoxysphingosine base
- Protonated deoxysphingosine - water
- Fatty acyl amide fragment

**PhytoCer**

- Water loss from precursor
- 2 x water loss from precursor
- Protonated phytosphingosine base
- Protonated phytosphingosine base - water
- Protonated phytosphingosine base - 2 x water
- Protonated phytosphingosine base - 3 x water
- Fatty acyl amide fragment

#### 9.3 ISF induced false positive identification

[M-$H_2O$+H]$^+$ and [M-2$H_2O$+H]$^+$ ions, usually formed as the result of in source fragmentation (ISF) of [M+H]$^+$ by the collisional decomposition of protonated Cer at the intermediate region of the instrument between the atmospheric pressure of the ESI source and the vacuum parts of the analyzer, were relatively high and significantly complicated accurate identification. To exclude false positive identifications (e.g. Cer 38:3;O instead of Cer 38:2;O2), RT of Cer different ionization states ([M+H]$^+$, [M+Na]$^+$, [M-$H_2O$+H]$^+$, and [M+HCOO]$^-$) were compared (Figure 9.5). When potentially identified Cer 38:3;O had the same RT as Cer 38:2;O2 [M+H]$^+$, [M+Na]$^+$, and [M+HCOO]$^-$ ions, it was assigned as [M-$H_2O$+H]$^+$ ion of Cer 38:2;O2, and Cer 38:3;O was excluded from identification list.

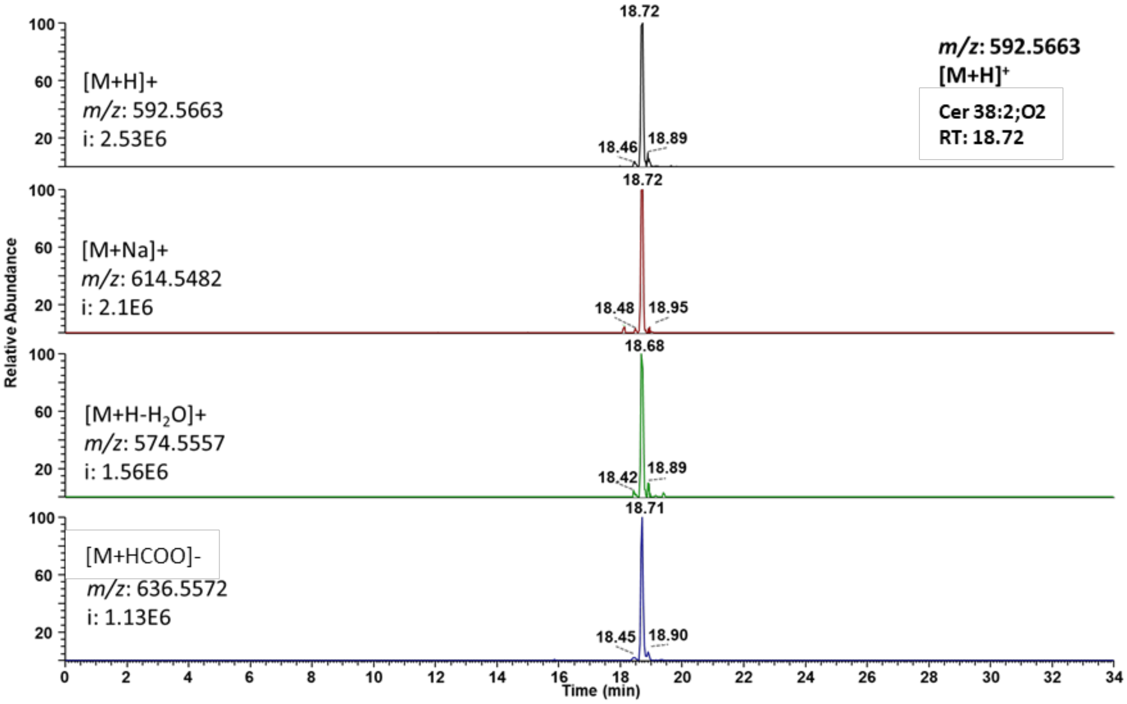


**Figure 9.5.** Filtering false positive identifications arising from ISF of [M+H]$^+$ of Cer. The *m/z* 574.5557 at RT 18.7 min was potentially identified as [M+H]$^+$ of Cer 38:3,O. However, signal at *m/z* 574.5557 co-eluted with [M+H]$^+$, [M+Na]$^+$, and [M+HCOO]$^-$ ions of Cer 38:2;O2, and thus was assigned as Cer 38:2;O2 [M-$H_2O$+H]$^+$ ion.

##### 9.4. RT mapping

To establish retention time rules, molecular Cer species with differing numbers of oxygen atoms were used. Here, structure – RT relationships for all identified Cer species were visualized by plotting Kendrick mass defect by hydrogen (KMD(H)) vs RT plot to control identification accuracy of Cer with different number of hydroxyl groups (1 OH = deoxyCer, 2 OH = dihydroCer/Cer, 3 OH = phytoCer) (Figure 9.6-9.8). All species falling out of the diagonal (different carbon number but the same DBE) and horizontal (same carbon number but different DBE) trend lines were excluded.

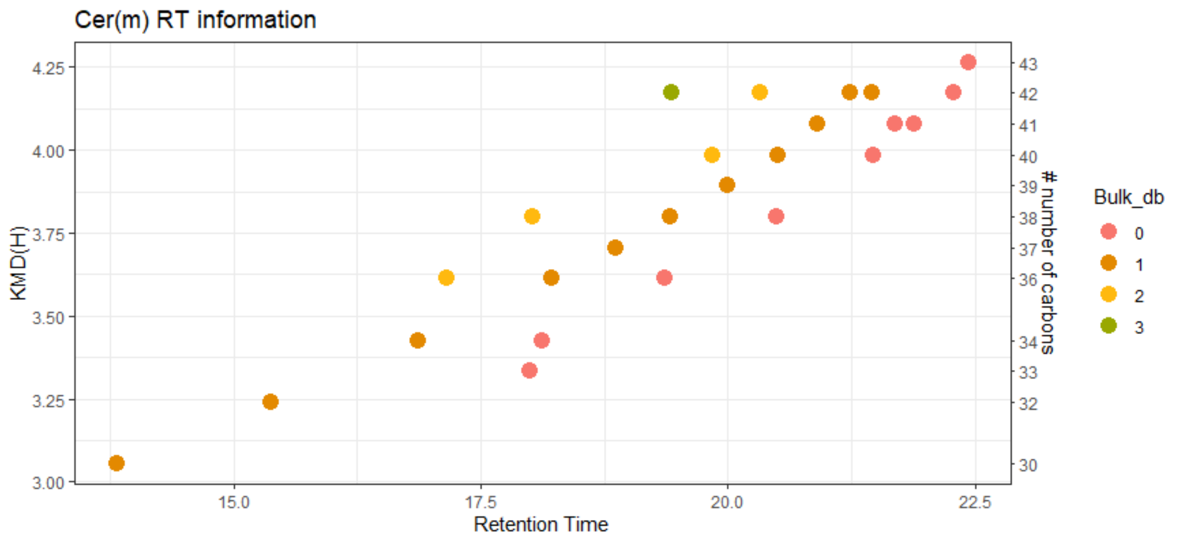


**Figure 9.6**. KMD(H), Cer carbohydrate chain carbon number vs RT plot for all Cer species containing one hydroxyl group (deoxyCer). Symbols color represents the number of double bounds in carbohydrate chains.

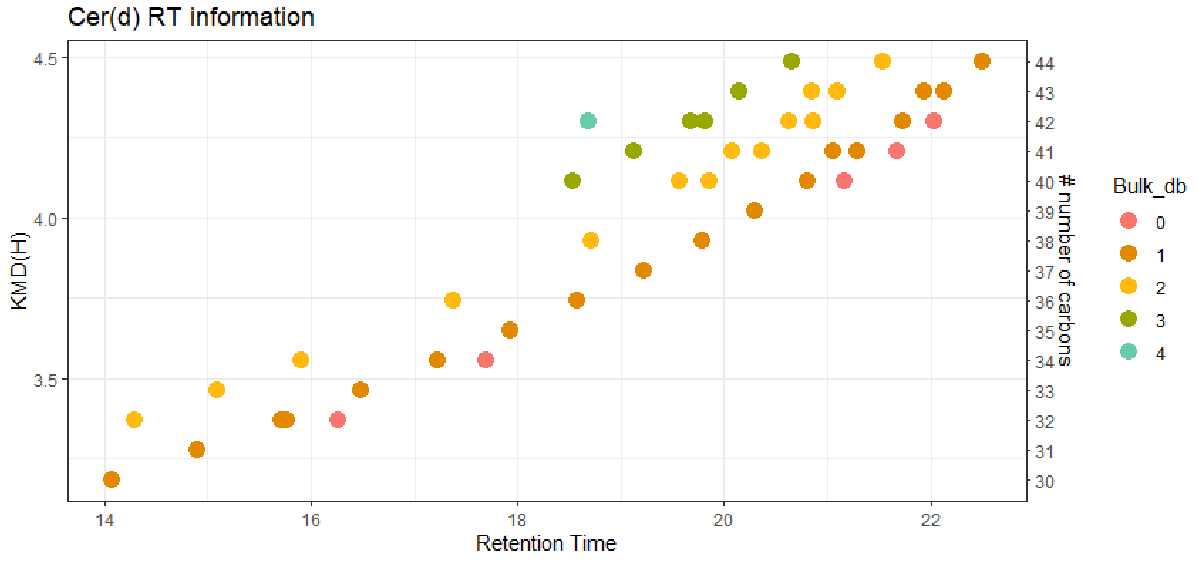


**Figure 9.7**. KMD(H), Cer carbohydrate chain carbon number vs RT plot for all Cer species containing two hydroxyl group (dihydroCer/Cer). Symbols color represents the number of double bounds in carbohydrate chains.

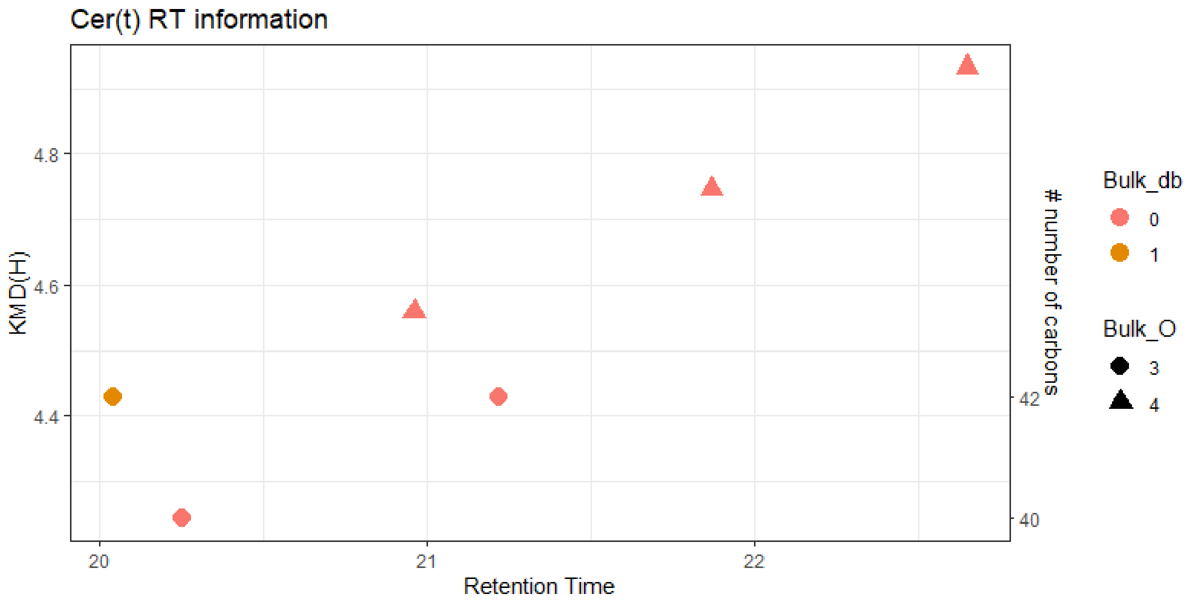


**Figure 9.7**. KMD(H), Cer carbohydrate chain carbon number vs RT plot for all Cer species containing three hydroxyl group (phytoCer). Symbols color represents the number of double bounds in carbohydrate chains.

#### 10. Manual confirmation of identities for hexosylated ceramide (HexCer) lipid molecular species:

Software assisted identification of HexCer molecular species required further manual confirmation to assure accurate identification of HexCer subclasses (HexCer, Hex2Cer, Hex3Cer) based on specific fragment ions and retention time mapping. HexCer subclasses contain varying number of hexoses bound to the ceramide scaffold, i.e. one hexose (HexCer), two hexoses (Hex2Cer) and three hexoses (Hex3Cer). The identification of Cer subclasses within HexCer followed the same logic as for Cer.

##### 10.1. Monitored adducts

HexCer were monitored as protonated adducts in positive mode.

##### 10.2. Fragmentation patterns

General fragmentation pattern and representative MS/MS spectra for HexCer ionized in positive mode are illustrated below:

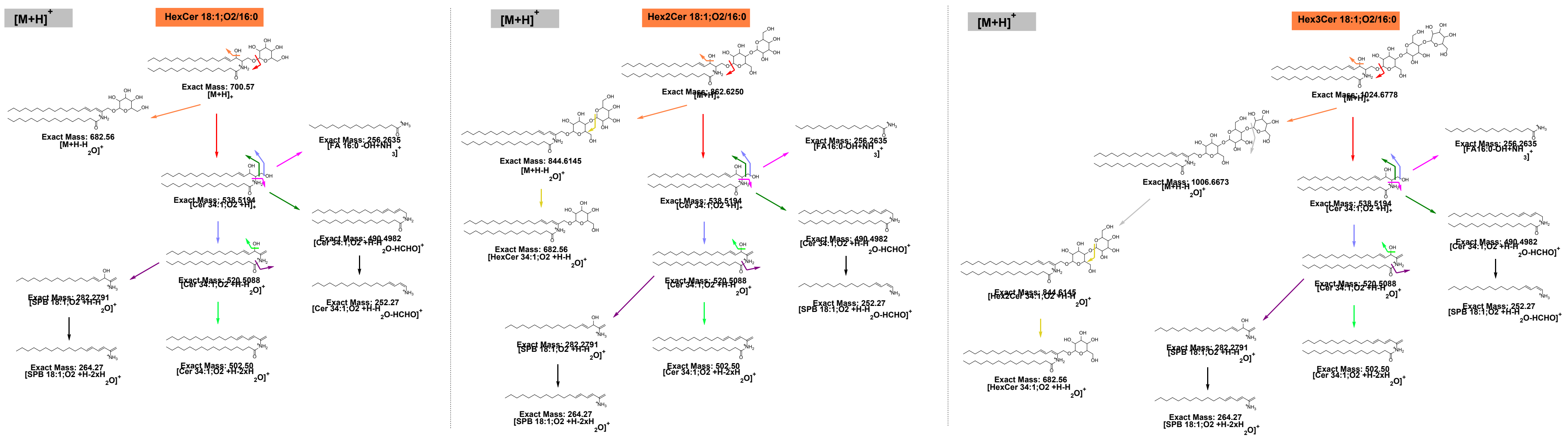


**Scheme 10.1.** General HCD fragmentation pattern of protonated HexCer (HexCer 18:1;O2/16:0), Hex2Cer (Hex2Cer 18:1;O2/16:0) and Hex3Cer (Hex3Cer 18:1;O2/16:0) as $[M+H]^+$.

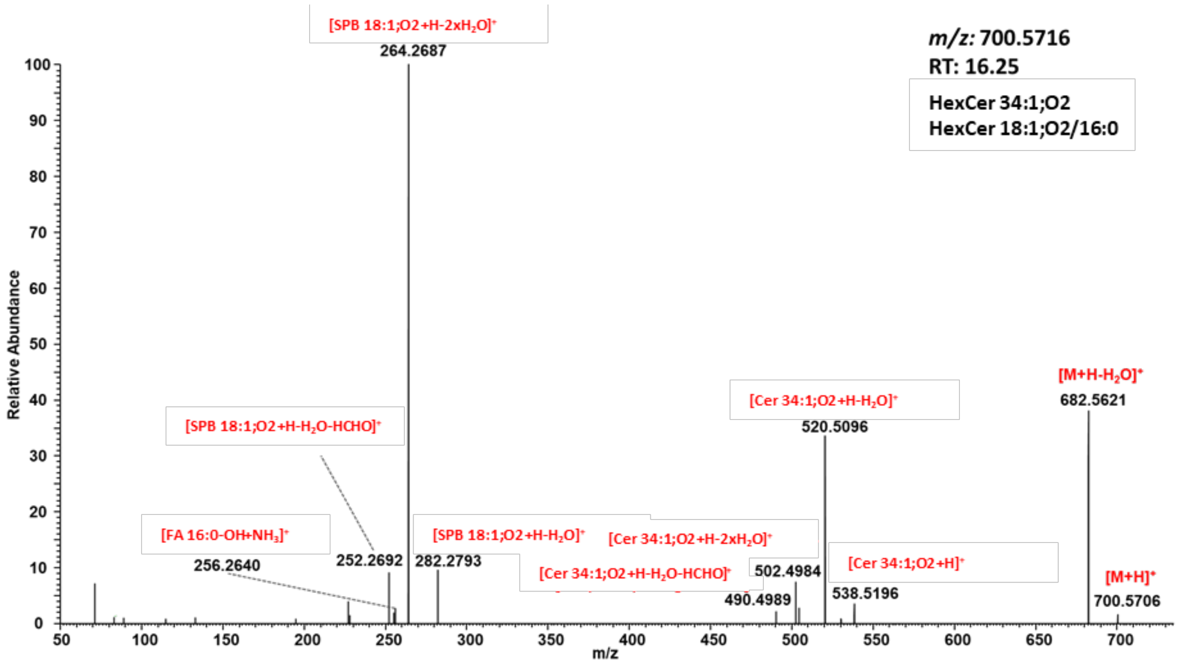


**Figure 10.1.** Representative HCD spectra of HexCer 18:1;O2/16:0 as protonated adduct $[M+H]^+$.

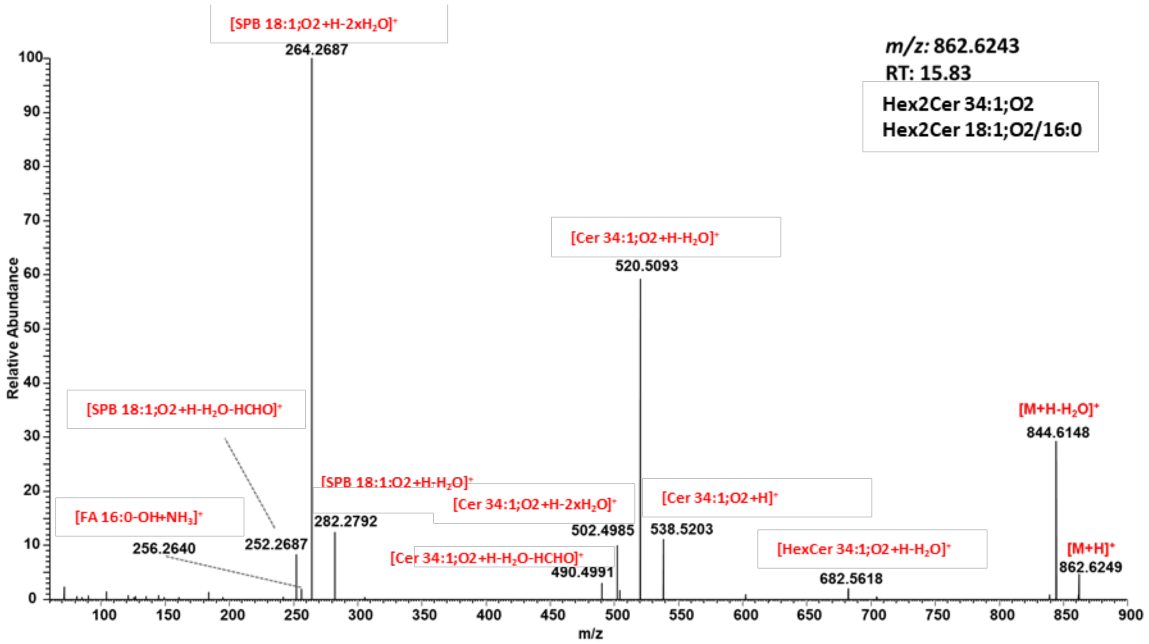


**Figure 10.2.** Representative HCD spectra of Hex2Cer 18:1;O2/16:0 as protonated adduct $[M+H]^+$.

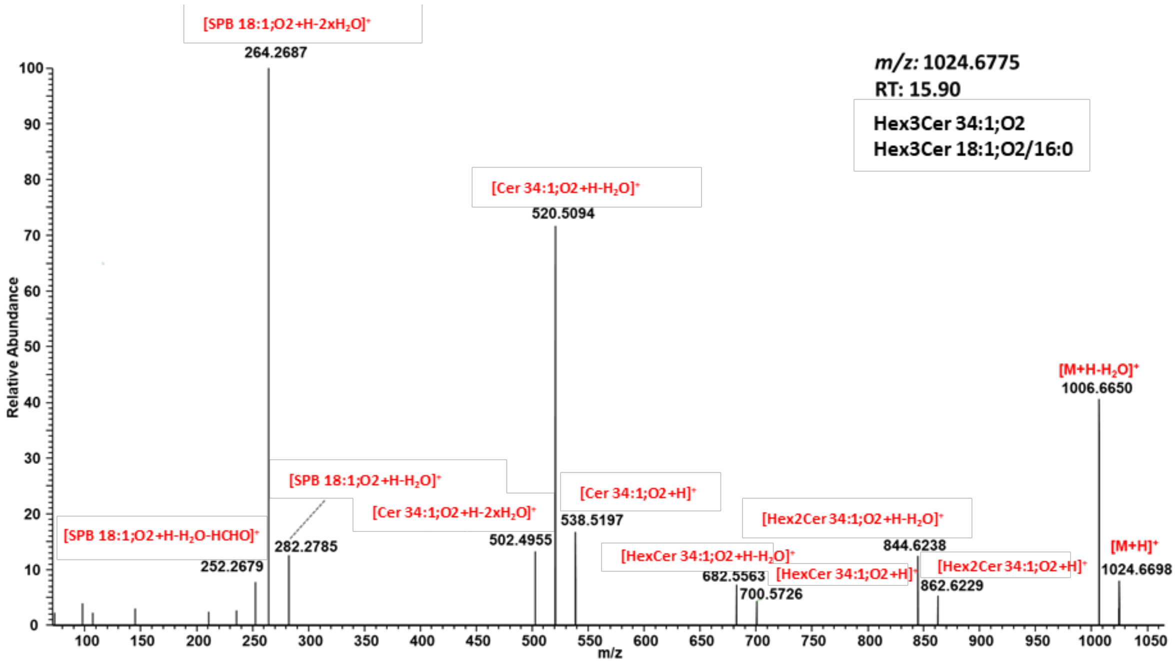


**Figure 10.3.** Representative HCD spectra of Hex3Cer 18:1;O2/16:0 as protonated adduct $[M+H]^+$.

**Table 10.1.** Summary of Cer subclass specific neutral loss (NL) and fragment ions (FI) obtained by positive HCD.

| NL/FI | HexCer | Hex2Cer | Hex3Cer |
|---|---|---|---|
| NL: 18 ($H_2O$) | + | + | + |
| NL: 180 (hexose) | | + | + |
| NL: 162 (hexose-$H_2O$) | + | | |
| NL: 342 (hexose + (hexose-$H_2O$)) | | | + |
| NL: 324 (2x(hexose-$H_2O$)) | | + | |
| NL: 486 (3x(hexose-$H_2O$)) | | | + |

**HexCer:**

- Water loss from precursor
- (hexose-$H_2O$) loss from precursor

**Hex2Cer**

- Water loss from precursor
- Hexose loss from precursor
- 2x(Hexose – water) loss from precursor

**Hex3Cer**

- Water loss from precursor
- Hexose loss from precursor
- Hexose loss + (Hexose-water) loss from precursor
- 3x(Hexose – water) loss from precursor

#### 10.3. RT mapping

To establish retention time rules, molecular HexCer species with differing numbers attached hexoses were used. Here, structure – RT relationships for all identified HexCer species were visualized by plotting Kendrick mass defect by hydrogen (KMD(H)) vs RT plot to control identification accuracy (Figure 10.4-10.6). All species falling out of the diagonal (different carbon number but the same DBE) and horizontal (same carbon number but different DBE) trend lines were excluded.

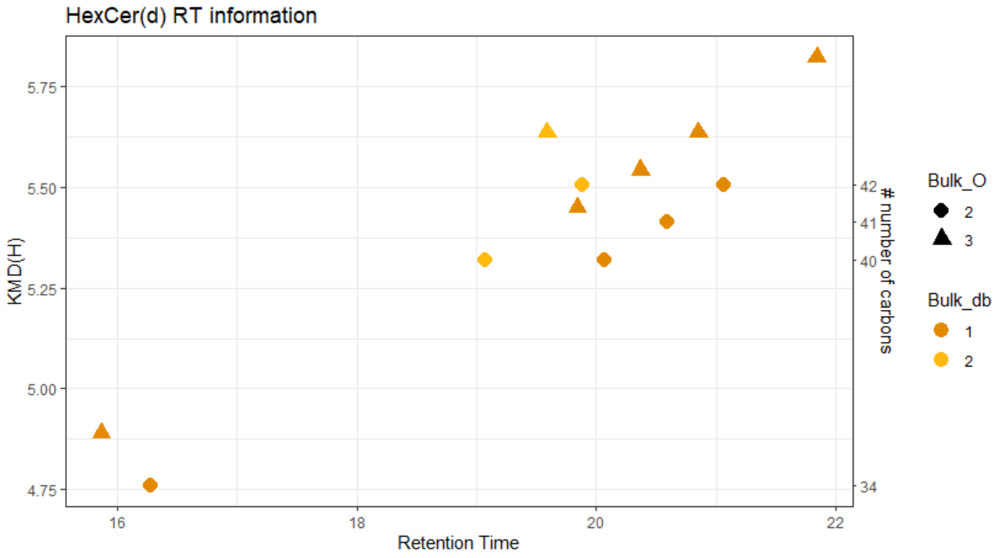


**Figure 10.4**. KMD(H), HexCer carbohydrate chain carbon number vs RT plot for all HexCer species containing one hexose. Symbols color represents the number of double bounds in carbohydrate chains.

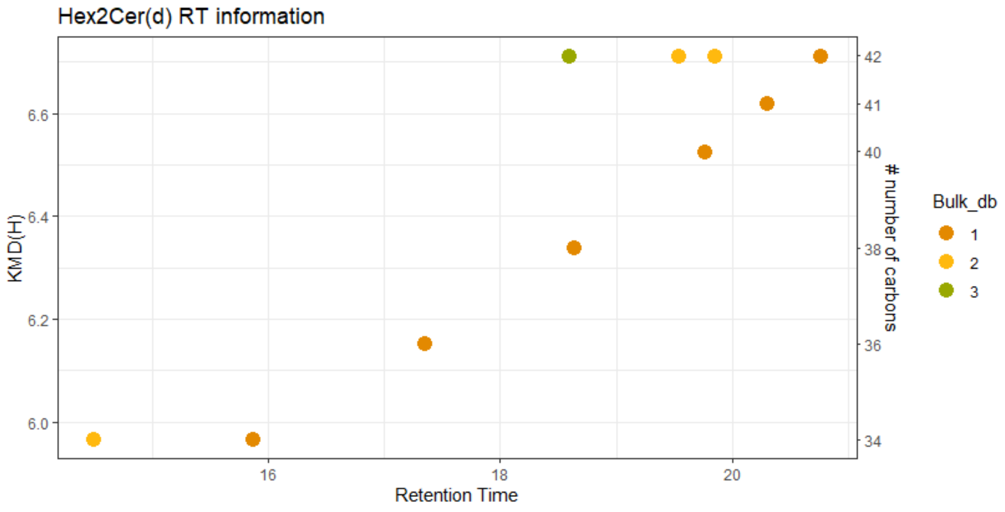


**Figure 10.5**. KMD(H), Hex2Cer carbohydrate chain carbon number vs RT plot for all HexCer species containing two hexoses. Symbols color represents the number of double bounds in carbohydrate chains.

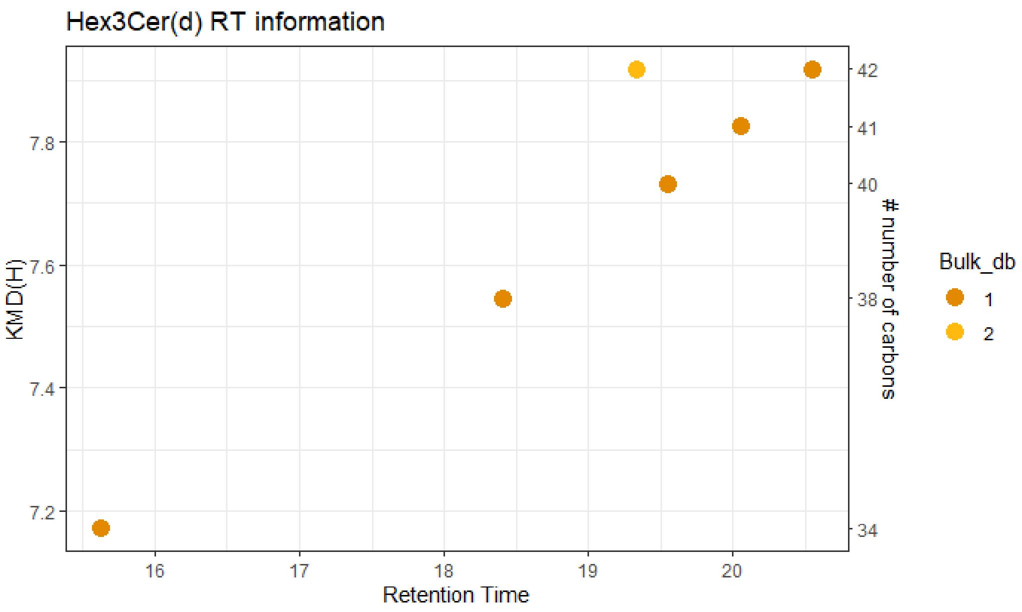


**Figure 10.6**. KMD(H), Hex3Cer carbohydrate chain carbon number vs RT plot for all HexCer species containing three hexoses. Symbols color represents the number of double bounds in carbohydrate chains.

#### Acyl Carnitines (CAR)

##### 11. Manual confirmation of identities for CAR lipid molecular species.

CAR molecular species were identified manually and to assure accurate identification the presence of CAR specific fragment ion was determined and retention time mapping was performed.

##### 11.1. Monitored adducts

CAR were monitored as protonated adducts in the positive ion mode.

##### 11.1. Fragmentation patterns

General fragmentation pattern and representative MS/MS spectra for CAR ionized in positive mode are illustrated below:

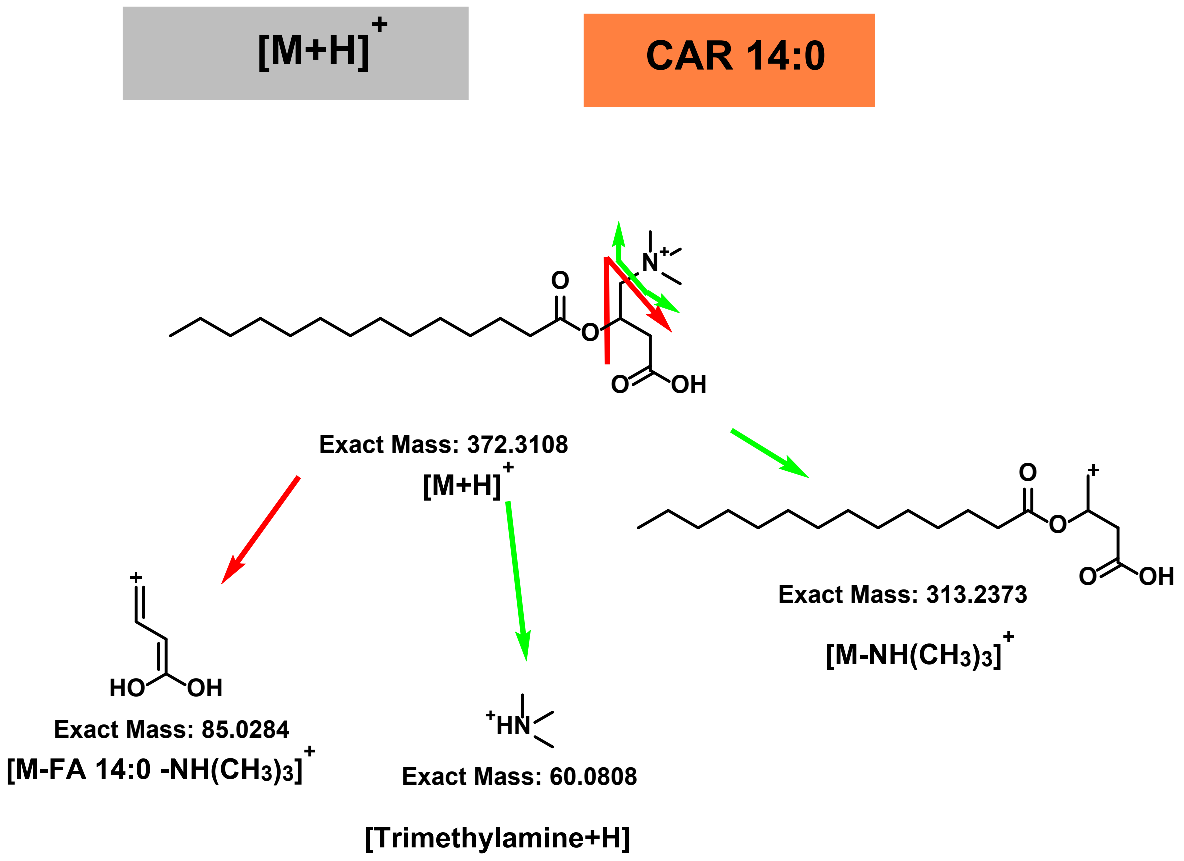


**Scheme 11.1.** General HCD fragmentation pattern of protonated $[M+H]^+$ adducts of CAR.

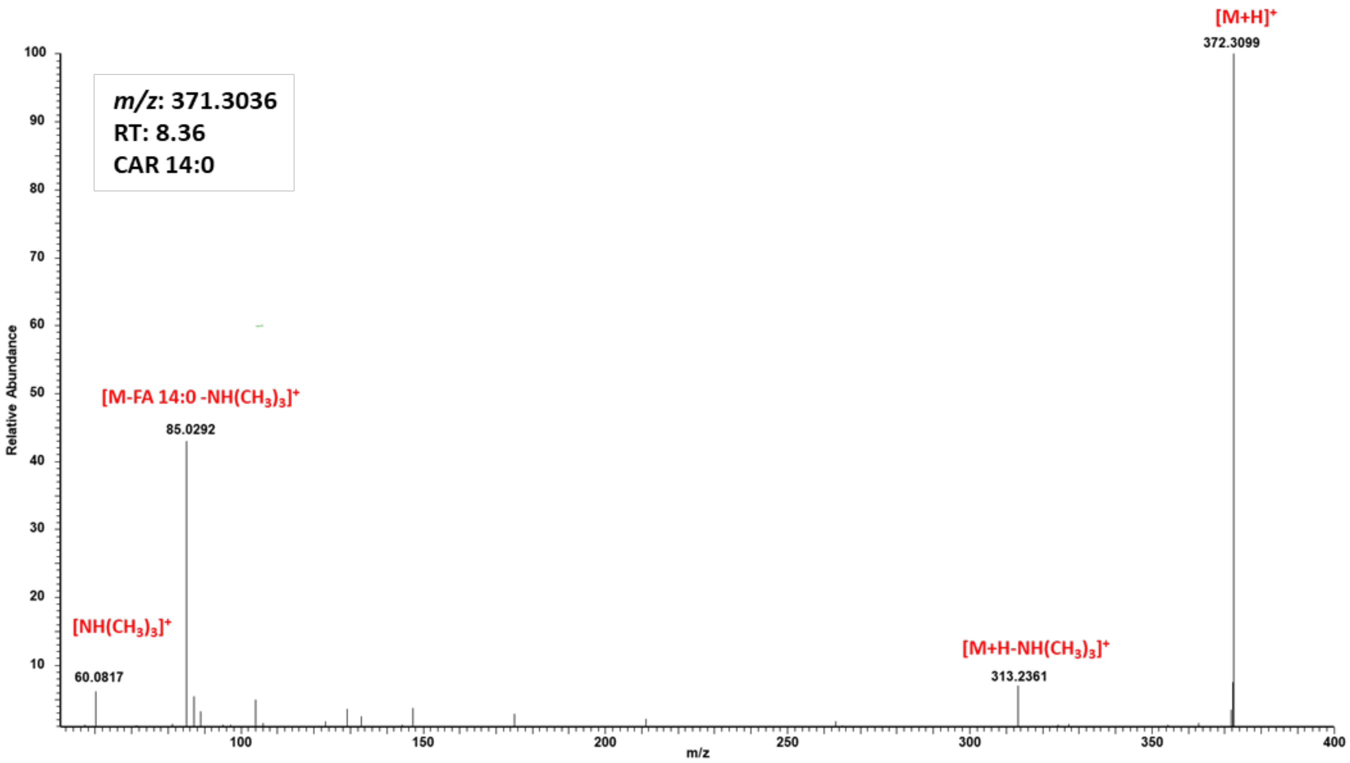


**Figure 11.1.** Representative HCD spectrum of the protonated adduct $[M+H]^+$ of CAR 14:0.

**Table 11.1.** Summary of CAR specific neutral loss and fragment ions obtained by positive ion mode HCD.

| **Positive ion mode HCD** | |
|---|---|
| NL: 59 (choline) | + |
| NL: fatty acid + choline | + |
| FI: 60 (choline) | + |

#### 11.3. RT mapping

Finally, structure – RT relationships for all identified CAR species were visualized by plotting Kendrick mass defect by hydrogen (KMD(H)) vs RT plot to control identification accuracy (Figure 11.2). All species falling out of the diagonal (different carbon number but the same DBE) and horizontal (same carbon number but different DBE) trend lines were excluded.

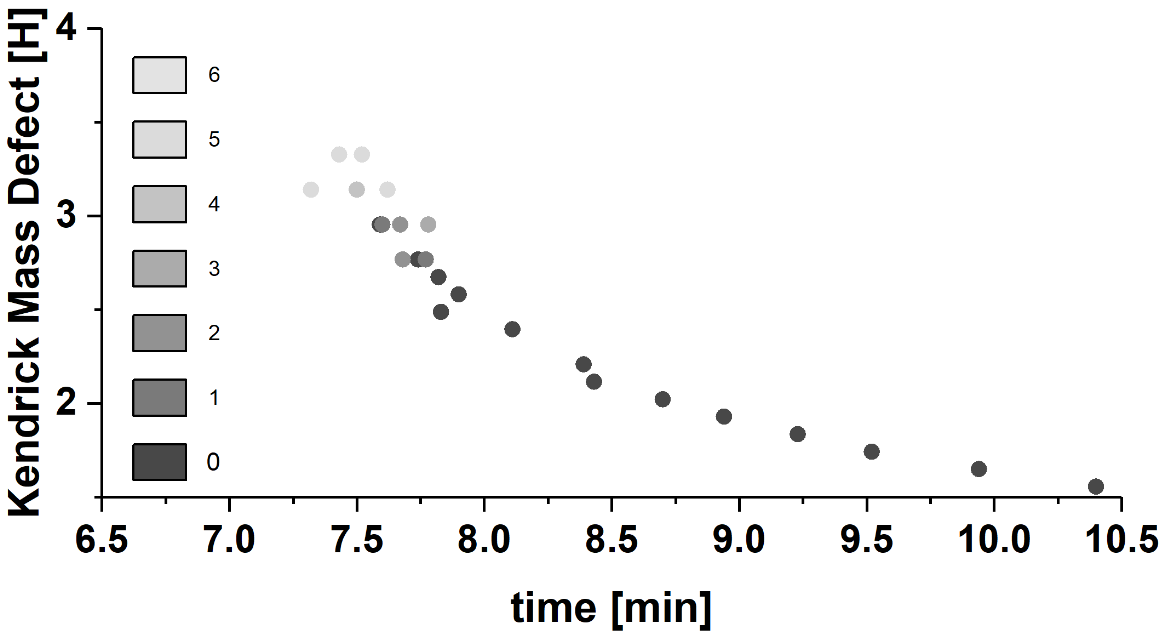


**Figure 11.2**. KMD(H), CAR carbohydrate chain carbon number vs RT plot for all identified CAR species. Symbols color represents the number of double bounds in carbohydrate chains.

#### Glycerolipids

##### 12. Manual confirmation of identities for DG lipid molecular species.

Software assisted identification of DG molecular species required further manual confirmation to assure accurate identification based on specific fragment ions and retention time mapping.

###### 12.1. Monitored adducts

DG were monitored as ammoniated ions in positive ion modes.

###### 12.2. Fragmentation patterns

General fragmentation pattern and representative MS/MS spectra for DG ionized in positive mode are illustrated below:

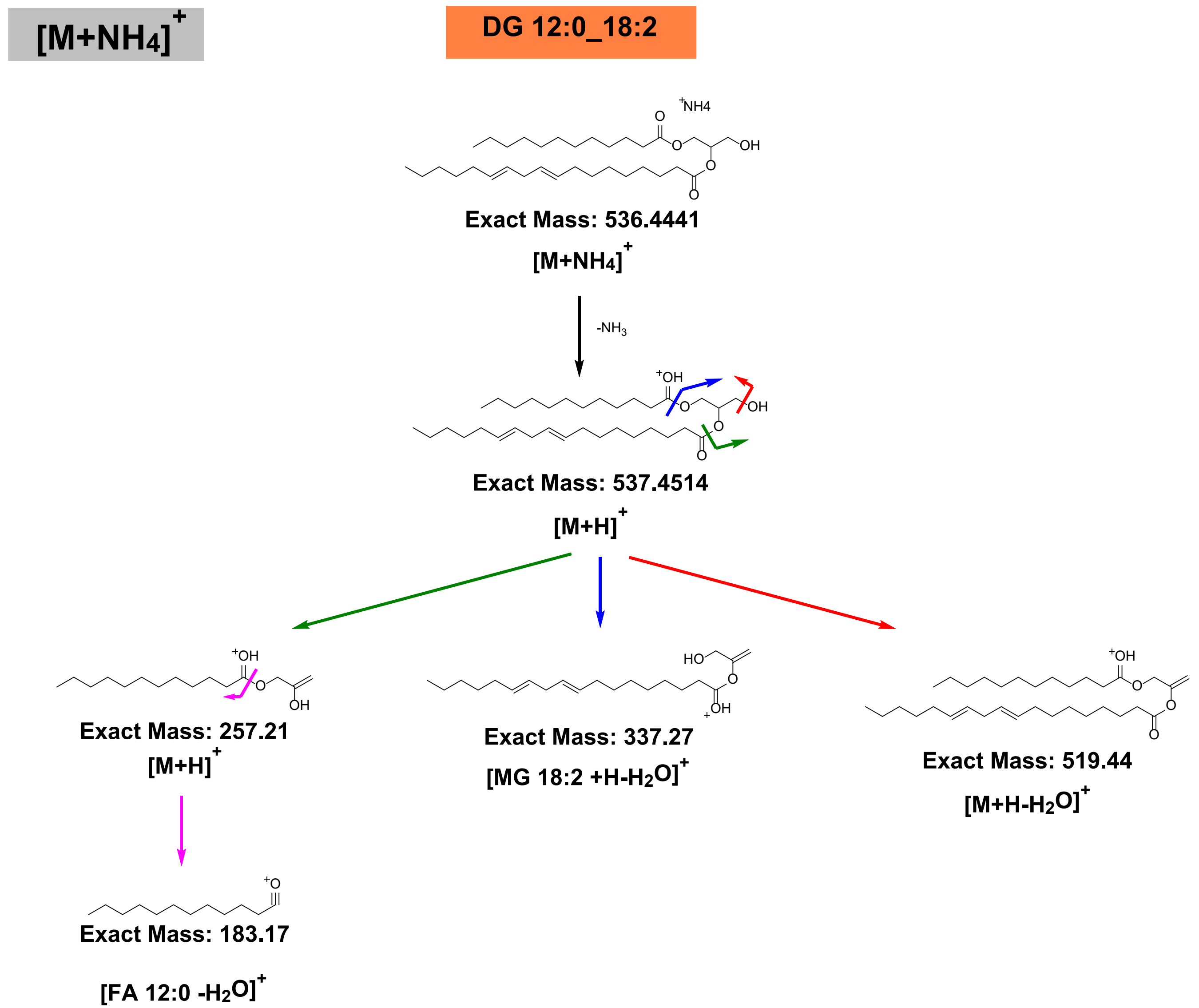


**Scheme 12.1.** General HCD fragmentation pattern of protonated $[M+ NH_4]^+$ adducts of DG.

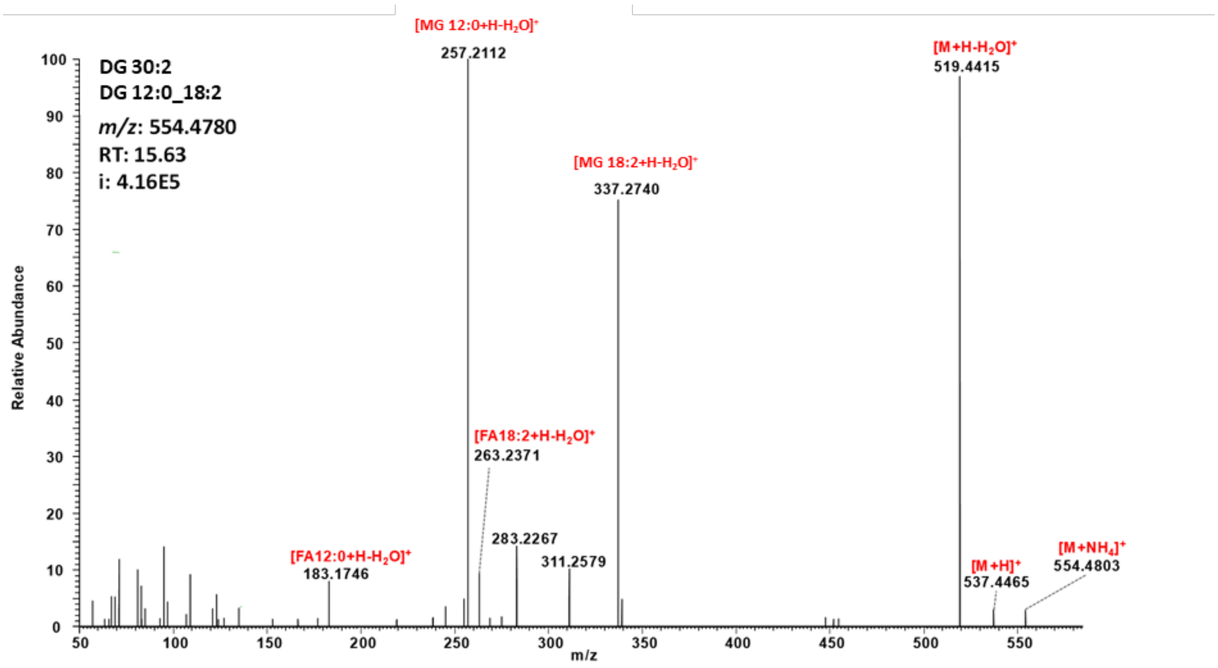


**Figure 12.1.** Representative HCD spectrum of the protonated adduct $[M+NH_4]^+$ of DG 12:0_18:2.

**Table 12.1.** Summary of DG specific neutral loss and fragment ions obtained by positive ion mode HCD.

| **Positive ion mode HCD** | |
|---|---|
| NL: 17 (ammonia) | + |
| NL: 18 (water loss) | + |
| FI: fatty acyl 1 monoglycerol – $H_2O$ | + |
| FI: fatty acyl 1 oxonium ion | + |
| FI: fatty acyl 2 oxonium ion | + |

##### 12.3. RT mapping

Finally, structure – RT relationships for all identified DG species were visualized by plotting Kendrick mass defect by hydrogen (KMD(H)) vs RT plot to control identification accuracy (Figure 12.2). All species falling out of the diagonal (different carbon number but the same DBE) and horizontal (same carbon number but different DBE) trend lines were excluded.

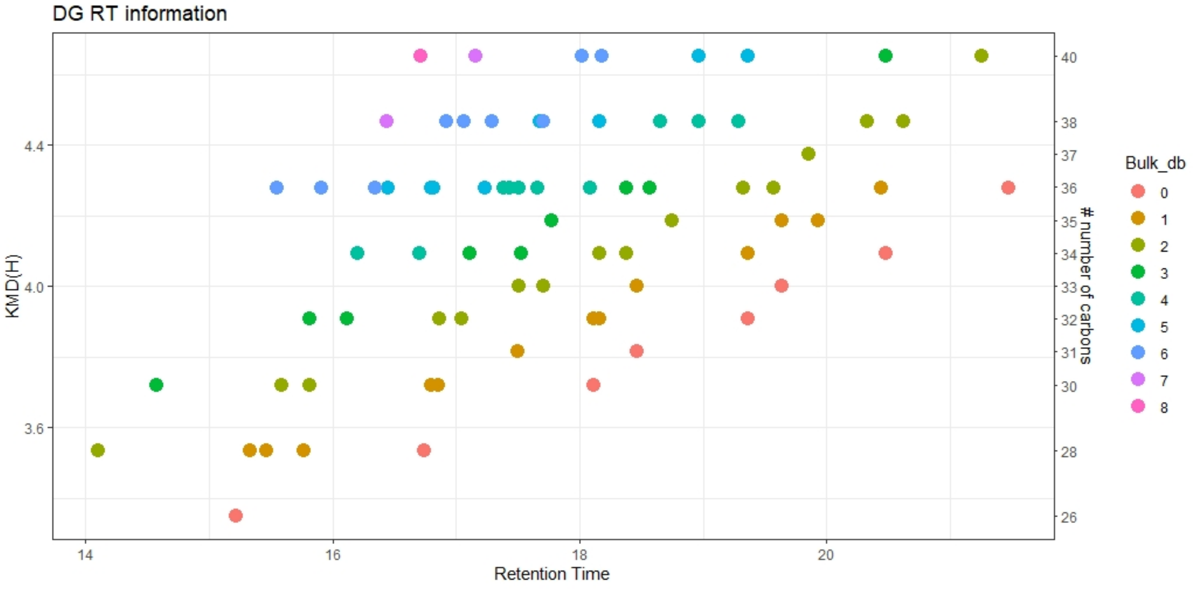


**Figure 12.2**. KMD(H), DG carbohydrate chain carbon number vs RT plot for all identified DG species. Symbols color represents the number of double bounds in carbohydrate chains.

##### 13. Manual confirmation of identities for TG lipid molecular species.

Software assisted identification of TG molecular species required further manual confirmation to assure accurate identification based on specific fragment ions and retention time mapping.

###### 13.1. Monitored adducts

TG were monitored as ammoniated ions in positive ion modes.

###### 13.2. Fragmentation patterns

General fragmentation pattern and representative MS/MS spectra for TG ionized in positive mode are illustrated below:

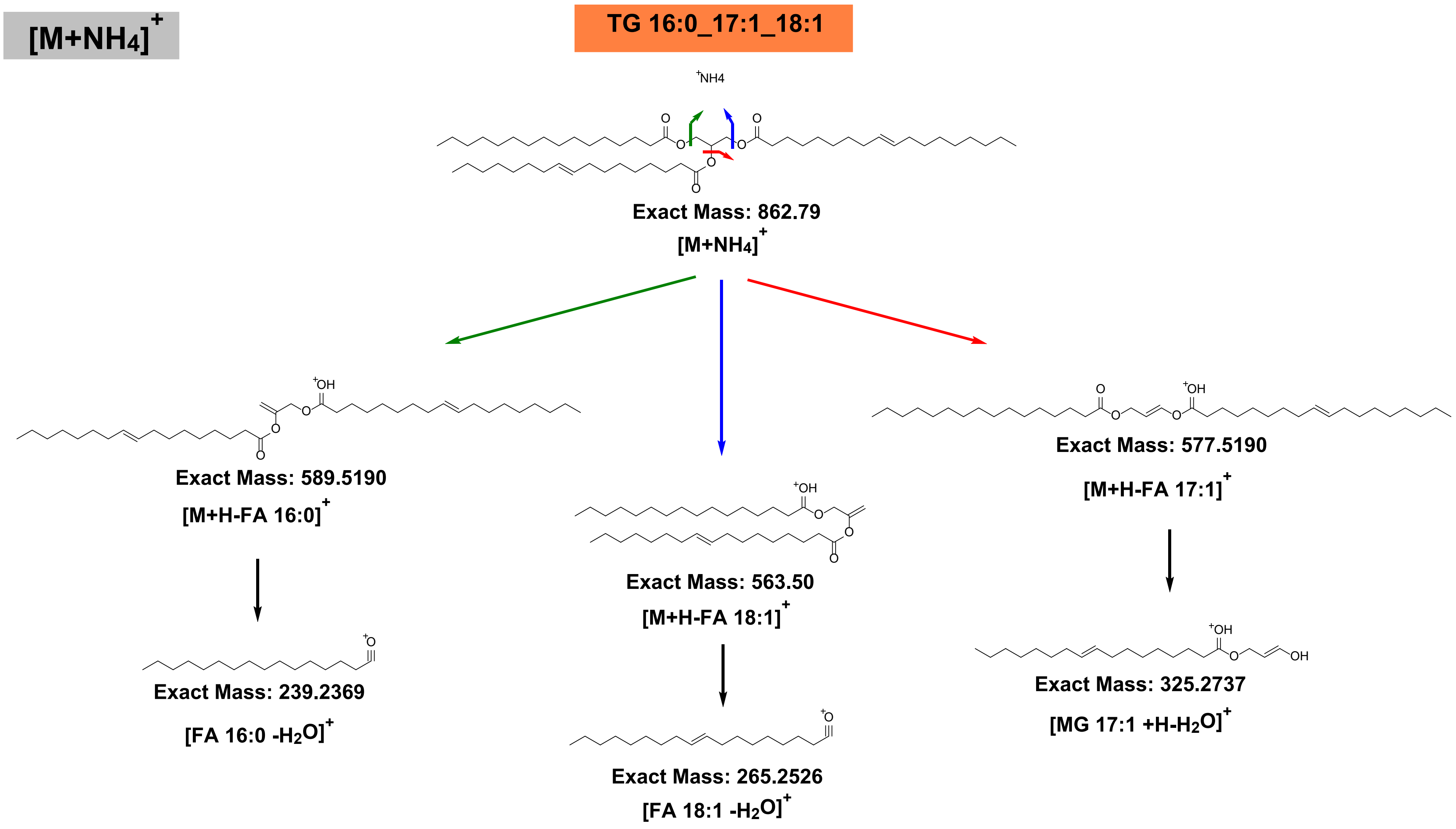


**Scheme 13.1.** General HCD fragmentation pattern of protonated $[M+ NH_4]^+$ adducts of TG.

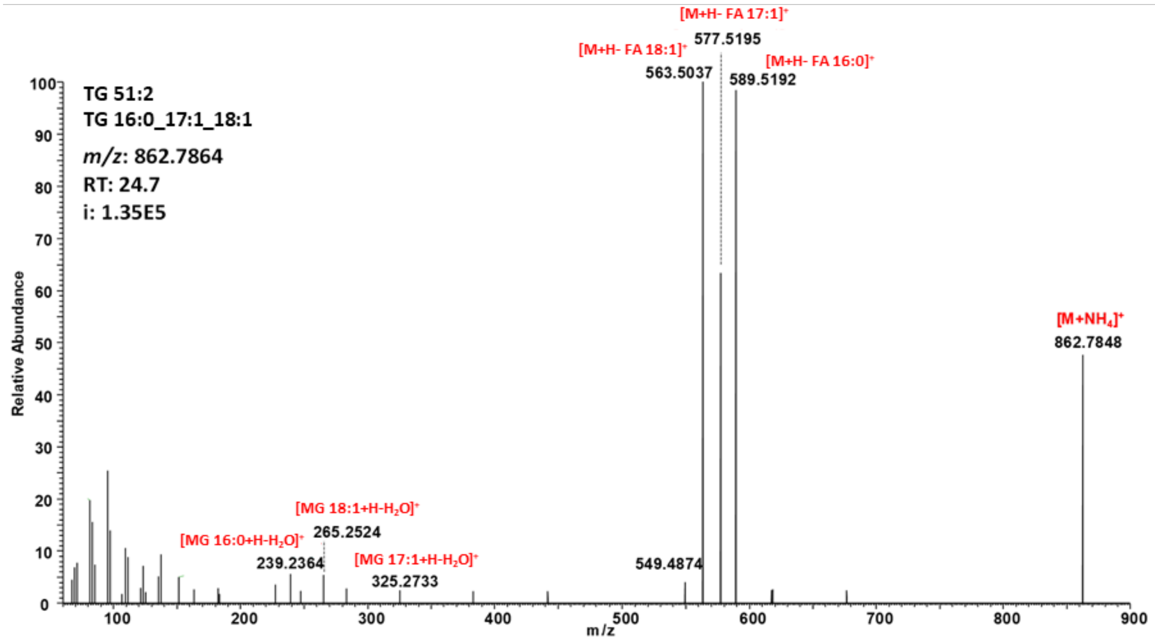


**Figure 13.1.** Representative HCD spectrum of the protonated adduct $[M+NH_4]^+$ of TG 16:0_17:1_18:1.

**Table 13.1.** Summary of TG specific neutral loss and fragment ions obtained by positive ion mode HCD.

| Positive ion mode HCD | |
|---|---|
| NL: 17 (ammonia) | + |
| FI: fatty acyl 1 monoglycerol – $NH_4$ – $H_2O$ | + |
| FI: fatty acyl 2 monoglycerol – $NH_4$ – $H_2O$ | + |
| FI: fatty acyl 3 monoglycerol – $NH_4$ – $H_2O$ | + |
| FI: fatty acyl 1 oxonium ion | + |
| FI: fatty acyl 3 oxonium ion | + |
| FI: fatty acyl 1 monoglycerol –$H_2O$ | + |

##### 13.3. RT mapping

Finally, structure – RT relationships for all identified TG species were visualized by plotting Kendrick mass defect by hydrogen (KMD(H)) vs RT plot to control identification accuracy (Figure 13.2). All species falling out of the diagonal (different carbon number but the same DBE) and horizontal (same carbon number but different DBE) trend lines were excluded.

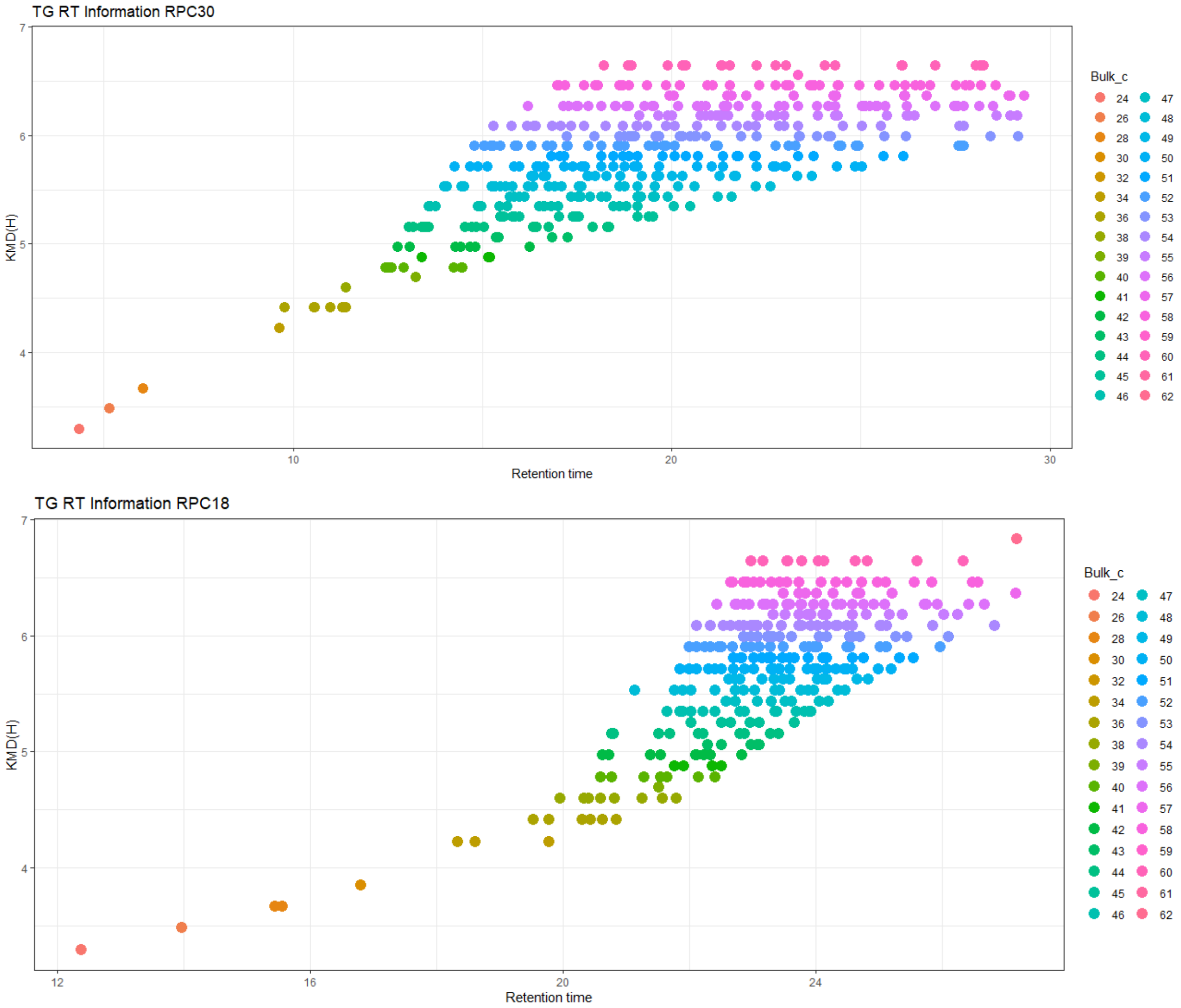


**Figure 13.2**. KMD(H), TG carbohydrate chain carbon number vs RT plot for all identified TG species. Symbols color represents the number of double bounds in carbohydrate chains. KMD(H) vs RT plots on RPC18 (upper panel) and RPC30 (lower panel).

#### Cholesteryl Esters

##### 14. Manual confirmation of identities for CE lipid molecular species.

CE molecular species were identified manually and to assure accurate identification the presence of CE specific fragment ion was determined and retention time mapping was performed.

##### 14.1. Monitored adducts

CE were monitored as ammoniated ions in the positive ion mode.

##### 14.1. Fragmentation patterns

General fragmentation pattern and representative MS/MS spectra for CE ionized in positive mode are illustrated below:

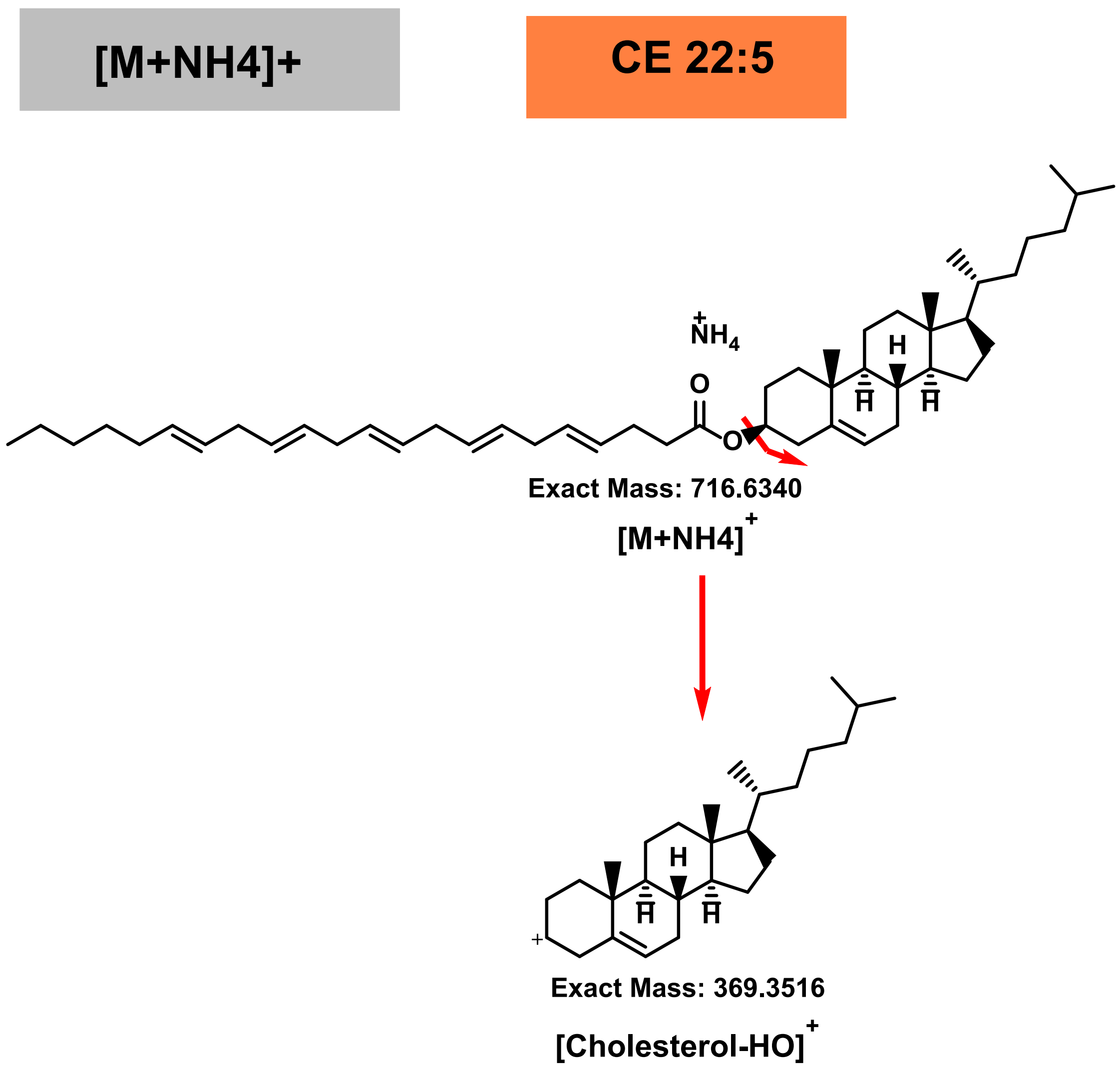


**Scheme 14.1.** General HCD fragmentation pattern of protonated $[M+NH_4]^+$ adducts of CE.

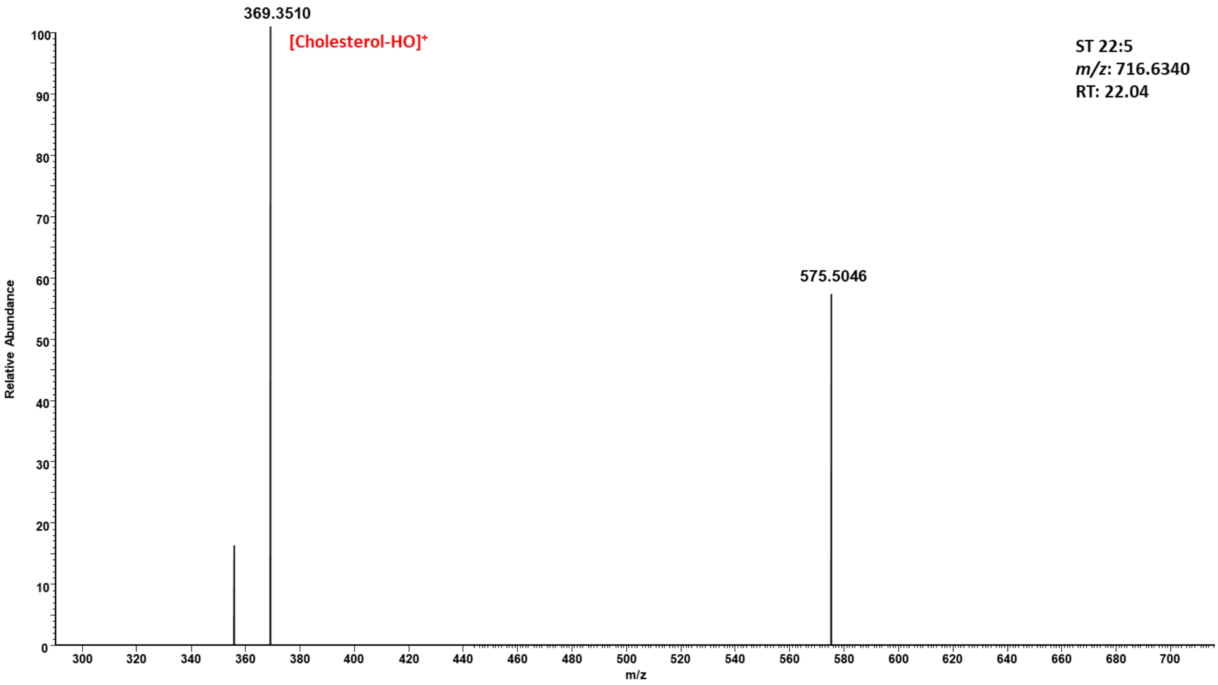


**Figure 14.1.** Representative HCD spectrum of the protonated adduct $[M+NH_4]^+$ of CE 22:5.

**Table 14.1.** Summary of CE specific neutral loss and fragment ions obtained by positive ion mode HCD.

| Positive ion mode HCD | |
|---|---|
| FI: 369 $[cholesterol – HO]^+$ | + |

#### 14.3. RT mapping

Finally, structure – RT relationships for all identified CE species were visualized by plotting Kendrick mass defect by hydrogen (KMD(H)) vs RT plot to control identification accuracy (Figure 14.2). All species falling out of the diagonal (different carbon number but the same DBE) and horizontal (same carbon number but different DBE) trend lines were excluded.

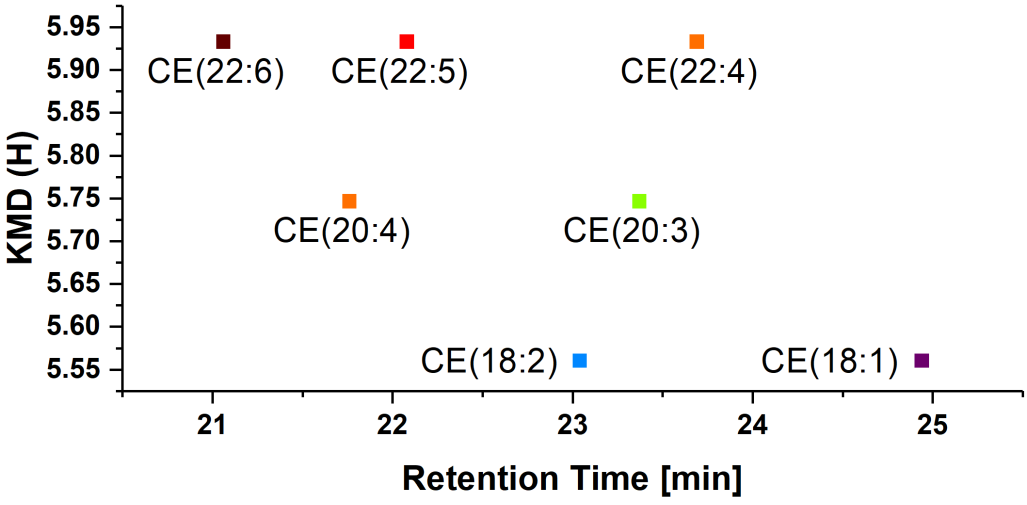


**Figure 14.2**. KMD(H), CE carbohydrate chain carbon number vs RT plot for all identified CE species. Symbols color represents the number of double bounds in carbohydrate chains.
