## Supplementary material for "AdipoAtlas: A Reference Lipidome for Human White Adipose Tissue": Material and Methods

### Materials and Methods

#### 1 Materials

**KEY RESOURCES TABLE**

| REAGENT or RESOURCE | SOURCE | IDENTIFIER |
|---|---|---|
| Lipids | | |
| Cer/Sph Mixture I | Avanti Polar Lipids Inc. | LM6002-1EA |
| Cer 18:0;O3/16:0 | Avanti Polar Lipids Inc. | 860617P |
| Cer 18:0;O3/8:0 | Avanti Polar Lipids Inc. | 860609P |
| Cer 18:0;O2/8:0 | Sigma Aldrich | C8605 |
| Cer 18:0;O2/12:0 | Avanti Polar Lipids Inc. | 860635 |
| Cer 18:1;O2/17:0,O[2R-OH] | Avanti Polar Lipids Inc. | 860817P |
| Cer 18:1;O/6:0 | Cayman Chemical | Cay25493-500 |
| SPLASH® LIPIDOMIX® | Avanti Polar Lipids Inc. | 330707-1EA |
| PC 16:0/18:1 | Avanti Polar Lipids Inc. | 850457 |
| PE 16:0/18:1 | Avanti Polar Lipids Inc. | 850757 |
| LPC 18:1 | Avanti Polar Lipids Inc. | 845875 |
| LPE 18:1 | Avanti Polar Lipids Inc. | 846725 |
| PA 16:0/18:1 | Avanti Polar Lipids Inc. | 840857 |
| PS 16:0/18:1 | Avanti Polar Lipids Inc. | 840034 |
| SM 18:1;O2/18:1 | Avanti Polar Lipids Inc. | 860587 |
| Deuterated Acylcarnitine Mix | EURISO-TOP GmbH | NSK-B-1 |
| FA 18:1 | Sigma Aldrich | O1008 |
| FA 18:0 ([13]C1) (99 atom% 13C) | Sigma Aldrich | 299162 |
| FC | Sigma Aldrich | C8667 |
| CE18:0 | Sigma Aldrich | C79409 |
| MAG Mix – 1 | Larodan Inc. | 90-3001 |
| DG 18:1/18:1/0:0 ([13]C3) | Larodan Inc. | 78-1892-7 |
| DG 16:0/16:0/0:0 | Sigma Aldrich | D9135 |
| TAG Mix – 10 | Larodan Inc. | 90-3010 |
| TG 18:1/18:1/18:1 ([13]C3) | Larodan Inc. | 78-1891-7 |
| TG 16:0/16:0/16:0 ([13]C3) | Larodan Inc. | 79-1600-7 |
| TG Standard Mix – GLC 768 | Nu-Chek Prep Inc. | GLC-768 |
| TG Standard Mix 2 – GLC 406 | Nu-Chek Prep Inc. | GLC-406 |
| TG 20:1/20:1/20:1 | Nu-Chek Prep Inc. | T-270 |
| TG 20:2/20:2/20:2 | Nu-Chek Prep Inc. | T-280 |
| TG 20:3/20:3/20:3 | Nu-Chek Prep Inc. | T-290 |
| TG 20:4/20:4/20:4 | Nu-Chek Prep Inc. | T-295 |
| TG 20:5/20:5/20:5 | Nu-Chek Prep Inc. | T-325 |
| TG 14:1/14:1/14:1 | Nu-Chek Prep Inc. | T-205 |
| TG 16:1/16:1/16:1 | Nu-Chek Prep Inc. | T-215 |
| TG 16:0/16:0/18:1 | Larodan Inc. | 34-1602 |
| Liquid and Thin-layer Chromatography Equipment | | |
| HybridSPE® – Phospholipid, 30 mg/1 ml | Merck KGaA | 55261-U |
| Strata® NH2, 55 µm, 70 Å, 200 mg/3 ml | Phenomenex Inc. | 8B-S009-FBJ |
| Accucore C30 column (150 x 2.1 mm; 2.6 µm, 150 Å) | Thermo Fisher Scientific | 27826-152130 |
| Accucore C18 column (150 x 2.1 mm; 2.6 µm, 150 Å) | Thermo Fisher Scientific | 16126-152130 |
| Acquity UPLC BEH HILIC Si column (100 x 1.0 mm; 1.7 µm, 130 Å) | Waters Corp. | 186003458 |
| HPTLC silica gel plates 60, 20x10 cm | Merck KGaA | 1.05633.0001 |
| Solvents and Additives | | |

| | | |
|---|---|---|
| Acetonitrile (ULC/MS-CC/SFC grade) | Biosolve | 0001204102BS |
| 2-Propanol (*i*-PrOH) (ULC/MS-CC/SFC grade) | Biosolve | 0016264102BS |
| Methanol (ULC/MS-CC/SFC grade) | Biosolve | 0013684102BS |
| Formic acid (ULC/MS-CC/SFC grade) | Biosolve | 00069141A8BS |
| Chloroform (Emsure®) | Sigma Aldrich | 1024451000 |
| Methyl-tert-butyl-ether (≥99%) | Sigma Aldrich | 34875 |
| Ammonium formate (MS grade) | Sigma Aldrich | 70221 |
| Ammonium acetate (MS grade) | Sigma Aldrich | 73594 |
| Ethanol (Rotisolv®) | Carl Roth GmbH+Co. KG | P076.1 |
| Acetone (≥99.9%) | Carl Roth GmbH+Co. KG | KK40.1 |
| n-hexane (Rotisolv®, HPLC) | Carl Roth GmbH+Co. KG | 7339.2 |
| Acetic acid (100%, p.a.) | Carl Roth GmbH+Co. KG | 3738.2 |
| Software and Algorithms | | |
| LipidHunter | (Ni et al., 2017) | github.com/SysMedOs/lipidhunter |
| LipoStar | (Goracci et al., 2017) | moldiscovery.com/software/lipostar/ |
| LipidSearch™ | Thermo Fisher Inc. | IQLAAEGABSFAPCMBFK |
| Merging identification lists from various software tools | This paper | https://github.com/SysMedOs/AdipoAtlasScripts |
| OriginPro 2017 | OriginLab Corp. | originlab.com/2017 |
| Graphpad Prism Version 5.02 | GraphPad Software | graphpad.com |
| Metaboanalyst | (Chong et al., 2019) | metaboanalyst.ca/ |
| Deposited Data | | |
| Raw data | Raw data | massive.ucsd.edu (MSV000086729) |
| Biological Samples | | |
| Visceral and subcutaneous white adipose tissue biopsies of lean and obese patients | Leipzig Obesity BioBank | NA |

#### 2 Human white adipose tissue samples

Samples of human white adipose tissue from a total of 86 donors were kindly provided by Matthias Blüher as a part of Leipzig Obesity BioBank. Tissue collection was approved by the Ethics committee of the University of Leipzig (approval number: 159-12-21052012) and all subjects gave written informed consent before taking part in the study. Removed tissue samples were flash frozen in liquid nitrogen and stored at -80°C until further analysis. For the purpose of this study, we included adipose tissue samples from abdominal visceral (VAT) and subcutaneous (SAT) fat depots of lean (BMI < 25kg/m²; n = 5) and obese (BMI > 40kg/m²; n = 81) individuals. Representative tissue pools were generated according to depot and phenotype specificity (Figure 1A).

#### 3 Sample Preparation

75 mg (for workflow optimization) or 50 mg (for LC-MS analysis) of frozen adipose tissue (AT) were cut on ice and collected into Lysing Matrix tubes containing ceramic beads (lysing matrix D, (1/8”), 2 ml, MP Biomedicals, Eschwege, Germany). All the applied solvents were supplemented with 0.1 % (w/v) BHT and extraction was performed on ice. Extraction ratio AT

[mg] / Extraction solvent [mL] was 10. For LC-MS analysis in-house designed WAT Lipid Standards Mixture (100 µL, in $CHCl_3$/MeOH (2:1, v/v); Table 1) was spiked before homogenization.

**Table 1.** Composition of in-house designed WAT Lipid Standards Mixture spiked (100 µL; in $CHCl_3$/MeOH (2:1, v/v)) into 50 mg of AT (in 1 mL of MeOH) before homogenization.

| ISTD Mix Composition | |
|---|---|
| **Lipid** | **Spiked in ≈50 mg WAT [nmol]** |
| **SPLASH® Lipidomix®** | |
| **PC 15:0_18:1 (d7)** | 2.134 |
| **PE 15:0_18:1 (d7)** | 0.080 |
| **PS 15:0_18:1 (d7)** | 0.054 |
| **PG 15:0_18:1 (d7)** | 0.381 |
| **PI 15:0_18:1 (d7)** | 0.107 |
| **PA 15:0_18:1 (d7)** | 0.107 |
| **LPC 18:1 (d7)** | 0.482 |
| **LPE 18:1 (d7)** | 0.109 |
| **CE 18:1 (d7)** | 5.411 |
| **MG 18:1 (d7)** | 0.055 |
| **DG 15:0_18:1 (d7)** | 0.160 |
| **TG 15:0_18:1_15:0 (d7)** | 0.705 |
| **SM d18:1_18:1 (d9)** | 0.419 |
| **Chol (d7)** | 2.499 |
| **Cer/SpH Mix I** | |
| **SPB 17:1;O2** | 0.375 |
| **SPB 17:0;O2** | 0.375 |
| **SPBP 17:1;O2** | 0.375 |
| **SPBP 17:0;O2** | 0.375 |
| **Lac-Cer 18:1;O2/12:0** | 0.375 |
| **Gluc-Cer d18:1;O2/12:0** | 0.375 |
| **SM 18:1;O2/12:0** | 0.375 |
| **Cer 18:1;O2/12:0** | 0.375 |
| **CerP 18:1;O2/12:0** | 0.375 |
| **Cer 18:1;O2/25:0** | 0.375 |
| **Acylcarnitine Mix NSK-B-1** | |
| **Free Carnitine (d9)** | 0.162 |
| **CAR 2:0 (d3)** | 0.037 |
| **CAR 3:0 (d3)** | 0.007 |
| **CAR 4:0 (d3)** | 0.007 |
| **CAR 5:0 (d9)** | 0.007 |
| **CAR 8:0 (d3)** | 0.007 |
| **CAR 14:0 (d9)** | 0.007 |
| **CAR 16:0 (d3)** | 0.015 |
| **Individual Lipid ISTD** | |
| **FA 18:0 ([13]C1)** | 10.000 |
| **TG 18:1/18:1/18:1 ([13]C3)** | 1000.000 |
| **TG 16:0/16:0/16:0 ([13]C3)** | 50.000 |

| | |
|---|---|
| **DG 18:1/18:1/0:0 ([13]C3)** | 10.000 |
| **Cer 18:0;O2/12:0** | 0.375 |
| **Cer 18:0;O2/8:0** | 0.375 |
| **Cer 18:1;O/6:0** | 0.375 |
| **Cer 18:0;O3/16:0** | 0.375 |
| **Cer 18:0;O3/8:0** | 0.375 |
| **Cer 18:1;O2/17:0,O[2*R*-OH]** | 0.375 |

##### 3.1 Lipid extraction

###### 3.1.1 Lipid extraction using Folch method (Iverson et al., 2001)

AT was homogenized in 1 mL of MeOH by FastPrep24$^{TM}$ 5G (3x30s, Lysing Matrix D) with cooling on ice after each homogenization round. Homogenate were transferred into glass tubes (11.5 mL, round bottom culture tubes, VWR) using glass Pasteur pipettes. Beads and lysing tubes were washed with MeOH (400 µL) and $CHCl_3$ (1000 µL), solution was transfer into glass tube. $CHCl_3$ (1.8 mL) was added (to reconstitute ratio $CHCl_3$/MeOH = 2:1, v/v). Mixture was incubated on the roller mixer (4°C, 1 h, 210 rpm), and $H_2O$ (840 µL) was added (to reconstitute ratio $CHCl_3$/MeOH/$H_2O$ = 8:4:3, v/v), incubated on roller mixer (4°C, 10 min, 210 rpm), and centrifuged to achieve phase separation (4°C, 10 min, 2000 x g). ThelLower phase was collected. Extracts were dried *in vacuo* (Eppendorf concentrator 5301, 1 mbar).

###### 3.1.2 Lipid extraction using methyl tert-butyl ether (MTBE) (Matyash et al., 2008)

AT was homogenized in 1 mL of MeOH by FastPrep24$^{TM}$ 5G (3x30s, Lysing Matrix D mode) with cooling on ice after each homogenization round. Homogenate were transferred into glass tubes (11.5 mL, round bottom culture tubes, VWR) using glass Pasteur pipettes. Beads and lysing tubes were washed with MTBE (1000 µL), and the solution was transferred to a glass tube. MTBE (3.95 mL) and MeOH (500 µL) were added (to reconstitute ratio MTBE/MeOH= 3.3:1, v/v). Mixture was incubated on roller mixer (4°C, 1 h, 210 rpm), and $H_2O$ (1240 µL) was added (to reconstitute ratio MTBE/MeOH/$H_2O$ = 3.3:1:0.8, v/v), incubated on roller mixer (4°C, 10 min, 210 rpm), and centrifuged to achieve phase separation (4°C, 10 min, 2000 × g). Upper organic phase with collected with Pasteur pipette. Re-extraction was done by adding MTBE/MeOH/$H_2O$ (1.95 mL; 10:3.3:2.5, v/v) with subsequent incubation on the roller mixer (4°C, 10 min, 210 rpm) and centrifugation (4°C, 10 min, 2000 × g) to achieve phase separation. Organic phases were combined and dried *in vacuo* (Eppendorf concentrator 5301, 1 mbar).

###### 3.1.3 Lipid extraction using Hexane/i-PrOH/HOAc (Slatter et al., 2016)

AT was homogenized in 1 mL of MeOH by FastPrep24$^{TM}$ 5G (3x30s, Lysing Matrix D mode) with cooling on ice after each homogenization round. Homogenate were transferred into glass tubes (11.5 mL, round bottom culture tubes, VWR) using glass Pasteur pipettes. Beads and lysing tubes were washed with hexane (Hex; 1000 µL), solution was transfer to glass tube. Hex (350 µL), i-PrOH (350 µL), HOAc (105 µL; 100%, Roth), and $H_2O$ (1.125 mL) were added to glass tube (to reconstitute ratio HOAc/i-PrOH/Hex = 2:20:30, v/v; Mix/$H_2O$ = 2.5:1, v/v). Mixture was incubated on a roller mixer (4°C, 1 h, 210 rpm), Hex (3.75 mL) was added, incubate on the roller mixer (4°C, 10 min, 210 rpm), and centrifuged to achieve phase separation (4°C, 10 min, 2000 x g). Upper organic phase was collected with Pasteur pipette. Re-extraction was done by adding Hex (3.75 mL) with subsequent incubation on the roller mixer (4°C, 10 min, 210 rpm), and centrifugation (4°C, 10 min, 2000 x g) to achieve phase separation. Organic phases were combined and dried *in vacuo* (Eppendorf concentrator 5301, 1 mbar).

##### 3.1.4 Protein Concentration Determination

Aqueous phases after lipid extraction were dried *in vacuo* (Eppendorf concentrator 5301, 1 mbar), redissolved in the buffer containing 7 M urea, 2 M thiourea, 1 % sodium deoxycholate, 50 mM Tris-HCl, pH 7.5 and protein concentration was determined by Bradford assay. (Bradford, 1976)

#### 3.2 Lipid Fractionation

##### 3.2.1 Liquid-Liquid Extraction (LLE) (GALANOS and KAPOULAS, 1965)

AT lipid extract corresponding to 20 (for workflow optimization) or 45 mg (for LC-MS analysis) of AT was dried in a glass tube (11.5 mL, round bottom culture tubes, VWR). Hex (4.5 mL) and EtOH/$H_2O$ (1.5 mL, 87:13, v/v) were added, tubes were vigorously vortexed (Tube A). Separated EtOH/$H_2O$ phase was transferred into second glass tube containing 4.5 mL Hex (Tube B), vigorously vortexed, and the resulting EtOH/$H_2O$ phase was transferred into an empty glass tube (Tube C). New portion of EtOH/$H_2O$ (1.5 mL, 87:13, v/v) was added to the remaining Hex phase in Tube A, vortexed and transferred to Tube B, vortexed again and transferred to Tube C. This procedure was repeated six more times (in total eight times of EtOH/$H_2O$ extraction from the original Hexane fraction). All collected EtOH/$H_2O$ phases were combined and dried *in vacuo* (Eppendorf concentrator 5301, 1 mbar).

##### 3.2.2 Zr-SPE (Xiaoning, Lu; Michael)

Dry AT lipid extract corresponding to 20 mg of AT dissolved in $CHCl_3$/MeOH (150 µL; 2:1, v/v). Formic acid (0.1% v/v) acidified MeCN (900 µL) was added. Solution was loaded onto dry SPE cartridge (HybridSPE® – Phospholipid, 30 mg/1 ml, Lot: 4800102, Supelco), washed subsequently with formic acid (0.1% v/v) acidified MeCN (1 mL), MeCN (1 mL), and eluted with MeCN containing 5 % (w/v) $NH_4OH$ (2 x 1 mL). Eluates were dried *in vacuo* (Eppendorf concentrator 5301, 1 mbar).

##### 3.2.3 LipFracSPE (based on (Kim and Salem, 1990) with modifications)

Dry AT lipid extract corresponding to 20 mg of AT dissolved in $CHCl_3$/MeOH (300 µL; 2:1, v/v). SPE cartridge (Strata® $NH_2$, 55 µm, 70 Å, 200 mg/3 ml, Lot: S17-003383, Phenomenex) was conditioned with Hexane (6 mL), and equilibrated with $CHCl_3$ (6 mL). AT lipids were loaded, and eluted subsequently with $CHCl_3$/i-PrOH (2:1, v/v, 15 mL, *unpolar fraction*), $Et_2O$/HOAc (100:2, v/v, 5 mL, *fatty acids fraction*), MeOH (5 mL, *neutral phospholipids fraction*) and Hex/i-PrOH/EtOH/0.1 M $NH_4OAc$ (420:350:100:50, v/v + 5 % HOAc (v/v), 5 mL, *acidic phospholipids fraction*). Acidic phospholipids fraction was neutralized by adding $NH_4OH$ (25 %; w/v; 0.75 mL), followed by the addition of $H_2O$ (0.3 mL) and $CHCl_3$ (1 mL). Mixture was centrifuge (3 min, RT, 2000 × g) to induce phase separation, and lower organic phase was collected. Re-extraction was done by adding $CHCl_3$ (2 x 1 mL) with subsequent centrifugation (3 min, RT, 2000 × g). Organic phases were combined and dried *in vacuo* (Eppendorf concentrator 5301, 1 mbar).

#### 4 TLC and NMR based assessment of extraction and fractionation efficiency

##### 4.1 Phospholipid Quantification by $^{31}$P-NMR Spectroscopy (Dannenberger et al., 2010).

Dried lipid extracts were resuspended in buffer containing 200 mM sodium cholate, 5 mM EDTA, 50 mM Tris-HCl, pH 7.65 (500 µL) by vigorous vortexing for 2 min. Samples were placed in 5 mm NMR tubes and $^{31}$P-NMR spectra were recorded on a Bruker DRX-600 spectrometer operating at 242.88 MHz. All measurements were performed using a selective $^{31}$P/$^{1}$H NMR probe at 37°C with composite pulse decoupling (Waltz-16) to eliminate $^{31}$P–$^{1}$H coupling. Pulse intervals of the order of $T_1$ were used to allow quantitative analysis of phospholipid integral intensities. Other NMR parameters were as follows: acquisition time 1 s, data size 8–16 k, 60° pulse, pulse delay 2 s and a line-broadening (LB) of 1 Hz. Chemical shifts were referenced to the resonance of di-lauroyl-phosphatidic acid that was added as concentration and frequency standard. Further details are available in.(Schiller et al., 2007)

##### 4.2 Quantitative high performance thin layer chromatography (qHPTLC) for unpolar lipid quantification

AT lipid extracts were dissolved in $CHCl_3$/MeOH (2:1, v/v), and an amount corresponding to 1-10 µg AT wet weight were loaded using Camag Linomat 5 (Camag, Switzerland) on TLC plates (HPTLC silica gel 60, 20x10 cm, Merck). On each plate 7 dilutions of unpolar lipid TLC standards were loaded for quantitative lipid class specific calibration (Table 2). Plates were developed using hexane/$Et_2O$/HOAc (8:2:1, v/v). Dried TLC plates were immersed in acetone/$H_2O$ (8:2, v/v) containing primuline (0.05 %, w/v) for 5 s (Camag Chromatogramm Immersion Device III). Images were acquired with a CCD camera (Bio-Rad ChemiDoc MP, Bio-Rad) using the primuline fluorescence (Ex: Blue Epi light illumination; Em: Filter 530/28). Densitometric analysis was performed with Image Lab (Version 5.2.1, Bio-Rad).

**Table 2.** Unpolar lipid TLC Standards used for qHPTLC lipid class specific quantification.

| Unpolar Lipid TLC - ISTD | | | | | | | |
|---|---|---|---|---|---|---|---|
| Lipid | Calibration range [nmol] | | | | | | |
| TG 16:0/16:0/18:1 | 36.00 | 18.00 | 7.20 | 3.60 | 1.80 | 0.36 | 0.18 |
| FC | 77.59 | 38.79 | 15.52 | 7.76 | 3.88 | 0.78 | 0.39 |
| DG 16:0/16:0/0 | 52.73 | 26.37 | 10.55 | 5.27 | 2.64 | 0.53 | 0.26 |
| CE 18:0 | 45.79 | 22.90 | 9.16 | 4.58 | 2.29 | 0.46 | 0.23 |
| FA 18:1 | 106.21 | 53.10 | 21.24 | 10.62 | 5.31 | 1.06 | 0.53 |

##### 4.3 Quantitative high performance thin layer chromatography (qHPTLC) for polar lipid quantification

AT lipid extract polar fractions were dissolved in $CHCl_3$/MeOH (2:1, v/v), and amount corresponding to 1-5 mg AT wet weight were loaded using Camag Linomat 5 (Camag, Switzerland) on TLC plates (HPTLC silica gel 60, 20x10 cm, Merck). On each plate 7 dilutions of polar lipid TLC standards were loaded for quantitative lipid class specific calibration (Table 3). Plates were developed using $CHCl_3$/EtOH/TEA/$H_2O$ (5:5:5:1, v/v). Dried TLC plates were immersed in acetone/$H_2O$ (8:2, v/v) containing primuline (0.05 %, w/v) for 5 s (Camag Chromatogramm Immersion Device III). Images were acquired with a CCD camera (Bio-Rad

ChemiDoc MP, Bio-Rad) using the primuline fluorescence (Ex: Blue Epi light illumination; Em: Filter 530/28). Densitometric analysis was performed with Image Lab (Version 5.2.1, Bio-Rad).

**Table 3.** Polar lipid TLC Standards used for qHPTLC lipid class specific quantification.

| Polar Lipid TLC - ISTD | | | | | | | |
|---|---|---|---|---|---|---|---|
| Lipid | Calibration range [nmol] | | | | | | |
| TG 16:0/16:0/18:1 | 36.00 | 18.00 | 7.20 | 3.60 | 1.80 | 0.36 | 0.18 |
| PC 16:0/18:1 | 39.47 | 19.73 | 7.89 | 3.95 | 1.97 | 0.39 | 0.20 |
| PA 16:0/18:1 | 43.05 | 21.52 | 8.61 | 4.30 | 2.15 | 0.43 | 0.22 |
| FA 18:1 | 106.21 | 53.10 | 21.24 | 10.62 | 5.31 | 1.06 | 0.53 |
| LPC 18:1 | 57.51 | 28.75 | 11.50 | 5.75 | 2.88 | 0.58 | 0.29 |
| LPE 18:1 | 62.55 | 31.28 | 12.51 | 6.26 | 3.13 | 0.63 | 0.31 |
| PE 16:0/18:1 | 41.78 | 20.89 | 8.36 | 4.18 | 2.09 | 0.42 | 0.21 |
| PS 16:0/18:1 | 37.98 | 18.99 | 7.60 | 3.80 | 1.90 | 0.38 | 0.19 |
| SM 18:1;O2/18:1 | 41.15 | 20.57 | 8.23 | 4.11 | 2.06 | 0.41 | 0.21 |

##### 4.4 Liquid-Extraction Static Acquisition Mass Spectrometry

TLC plates were analyzed by using a TLC spot extraction system (Plate Express, Advion) coupled online to electrospray (ESI) Ion-Trap (IT) MS (amaZon SL, Bruker). TLC spots were extracted using pure MeOH as extraction solvent. ESI-IT parameter were as follows: electrospray voltage: 5.5 kV, end plate offset: 500 V; nebulizer gas: 7 psi; dry gas: 3 L/min; capillary temperature: 240 °C; sheath gas (He) flow rate: 25 arbitrary units. Spectra were acquired in enhanced resolution mode and recorded in either positive or negative polarity. A maximum ionization time of 50 ms was applied. Data were subsequently analyzed using *DataAnalysis* (Bruker Daltonics, Bremen, Germany)

#### 5 Liquid chromatography mass spectrometry (LC-MS) analysis

##### 5.1 Chromatography

##### 5.1.1 Unpolar lipid separation (C30 RPC)

Total lipid extracts (represented mostly by triacylglycerols) were reconstituted in $CHCl_3$/MeOH (2:1, v/v), required amount was transferred into HPLC vials and dried *in vacuo*. The dried lipids were reconstituted in i-PrOH/$CHCl_3$ (1:1, v/v) to a concentration of 2.5 $\mu g_{tissue}/\mu l_{i\text{-}PrOH}$ and 5 µl (= 12.5 $\mu g_{tissue}$) were loaded onto the column. Triacylglycerols were separated by reversed phase liquid chromatography (RPLC) on a Thermo Scientific Vanquish Horizon UHPLC system (Thermo Fisher Scientific, Germering, Germany) equipped with an Thermo Scientific Accucore C30 column (150 x 2.1 mm; 2.6 µm, 150 Å, Thermo Fisher Scientific, Sunnyvale, USA). Lipids were separated by gradient elution with solvent A (MeCN/$H_2O$, 1:1, v/v) and B (i-PrOH/MeCN/$H_2O$, 85:10:5, v/v) both containing 5 mM $NH_4HCO_2$ and 0.1% (v/v) formic acid. Separation was performed at 50°C with a flow rate of 0.3 mL/min using the following gradient: 0-5 min – 50 to 80 % B (curve 5), 5-22 min – 80 to 95 % B (curve 4), 22-26 min – 95 % isocratic, 26-27 min – 95 to 100 % B (curve 5), 27-47 min – 100 % B isocratic, 47-47.1 min – 100 to 50 % B followed by 8 min re-equilibration at 50% B.

##### 5.1.2 Polar lipid separation (C18 RPC)

Polar lipid fraction was reconstituted in $CHCl_3$/MeOH (2:1, v/v), required amount was transferred into HPLC vials and dried *in vacuo*. The dried lipids were reconstituted in pure i-PrOH to a concentration of 0.5 $mg_{tissue}/\mu l_{i\text{-}PrOH}$ and 5 µl (= 2.5 $mg_{tissue}$) were loaded onto the column. Polar lipid fractions (lyso-/phospholipids, sphingolipids, diacylglycerols) were separated by reversed phase liquid chromatography (RPLC) on a Vanquish Horizon UHPLC system (Thermo Fisher Scientific, Germering, Germany) equipped with an Thermo Scientific Accucore C18 column (150 x 2.1 mm; 2.6 µm, 150 Å, Thermo Fisher Scientific, Sunnyvale, USA). Lipids were separated by gradient elution with solvent A (MeCN/$H_2O$, 1:1, v/v) and B (i-PrOH/MeCN/$H_2O$, 85:10:5, v/v/v) both containing 5 mM $NH_4HCO_2$ and 0.1% (v/v) formic acid. Separation was performed at 50°C with a flow rate of 0.3 mL/min using following gradient: 0-20 min – 10 to 86 % B (curve 4), 20-22 min – 86 to 95 % B (curve 5), 22-26 min – 95 % isocratic, 26-26.1 min – 95 to 10 % B (curve 5) followed by 5 min re-equilibration at 10% B.

##### 5.1.3 Acylcarnitines separation (Si HILIC)

Polar lipid fraction was reconstituted in $CHCl_3$/MeOH (2:1, v/v), required amount was transferred into HPLC vials and dried *in vacuo*. The dried lipids were reconstituted in pure i-PrOH to a concentration of 0.5 $mg_{tissue}/\mu l_{i\text{-}PrOH}$ and 5 µl (= 2.5 $mg_{tissue}$) were loaded onto the column. Acylcarnitines were separated by hydrophilic interaction chromatography (HILIC) on a Vanquish Horizon UHPLC system (Thermo Fisher Scientific) equipped with an Acquity UPLC BEH HILIC Si column (100 x 1.0 mm; 1.7 µm, 130 Å, Waters Corp.). Lipids were separated as described previously(Lísa et al., 2017) by gradient elution with solvent A (MeCN/$H_2O$, 96:4, v/v) and B ($H_2O$) both containing 7 mM $NH_4OAc$. Separation was performed at 40°C with a flow rate of 0.15 ml/min using following gradient: 0-10 min – 0 to 10 % B (curve 5), 10-10.1 min – 10 to 0 % B (curve 5) followed by 5 min re-equilibration at 0% B.

##### 5.2 Mass Spectrometry

Each sample was analyzed on different LC-MS platforms (Figure 1A). High resolution accurate mass (HRAM) orbitrap based MS was employed in combination with C18 and C30 RPC, or HILIC chromatographic separation (Figure 1B). Overall, 111 LC-MS/MS analysis were performed for WAT lipids identification.

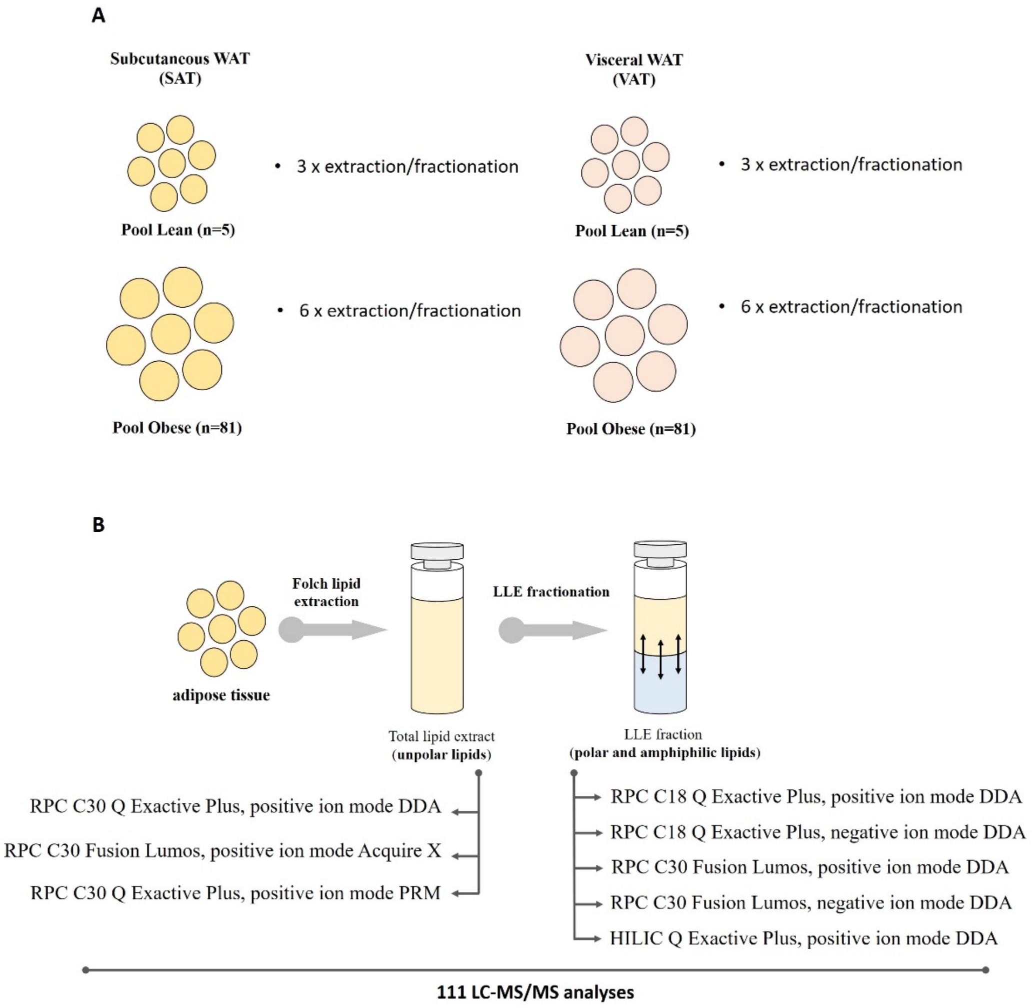


**Figure 1:** Scheme of LC-MS methodologies employed to increase lipid identification rates. **A** – schematic representation of pooled WAT samples used in the study with phenotype and depot specific number of lipid extraction replicates for each WAT pooled sample type. **B** – overview of LC and MS/MS acquisition strategies for each type of lipid extracts (total/unpolar lipids and LLE enriched polar and amphiphilic fraction).

##### 5.2.1 Data Dependent Acquisition on Q Exactive Plus Hybrid Quadrupole Orbitrap mass spectrometer

C18, C30 RPC and HILIC were coupled on-line to Thermo Scientific Q Exactive Plus Hybrid Quadrupole Orbitrap mass spectrometer (Thermo Fisher Scientific, Bremen, Germany) equipped with a HESI probe. Mass spectra were acquired in positive and negative modes with the following ESI parameters: sheath gas – 40 a.u., auxiliary gas – 10 L/min, sweep gas – 1 L/min, spray voltage – 3.5 kV (positive ion mode); -2.5 kV (negative ion mode), ion transfer temperature - 300 °C, S-lens RF level – 35% and aux gas heater temperature – 370 °C. For polar, unpolar lipids and acylcarnitines identification data were acquired in data dependent acquisition (DDA) modes with survey scan resolution of 140 000 (at *m/z* 200), AGC target 1e6 Maximum IT 100 ms in a scan range of *m/z* 350-1200 (380-1200 for unpolar lipids, 150-1200 for acyl carnitines). Data dependent MS2 were acquired with a resolution settings of 17 500 at 200 *m/z*, AGC target 1e5 counts, Maximum IT 60 ms, loop count 15, isolation window 1.2 *m/z* and stepped normalized collision energies of 10, 20 and 30 % (15, 20 and 30 % for unpolar lipids). A data dependent MS2 was triggered when an AGC target of 2e2 (2e3 for unpolar lipids, 2e1 for acyl carnitines) was reached followed by a Dynamic Exclusion for 10 s. All isotopes and charge states > 1 were excluded. All data was acquired in profile mode.

##### 5.2.2 Data Dependent Acquisition on Orbitrap Fusion Lumos Tribrid mass spectrometer

C30 RPC were coupled on-line to Thermo Scientific Orbitrap Fusion Lumos Tribrid mass spectrometer (Thermo Fisher Scientific, San Jose, USA) equipped with a HESI probe. Mass spectra were acquired in positive and negative modes with the following ESI parameters: sheath gas – 40 L/min, auxiliary gas – 10 L/min, spray voltage – +3.5 kV and -2.8 kV, capillary temperature – 250 °C, S-lens RF level – 25 and aux gas heater temperature – 320 °C. Data was acquired in data dependent acquisition mode with survey scan resolution of 120 000 (at *m/z* 200), AGC target 4e5, Maximum IT 100 ms in a scan range of *m/z* 500-950. Data dependent MS2 spectra were acquired with the following settings: automatic gain control target: 4e5 counts; max. injection time: 100 ms. The filters used were MIPS (small molecule), precursor selection range (500-950), charge state (1), dynamic exclusion (exclusion duration 8 seconds, exclude isotopes, mass tolerance ± 5 ppm) and target exclusion (polarity specific). MS/MS spectra were acquired in the Ion Trap mass analyzer at rapid scan rate (HCD, stepped collision energy: 10, 20, 30%; Isolation Window: 1.2 *m/z*; automatic gain control target: 2e4 counts; max. injection time: 35 ms, centroid data type).

##### 5.2.3 Acquire X on Orbitrap Fusion Lumos Tribrid mass spectrometer for in-depth triacylglycerol identification

Deep scan AcquireX Intelligent Data Acquisition Technology method was used to ensure in-depth identification of AT triacylglycerols. Two instrument methods were used – Full MS and ddMS$^4$. The exclusion override factor was set to 10 and $[M+H]^+$, $[M+NH_4]^+$, $[M+Na]^+$ were selected as preferred ions. Three solvents blank replicate were analyzed before sample injection and the latest was used as exclusion reference. Injection volume was always 1 µL. A Orbitrap Fusion Lumos Tribrid Mass Spectrometer (Thermo Fisher Scientific, San Jose, USA) using a HESI source was operated in positive ion mode using the following parameters: spray voltage – +3.5 kV, ion transfer tube temperature – 250°C, sheath gas – 30 arbitrary units, aux gas – 10, sweep gas – 1, arbitrary units, vaporizer temperature – 300°C. Full MS were performed by the Orbitrap mass analyzer, operated at a resolution setting of 120,000 for *m/z* 200, scan range of *m/z* 500–1200, AGC target 4e5 counts, maximum IT 100 ms, RF level: 25%, data type profile, EASY-IC internal calibration. For MS/MS resolution setting of 15,000 for *m/z* 200, HCD (collision energy: 20, 35, 50%), isolation window of 1.2 *m/z*, AGC target 2e4, Maximum IT 40 ms, data type centroid were used. The filters used were MIPS small molecule, precursor selection range (*m/z* 500-1200), charge state 1, dynamic exclusion for 6 s, exclude isotopes, mass tolerance ± 10 ppm, and Xcalibur Acquire X generated exclusion and inclusion lists (mass tolerance 25 ppm). The MS3 and relative MS4 spectra were acquired only for ions which fulfilled the two following filters: acquisition neutral loss ion trigger and loss trigger (list of FA, tolerance ±20 ppm). MS3 spectra were acquired in the orbitrap mass spectrometer at resolution settings of 15000 for *m/z* 200 (HCD, Collision Energy: 35%; Isolation Window: 1; automatic gain control target: 2e4 counts; max. injection time: 25 ms, centroid). MS4 analysis were performed in the ion trap at normal scan rate (CID, Collision Energy: 30%; Activation time: 10 ms; Activation Q: 0.25, Isolation Window: 1; automatic gain control target: 2e4 counts; max. injection time: 35 ms, centroid).

##### 5.2.4 Targeted identification of cholesteryl esters

The total lipid extract (unpolar fraction) was analyzed via C30 RPC coupled on-line to a Q Exactive Plus Hybrid Quadrupole Orbitrap mass spectrometer (Thermo Fisher Scientific, Bremen, Germany) equipped with a HESI probe. Mass spectra were acquired in positive mode with the following ESI parameters: sheath gas – 40 au, auxiliary gas – 10 au, sweep gas – 1 au, spray voltage – 3.5 kV, ion transfer temperature – 300 °C, S-lens RF level – 35% and aux gas heater temperature – 370 °C. For cholesteryl ester identification data were acquired in parallel reaction monitoring (PRM) mode. An inclusion list of the ammoniated adducts of 36 cholesteryl esters covering the range of fatty acids from 2 up to 22 carbons and 0 to 6 double bonds was employed. PRM data was acquired with a resolution setting of 17 500 at 200 *m/z*, AGC target 2e5 counts, Maximum IT 200 ms, isolation window 1.2 *m/z* and a normalized collision energy of 20 % in profile mode.

##### 5.2.5 Co-elution of short chain TGs, MGs, and acylcarnitine's with corresponding internal standards

The retention behavior of short chain TGs, MGs and acylcarnitines was additionally studied in C18 RPC for verification of their identification. The short chain TG mixture TAG-MIX-10 (Larodan, Solna, Sweden) was dissolved in $CHCl_3$/MeOH (2:1, v/v) and an amount corresponding to 2.5 µg total TG was transferred to an HPLC vial and dried *in vacuo*. Dried TAGs were redissolved in pure *i*-PrOH (100 µL) and 5 µL were injected. TAGs were chromatographically separated by C18 RP and analyzed by MS in DDA mode as described above (Supplemental Figure 2).

The MG mixture MAG-MIX-1 (Larodan, Solna, Sweden) was dissolved in $CHCl_3$/MeOH (2:1, v/v) and an amount corresponding to 1.5 µg total MG was transferred to an HPLC vial and dried *in vacuo*. Dried MGs were redissolved in pure *i*-PrOH (100 µL) and 5 µL were injected. MGs were chromatographically separated by C18 RP and analyzed by MS in DDA mode as described above (Supplemental Figure 2).

The non-deuterated acylcarnitine mixture NSK-B-US-1 (Cambridge Isotope Laboratories, Inc., Tewksbury, Massachusetts, USA) was dissolved in 1 mL $CHCl_3$/MeOH (2:1, v/v) and 75 µL were transferred to an HPLC vial and dried *in vacuo*. Dried acylcarnitines were redissolved in pure *i*-PrOH (75 µL) and 5 µL were injected. Acylcarnitines were chromatographically separated by C18 RP or HILIC and analyzed by MS in DDA mode as described above (Supplemental Figure 2).

#### 6 Lipid Identification

DDA and Acquire X LC-MS/MS datasets were used for lipid molecular species identification. Identification strategy relied on 3 software followed by manual annotation.

##### 6.1 LipidSearch

Lipids were identified using Thermo Scientific LipidSearch software version 4.1 SP1 using the following key processing parameters: target database – general, precursor tolerance ± 5 ppm, product tolerance ± 20 ppm, product ion threshold 5%, m-score threshold 1, Quan *m/z* tolerance ± 5 ppm, Quan RT (retention time) range ± 0.5 min. According to the lipid class, the ion mode and the structure of the lipids the results where filtered according to the "Grades". (Lyso)phospholipids were identified with Grade A (diacyl forms) and Grade B (ether forms). Sphingomyelins were accepted at Grade C. Ceramides were used after filtering with Grade A and Grade C for corresponding water loss ions, and TG/DG lipids were identified with Grade A only.

##### 6.2 LipidHunter (Ni et al., 2017)

LipidHunter 2 RC source code version was used (https://github.com/SysMedOs/lipidhunter) to identify phospholipids in negative mode ($[M+HCOO]^-$ for PC and $[M-H]^-$ all others) and glycerolipids ($[M+NH_4]^+$) in positive mode. Raw files were converted into mzML format using MSConvert from Proteowizard project (version 3.0.9134). Lipids were identified using the following parameters: mass accuracy at MS level - 5 ppm, MS intensity threshold - 3e3 counts, mass accuracy at MS/MS level - 20 ppm, MS/MS intensity threshold - 100 counts. Identifications were filtered for isotopic score ≥ 75, and rank score ≥ 30. The white list of considered fatty acyl chains included 123 fatty acids corresponding to the LipidSearch default configuration file "FattyAcidDefinition.xml" for inter-software compatibility. Weight Factors assigned to each FA neutral loss fragment ions were 40, 48 and 30% for PLs, DGs and TGs, respectively. The identification results were reviewed using interactive HTML report. The table output files from LipidHunter were filtered and merged. Additional filters applied for PLs identification included: isotope score ≥80, rank score ≥40, both FA residues identified (for O/P-containing PLs, only one FA residue should be present, rank score ≥30). For DGs and TGs, all FA residues have to be identified, the isotope score filter was set to ≥ 85.

##### 6.3 Lipostar (Goracci et al., 2017)

Lipostar (version 1.0.6, Molecular Discovery, Hertfordshire, UK) equipped with LIPID MAPS structure database (version December 2017) was used. The raw files were imported directly, and aligned using default settings. Automatic peak picking was performed with SDA smoothing level set to low and minimum S/N ratio 3. Automatic isotope clustering settings were set to 7 ppm with RT tolerance 0.2 min. An "MS2 only" filter was applied to keep only features with MS/MS spectra for identification. Following parameters were used for lipid identification: 5 ppm precursor ion mass tolerance and 10 ppm product ion mass tolerance. The automatic approval was performed to keep structures with quality of 3-4 stars. Identification results of each lipid class were exported separately into 3 files using the Lipostar export function: feature table, best 3 matches of each feature (with check chain fragments enabled), and all approved matches. Exported tables were used for additional filtering and generation of merged identification list. For all classes, a fragmentation score filter of 60 and at least 2 fragment matches were applied (for TG at least 3 fragment matches).

##### 6.4 Merge of identification results

Results of software assisted lipid identification were merged, and only lipids fulfilling the following parameter were kept in the final list: lipid must be identified by at least two software, within $\Delta RT < 0.3$ min. A chain length filter according to the FA included 58 fatty acids (from C4 to C26; maximum of six double bonds) were applied to all software results. A set of customized python scripts was used to filter and merge the output files. The corresponding source code is available on GitHub (https://github.com/SysMedOs/AdipoAtlasScripts). Ether PLs identified by the software were manually corrected based of the retention time mapping.

##### 6.5 Manual lipid annotation

Manual annotation and retention time mapping were performed as described in Supplementary File 1.

#### 7 Quantification

##### 7.1 LC-MS quantification of acylcarnitines, polar and unpolar lipids

For quantification purposes, the respective lipid classes were separated on a RPC C30, C18 or HILIC as described above. MS data were acquired in Full MS mode on a Q Exactive Plus Hybrid Quadrupol Orbitrap mass spectrometer in the positive and negative ion mode at the resolution of 140,000 at *m/z* 200, AGC target of 1e6 and a Maximum IT of 100 ms in the mass range from *m/z* 100 – 1500. Data were acquired in profile mode.

##### 7.2 Generation of calibration curves of employed ISTD

In order to ensure linear response of the employed standards, internal calibration curves were generated for the respective ISTD. Varying concentrations of ISTD were spiked into ≈50 mg pooled WAT prior to lipid extraction to generate a 5-point internal calibration curve (Table 4). Lipids were extracted and ISTD derived signals were quantified. Additionally, isobaric or isomeric overlap with endogenous compounds during LC-MS analysis was excluded by close inspection of LC-MS derived data. Only calibration points resulted in a calibration curve with $R > 0.98$ were approved. Final ISTD concentration to spike for subsequent quantification had to display a relative standard deviation of < 20% and represent the linear response range. Final spiked concentrations are marked in green in Table 4.

**Table 4.** Spiked concentrations to generate an internal calibration curve. Marked in green are the final used ISTD concentrations.

| Lipid | Spiked in ≈50 mg WAT [nmol] | | | | | |
|---|---|---|---|---|---|---|
| SPLASH LIPIDOMIX | | | | | | |
| PC 15:0_18:1 (d7) | 0 | 0.107 | 0.213 | 1.067 | 2.134 | 4.268 |
| PE 15:0_18:1 (d7) | 0 | 0.004 | 0.008 | 0.040 | 0.080 | 0.160 |
| PS 15:0_18:1 (d7) | 0 | 0.003 | 0.005 | 0.027 | 0.054 | 0.108 |
| PG 15:0_18:1 (d7) | 0 | 0.019 | 0.038 | 0.190 | 0.381 | 0.762 |
| PI 15:0_18:1 (d7) | 0 | 0.005 | 0.011 | 0.054 | 0.107 | 0.215 |
| PA 15:0_18:1 (d7) | 0 | 0.005 | 0.011 | 0.054 | 0.107 | 0.215 |
| LPC 18:1 (d7) | 0 | 0.024 | 0.048 | 0.241 | 0.482 | 0.965 |
| LPE 18:1 (d7) | 0 | 0.005 | 0.011 | 0.054 | 0.109 | 0.218 |
| CE 18:1 (d7) | 0 | 0.271 | 0.541 | 2.705 | 5.411 | 10.821 |
| MG 18:1 (d7) | 0 | 0.003 | 0.006 | 0.028 | 0.055 | 0.110 |
| DG 15:0_18:1 (d7) | 0 | 0.008 | 0.016 | 0.080 | 0.160 | 0.320 |
| TG 15:0_18:1_15:0 (d7) | 0 | 0.035 | 0.071 | 0.353 | 0.705 | 1.411 |
| SM d18:1_18:1 (d9) | 0 | 0.021 | 0.042 | 0.209 | 0.419 | 0.837 |
| Chol (d7) | 0 | 0.125 | 0.250 | 1.250 | 2.499 | 4.999 |

| | | | | | | |
|---|---|---|---|---|---|---|
| **Cer/SpH Mix I (Avanti)** | | | | | | |
| **SPB 17:1;O2** | 0 | 0.019 | 0.038 | 0.188 | 0.375 | 0.750 |
| **SPB 17:0;O2** | 0 | 0.019 | 0.038 | 0.188 | 0.375 | 0.750 |
| **SPBP 17:1;O2** | 0 | 0.019 | 0.038 | 0.188 | 0.375 | 0.750 |
| **SPBP 17:0;O2** | 0 | 0.019 | 0.038 | 0.188 | 0.375 | 0.750 |
| **Lac-Cer 18:1;O2/12:0** | 0 | 0.019 | 0.038 | 0.188 | 0.375 | 0.750 |
| **Gluc-Cer d18:1;O2/12:0** | 0 | 0.019 | 0.038 | 0.188 | 0.375 | 0.750 |
| **SM 18:1;O2/12:0** | 0 | 0.019 | 0.038 | 0.188 | 0.375 | 0.750 |
| **Cer 18:1;O2/12:0** | 0 | 0.019 | 0.038 | 0.188 | 0.375 | 0.750 |
| **CerP 18:1;O2/12:0** | 0 | 0.019 | 0.038 | 0.188 | 0.375 | 0.750 |
| **Cer 18:1;O2/25:0** | 0 | 0.019 | 0.038 | 0.188 | 0.375 | 0.750 |
| **Acylcarnitine NSK-1-B Mix** | | | | | | |
| **Free Carnitine (d9)** | 0 | 0.008 | 0.016 | 0.081 | 0.162 | 0.324 |
| **CAR 2:0 (d3)** | 0 | 0.002 | 0.004 | 0.018 | 0.037 | 0.074 |
| **CAR 3:0 (d3)** | 0 | 0.000 | 0.001 | 0.004 | 0.007 | 0.015 |
| **CAR 4:0 (d3)** | 0 | 0.000 | 0.001 | 0.004 | 0.007 | 0.015 |
| **CAR 5:0 (d9)** | 0 | 0.000 | 0.001 | 0.004 | 0.007 | 0.015 |
| **CAR 8:0 (d3)** | 0 | 0.000 | 0.001 | 0.004 | 0.007 | 0.014 |
| **CAR 14:0 (d9)** | 0 | 0.000 | 0.001 | 0.004 | 0.007 | 0.015 |
| **CAR 16:0 (d3)** | 0 | 0.001 | 0.002 | 0.008 | 0.015 | 0.030 |
| **Additionally added** | | | | | | |
| **FA 18:0 ([13]C1)** | 0 | 0.500 | 1.000 | 5.000 | 10.000 | 20.000 |
| **TG 18:1/18:1/18:1 ([13]C3)** | 0 | 50.000 | 100.000 | 500.000 | 1000.000 | 2000.000 |
| **TG 16:0/16:0/16:0 ([13]C3)** | 0 | 2.500 | 5.000 | 25.000 | 50.000 | 100.000 |
| **DG 18:1/18:1/0:0 ([13]C3)** | 0 | 0.500 | 1.000 | 5.000 | 10.000 | 20.000 |
| **Cer 18:0;O2/12:0** | 0 | 0.019 | 0.038 | 0.188 | 0.375 | 0.750 |
| **Cer 18:0;O2/8:0** | 0 | 0.019 | 0.038 | 0.188 | 0.375 | 0.750 |
| **Cer 18:1;O/6:0** | 0 | 0.019 | 0.038 | 0.188 | 0.375 | 0.750 |
| **Cer 18:0;O3/16:0** | 0 | 0.019 | 0.038 | 0.188 | 0.375 | 0.750 |
| **Cer 18:0;O3/8:0** | 0 | 0.019 | 0.038 | 0.188 | 0.375 | 0.750 |
| **Cer 18:1;O2/17:0,O[2*R*-OH]** | 0 | 0.019 | 0.038 | 0.188 | 0.375 | 0.750 |

##### 7.3 Quantification and Data Processing

For quantification raw data sets of Full MS measurements were processed using Thermo Scientific TraceFinder 4.1 (Thermo Fisher Scientific, Bremen, Germany). Quantification was based on determination of area under curve (AUC) using following settings: mass tolerance – 5 ppm, area noise factor – 5, peak noise factor – 10, baseline window – 150, S/N ≥ 3 using ICIS detection algorithm. Signals to be quantified (adducts and in-source fragments) were determined based on measurement of representative lipid standards for each class as shown below (Table 5).

For quantification of cholesteryl esters, raw data sets were acquired in PRM mode as described above. Measurements were processed using TraceFinder$^{TM}$ 4.1 (Thermo Fisher Scientific, Bremen, Germany). Quantification was based on determination of area under curve (AUC) using following settings: mass tolerance – 20 ppm, area noise factor – 5, peak noise factor – 10, baseline window – 150, S/N ≥ 3 using ICIS detection algorithm.

**Table 5.** Adducts/in-source fragments and corresponding internal standards used for the quantification of each lipid class.

| Lipid Class | Adducts/Fragments | Used ISTD |
|---|---|---|
| **TG** | [M+NH4]+, [M+Na]+, [M+K]+ | TG 18:1/18:1/18:1 ([13]C3) |
| **DG** | [M+H]+, [M+Na]+ | DG 18:1/18:1 ([13]C3) |
| **PC** | [M+H]+ | PC 15:0_18:1 (d7) |
| **PE** | [M+H]+ | PE 15:0_18:1 (d7) |
| **LPC** | [M+H]+ | LPC 18:1 (d7) |
| **LPE** | [M+H]+ | LPE 18:1 (d7) |
| **PG** | [M+H]+, [M+NH4]+ | PG 15:0_18:1 (d7) |
| **PS** | [M-H]- | PS 15:0_18:1 (d7) |
| **PI** | [M-H]- | PI 15:0_18:1 (d7) |
| **Cer** | [M+H]+, [M+Na]+, [M-H2O+H]+ | Cer 18:1;O2/12:0 ; Cer 18:1;O2/25:0 |
| **Dihydro-Cer** | [M+H]+, [M+Na]+, [M-H2O+H]+ | Cer 18:0;O2/12:0 ; Cer 18:0;O2/8:0 |
| **Deoxy-Cer** | [M+H]+, [M+Na]+ | Cer 18:1;O/6:0 |
| **Phyto-Cer** | [M+H]+, [M+Na]+ | Cer 18:0;O3/16:0 ; Cer 18:0;O3/8:0 |
| **Cer + a-OH FA** | [M+H]+, [M+Na]+, [M-H2O+H]+ | Cer 18:1;O2/17:0;O |
| **Hex-Cer** | [M+H]+, [M+Na]+, [M-H2O+H]+ | Gluc-Cer 18:1;O2/12:0 |
| **Hex-2-Cer** | [M+H]+, [M+Na]+, [M-H2O+H]+ | Lac-Cer 18:1;O2/12:0 |
| **SM** | [M+H] | SM 18:1;O2/18:1 (d7) |
| **CAR** | [M+H]+ | CAR 2:0 - CAR 16:0 (d3-d9) |
| **Carnitine** | [M+H]+ | Free Carnitine (d9) |
| **Cholesteryl Esters** | PRM: [M+NH4]+ → [Cholesterol-$H_2O$+H]+ | CE 18:1 (d7) |

Obtained AUC values for all adducts and in-source fragments of each lipid specie were summed up in order to display all-ion abundance of studied lipid species. AUC values were corrected for $^{13}C$ abundance (Type I correction following the guidelines of Lipidomics Standards Initiative(Burla et al., 2018) as described before(Lange and Fedorova, 2020):

$$AUC_{n(k)\ total} = AUC_{n(k)}\left( 1 + 0.0109n + \frac{0.0109^2 n(n-1)}{2}\right)$$

$AUC_{n(k)\ total}$ = total ion area under curve, $AUC_{n(k)}$ = quantified area under curve of monoisotopic mass, n = No. of C-Atoms, k = No. of double bonds

Due to incomplete labeling of ISTDs, AUC of deuterated ISTD for phospholipids, sphingomyelins and acyl carnitines were determined by summing up the AUC of $[M-2]^+$ ($d_{x-2}$), $[M-1]^+$ ($d_{x-1}$),$[M]^+$ ($d_x$), $[M+1]^+$ ($d_x$; $^{13}C_1$) and $[M+2]^+$ ($d_x$; $^{13}C_2$).

AUC of non-labeled ISTD for different ceramide classes were corrected for $^{13}C$ abundance (Type I correction following the guidelines of Lipidomics Standards Initiative(Burla et al., 2018) as described before(Lange and Fedorova, 2020):

$$AUC_{n(k)\ total} = AUC_{n(k)}\left( 1 + 0.0109n + \frac{0.0109^2 n(n-1)}{2}\right)$$

Quantitative values for lipid species were determined by relating AUC of the used ISTD to the lipid specie AUC:

$$C_{Lipid} = \frac{AUC_{Lipid}}{AUC_{ISTD}} * C_{ISTD}$$

$C_{Lipid}$ = Concentration of lipid specie, $C_{ISTD}$ = concentration of ISTD, $AUC_{Lipid}$ = corrected area under curve for lipid specie, $AUC_{ISTD}$ = area under curve for ISTD.

##### 7.4 Response factors calculations for triacylglycerols

TGs were quantified using a single $^{13}C_3$ labeled TG molecular species (TG 18:1/18:1/18:1 ([13]C3)). Type II isotopic correction was applied for determination of AUC of $^{13}C_3$ labeled TG, due to coelution with the native TG and isobaric overlap of $[M]^+$ of $^{13}C_3$ labeled TAG and $[M+3]^+$ of native TG.

**Table 6.** TG species employed for the generation of response factors. TGs were mixed and sequentially diluted in order to generate calibration curves for each single TG species.

| Bulk | Molecular species | on column [pmol] | | | | |
|---|---|---|---|---|---|---|
| **TG 24:0** | TG 8:0/8:0/8:0 | 132.19 | 26.44 | 5.29 | 1.06 | 0.21 |
| **TG 27:0** | TG 9:0/9:0/9:0 | 121.35 | 24.27 | 4.85 | 0.97 | 0.19 |
| **TG 30:0** | TG 10:0/10:0/10:0 | 112.14 | 22.43 | 4.49 | 0.90 | 0.18 |
| **TG 33:0** | TG 11:0/11:0/11:0 | 104.24 | 20.85 | 4.17 | 0.83 | 0.17 |
| **TG 36:0** | TG 12:0/12:0/12:0 | 97.37 | 19.47 | 3.89 | 0.78 | 0.16 |
| **TG 39:0** | TG 13:0/13:0/13:0 | 91.36 | 18.27 | 3.65 | 0.73 | 0.15 |
| **TG 42:0** | TG 14:0/14:0/14:0 | 86.04 | 17.21 | 3.44 | 0.69 | 0.14 |
| **TG 42:3** | TG 14:1/14:1/14:1 | 92.97 | 18.59 | 3.72 | 0.74 | 0.15 |
| **TG 45:0** | TG 15:0/15:0/15:0 | 81.31 | 16.26 | 3.25 | 0.65 | 0.13 |
| **TG 48:0** | TG 16:0/16:0/16:0 | 93.59 | 18.72 | 3.74 | 0.75 | 0.15 |
| **TG 48:3** | TG 16:1/16:1/16:1 | 83.20 | 16.64 | 3.33 | 0.67 | 0.13 |
| **TG 51:0** | TG 17:0/17:0/17:0 | 73.25 | 14.65 | 2.93 | 0.59 | 0.12 |
| **TG 54:0** | TG 18:0/18:0/18:0 | 77.27 | 15.45 | 3.09 | 0.62 | 0.12 |
| **TG 54:3** | TG 18:1/18:1/18:1 | 229.64 | 45.93 | 9.19 | 1.84 | 0.37 |
| **TG 54:6** | TG 18:2/18:2/18:2 | 79.60 | 15.92 | 3.18 | 0.64 | 0.13 |
| **TG 54:9** | TG 18:3/18:3/18:3 | 34.35 | 6.87 | 1.37 | 0.27 | 0.05 |
| **TG 57:0** | TG 19:0/19:0/19:0 | 66.65 | 13.33 | 2.67 | 0.53 | 0.11 |
| **TG 60:0** | TG 20:0/20:0/20:0 | 67.19 | 13.44 | 2.69 | 0.54 | 0.11 |
| **TG 60:3** | TG 20:1/20:1/20:1 | 72.19 | 14.44 | 2.89 | 0.58 | 0.12 |
| **TG 60:6** | TG 20:2/20:2/20:2 | 69.19 | 13.84 | 2.77 | 0.55 | 0.11 |
| **TG 60:9** | TG 20:3/20:3/20:3 | 69.61 | 13.92 | 2.78 | 0.56 | 0.11 |
| **TG 60:12** | TG 20:4/20:4/20:4 | 70.07 | 14.01 | 2.80 | 0.56 | 0.11 |
| **TG 60:15** | TG 20:5/20:5/20:5 | 70.50 | 14.10 | 2.82 | 0.56 | 0.11 |
| **TG 63:0** | TG 21:0/21:0/21:0 | 61.14 | 12.23 | 2.45 | 0.49 | 0.10 |
| **TG 54:3 (13[C]3)** | TG 18:1/18:1/18:1 (13[C]3) | 85.00 | 17.00 | 3.40 | 0.68 | 0.14 |
| **TG 48:0 (13[C]3)** | TG 16:0/16:0/16:0 (13[C]3) | 85.00 | 17.00 | 3.40 | 0.68 | 0.14 |
| **TG 34:1 (d5)** | TG 14:0/16:1/14:0 (d5) | 74.81 | 14.96 | 2.99 | 0.60 | 0.12 |
| **TG 38:1 (d5)** | TG 15:0/18:1/15:0 (d5) | 74.44 | 14.89 | 2.98 | 0.60 | 0.12 |
| **TG 50:0 (d5)** | TG 16:0/18:0/16:0 (d5) | 75.94 | 15.19 | 3.04 | 0.61 | 0.12 |
| **TG 51:1 (d5)** | TG 17:0/17:1/17:0 (d5) | 77.63 | 15.53 | 3.11 | 0.62 | 0.12 |

| | | | | | | |
|---|---|---|---|---|---|---|
| **TG 50:0 (d5)** | TG 19:0/12:0/19:0 (d5) | 75.19 | 15.04 | 3.01 | 0.60 | 0.12 |
| **TG 60:1 (d5)** | TG 20:0/20:1/20:0 (d5) | 71.44 | 14.29 | 2.86 | 0.57 | 0.11 |
| **TG 58:7 (d5)** | TG 20:2/18:3/20:2 (d5) | 74.25 | 14.85 | 2.97 | 0.59 | 0.12 |
| **TG 58:10 (d5)** | TG 20:4/18:2/20:4 (d5) | 73.13 | 14.63 | 2.93 | 0.59 | 0.12 |
| **TG 62:11 (d5)** | TG 20:5/22:6/20:5 (d5) | 75.56 | 15.11 | 3.02 | 0.60 | 0.12 |

TG molecular species experience differential ionization and ion transmission yielding varying intensities depending on the molecular structure. Response factors of different native and isotopically labeled standards (Table S6) relative to the used $^{13}C_3$ labeled ISTD TG 18:1/18:1/18:1 ([13]C3) were determined in order to increase accuracy of TG quantification. Dilution series of 34 different labeled and unlabeled standards spanning over three orders of magnitude were recorded (Table 6). Calibration curves were generated for each species to yield linear regression models with a minimum $R^2 > 0.99$ and RSD < 10 % for each point of the dilution curve. Obtained slopes of concentration-response curves were related to the concentration-response curve of the isotopically labeled ISTD to establish response factors (RF). RF were normalized to the number of carbons and double bonds (DB) by dividing RF with the equivalent-carbon-number (ECN) yielding the normalized response factor ($RF_{norm}$). Deviations in response mainly arise from increasing unsaturation and therefore for each DB number $RF_{norm}$ was separately related to *m/z* of $[M+NH_4]^+$. Resulting slopes and y-intercepts for each double bond number were used to establish a linear regression model for determination of slopes and y-intercepts for which no standard were available. Subsequently, $RF_{norm}$ is calculated for each identified TG species and used for the quantification. A detailed description of the workflow provided in Supplementary Figure S4. Finally, accurate quantification of TG was performed by relating the intensity of the ISTD and the RF-corrected intensity of the TG molecular species as expressed by the following equation:

$$AUC_{TG,corrected} = \frac{AUC_{TG}}{RF_{TG}}$$

$$C_{TG} = \frac{AUC_{TG,corrected}}{AUC_{ISTD}} * C_{ISTD}$$

**8 Software**

Tracefinder 4.1 (Thermo Fisher Scientific, Bremen, Germany) was used for targeted lipid quantification. Quantitative data analysis including isotopic correction, ISTD normalization and response factor normalization was performed with Microsoft Excel 2016. Graphical representations were generated with Graphpad Prism® 5.02 and OriginPro© 2017. Metaboanalyst (www.metaboanalyst.ca/) was used to generate heatmaps and perform statistical analysis. A lipid was found to be statistically significantly regulated by Students t-test with a threshold of $p \leq 0.05$ (FDR adjusted) assuming equal variances and a fold change $\geq 2$, or with an ANOVA $p \leq 0.05$.

**9 Data availability**

All raw LC-MS/MS and LC-MS files used for WAT lipidome identification and quantification, respectively, are available at massive.ucsd.edu (MSV000086729). File names correspond to the following LC-MS/MS datasets (Table S7).

Xiaoning, Lu; Michael, Y. Enrichment of Phospholipids in Biological Samples Using HybridSPE-PL". Report. US *28.3*.
