## Supplemental Figures for "AdipoAtlas: A Reference Lipidome for Human White Adipose Tissue"

### Supplementary information

**Table S1:** Lipid molecular species identified in human white adipose tissue lipid extracts by various software tools and/or manual annotation. PRM = parallel reaction monitoring; HILIC = hydrophilic interaction chromatography, RPC = reversed phase chromatography, coupled on-line to Q Exactive Plus MS; C30* RPC - C30 RPC coupled on-line to Fusion Lumos MS.

**Table S2:** Mixture of internal standards (ISTD) spiked in white adipose tissue samples prior to lipid extraction.

**Table S3:** Lipid class specific adducts and in-source fragments used for quantification of molecular lipid species. Internal standards (ISTD) used for quantification of naturally occurring lipids in lipid extracts of human WAT.

**Table S4:** Quantification of lipid molecular species from human white adipose tissue samples (fmol/µg protein). Marked in gray/italic are lipids that were detected in <75% of the samples.

**Table S5:** Signal intensity of spiked internal standards (ISTD) within the measured sample cohort. STDEV = standard deviation, RSD = relative standard deviation.

**Table S6:** Response factors (RF) for TG molecular species to increase quantitative accuracy. Determination of TG RF follows workflow as presented in FigS5.

**Table S7:** Data availability. List of filenames corresponding to the analyses employed for identification and quantification of lipids from human white adipose tissue. DDA = data dependent acquisition, PRM = parallel reaction monitoring, RPC = reversed phase chromatography, HILIC = hydrophilic interaction chromatography.

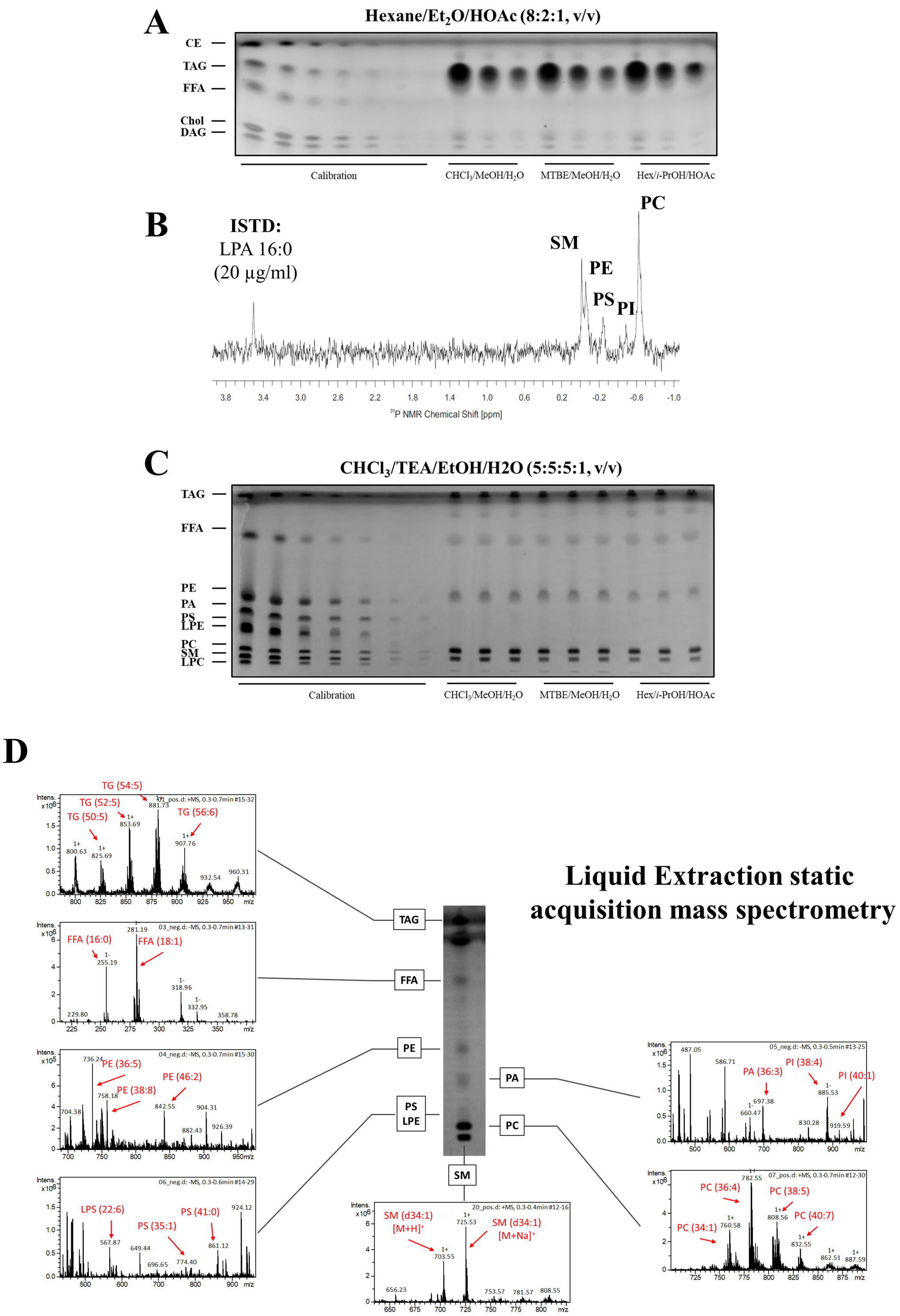


**Figure S1:** Quantification of lipid extraction efficiency of unpolar and polar lipid classes. **A:** Representative image of unpolar lipids extracted using Folch, MTBE, and Hex/IPA methods quantified by high performance thin layer chromatography (qHPTLC). **B:** Representative spectrum of $^{31}$P-NMR based quantification of phosphate containing lipid classes from a white adipose tissue (WAT) lipid extract. **C:** Representative image of qHPTLC based polar lipid quantification from the enriched polar fraction of WAT lipid extracted by Folch, MTBE, and Hex/IPA methods. **D:** Verification of identity lipid classes separated by qHPTLC using liquid extraction-static acquisition low resolution mass spectrometry.

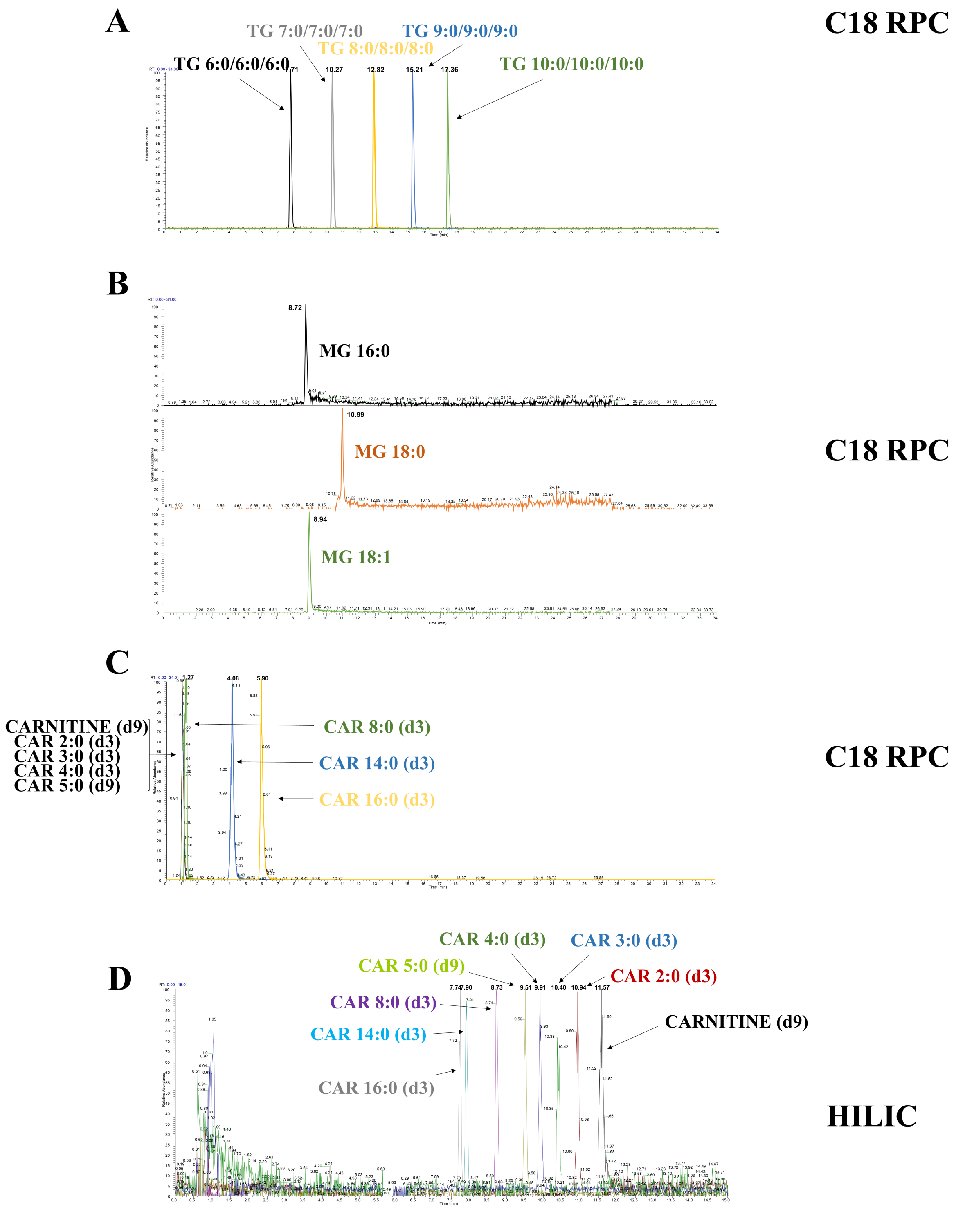


**Figure S2:** Investigation of retention behavior of lipids on various liquid chromatography (LC) platforms coupled to Q-Orbitrap high resolution mass spectrometry (QExactive). **A:** Separation of a mixture of short-chain triacylglycerols (TG) on C18 reversed phase chromatography (RPC). **B:** Separation of a mixture of monoacylglycerols (MG) on C18 RPC. Separation of a mixture of deuterated acylcarnitines (CAR) on **C:** C18 RPC and **D:** hydrophilic interaction chromatography (HILIC).

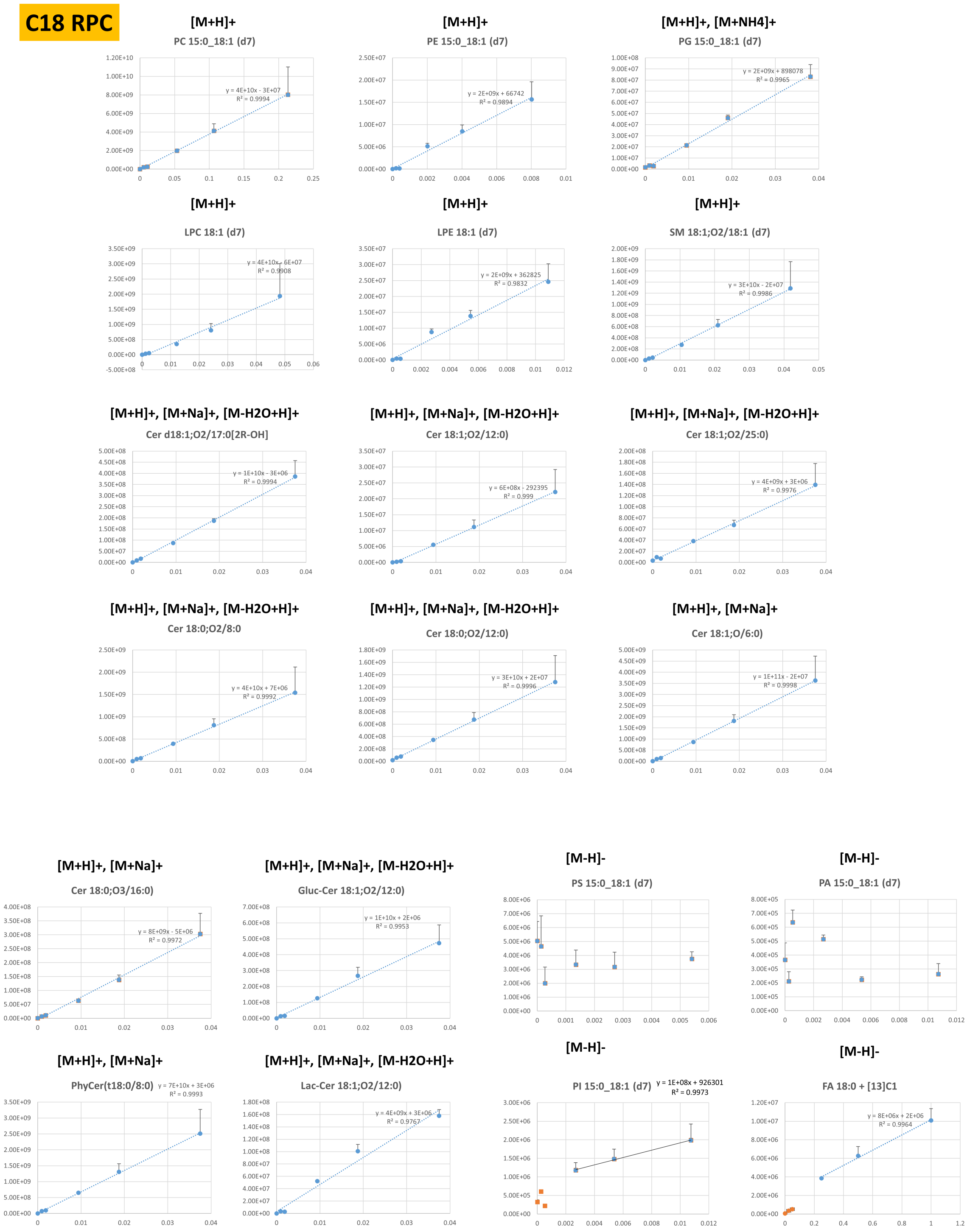


**Figure S3:** Internal calibration curves of spiked internal standards (ISTD) into white adipose tissue (WAT) samples prior to lipid extraction and fractionation. Phospholipid and Sphingolipid ISTD were separated on C18 reversed phase chromatography (C18 RPC). Only calibration points resulting in a linear regression $R < 0.98$ (blue) were used for calibration curve generation. Other calibration points were excluded (orange).

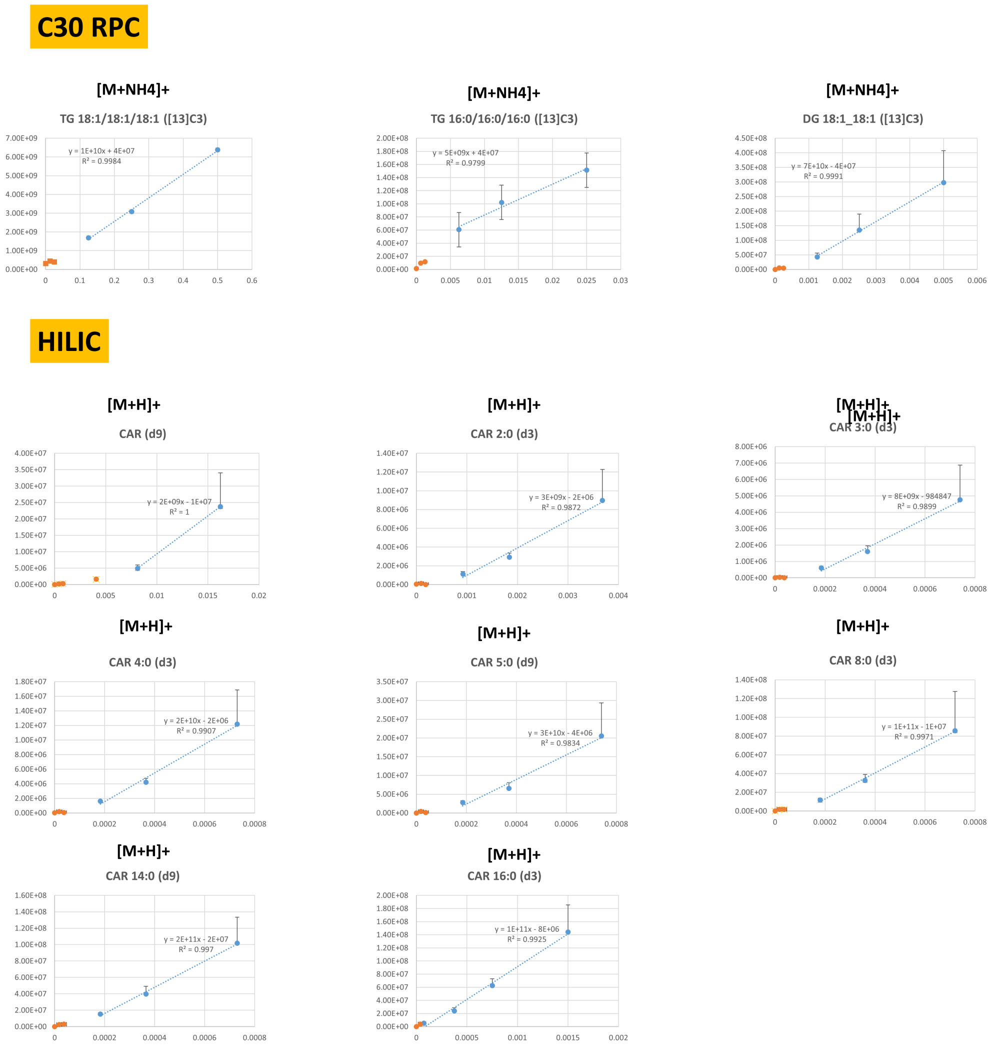


**Figure S4:** Internal calibration curves of spiked internal standards (ISTD) into white adipose tissue (WAT) samples prior to lipid extraction and fractionation. Glycerolipid and acylcarnitine ISTDs were separated on C30 reversed phase chromatography (C18 RPC) or hydrophilic interaction chromatography (HILIC), respectively. Only calibration points resulting in a linear regression R < 0.98 (blue) were used for calibration curve generation. Other calibration points were excluded (orange).

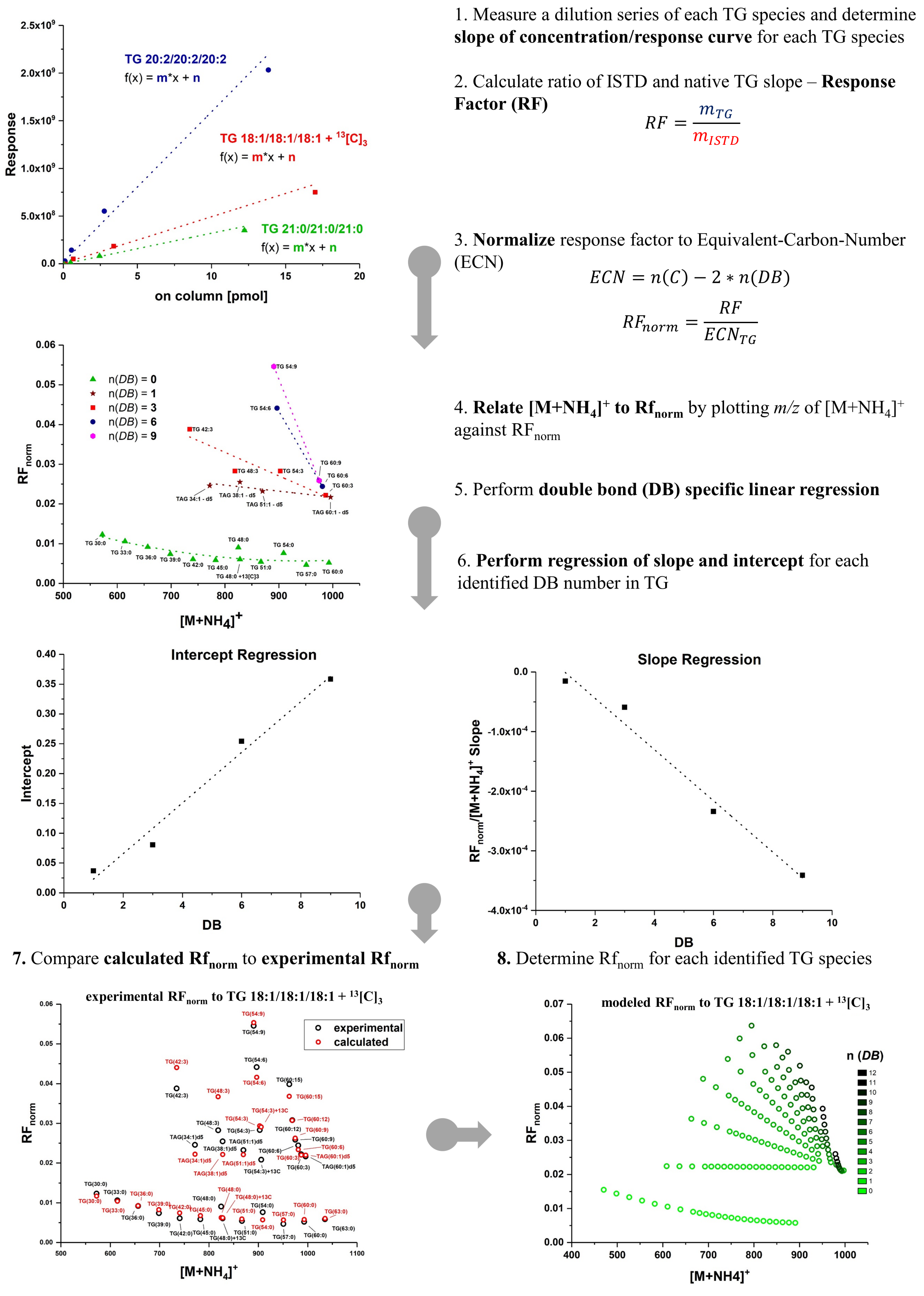


**Figure S5:** Generation of response factors (RF) for triacylglycerols (TG) on C30 reversed phase chromatography. The slope of external calibration curves for 33 native and deuterated TG molecular species with varying degrees of fatty acid unsaturation and length was determined. Each slope was related to the slope of $^{13}C$ labeled TG used as ISTD during lipid extraction, i.e. a response factor (RF) was generated. The dependence of fatty acid chain length and unsaturation was plotted and used to determine RFs for all TG species identified from WAT. The resulting TG RFs were used to increase TG quantification accuracy.
